# Sequential and efficient neural-population coding of complex task information

**DOI:** 10.1101/801654

**Authors:** Sue Ann Koay, Adam S. Charles, Stephan Y. Thiberge, Carlos D. Brody, David W. Tank

## Abstract

Recent work has highlighted that many types of variables are represented in each neocortical area. How can these many neural representations be organized together without interference, and coherently maintained/updated through time? We recorded from large neural populations in posterior cortices as mice performed a complex, dynamic task involving multiple interrelated variables. The neural encoding implied that correlated task variables were represented by uncorrelated neural-population modes, while pairs of neurons exhibited a variety of signal correlations. This finding relates to principles of efficient coding for task-specific information, with neural-population modes as the encoding unit, and applied across posterior cortical regions and layers 2/3 and 5. Remarkably, this encoding function was multiplexed with sequential neural dynamics as well as reliably followed changes in task-variable correlations through time. We suggest that neural circuits can implement time-dependent encoding in a simple way by using random sequential dynamics as a temporal scaffold.

## Introduction

Hypothesized neocortical functions such as predictive coding (Keller and Mrsic-Flogel 2018; Rao and Ballard 1999; Bastos et al. 2012) and Bayesian inference (Helmholtz, n.d.; Ma et al. 2006) have emphasized that a crucial component of cortical computation is context: variables that indicate the external state of the world, as well as the internal state of the animal. Our work here, as well as several recent studies (Stringer, Pachitariu, Steinmetz, Carandini, et al. 2019; Steinmetz et al. 2018; Stringer, Pachitariu, Steinmetz, Reddy, et al. 2019; Minderer, Brown, and Harvey 2019; Musall et al. 2018), have indeed found that many different variables are all represented in almost every region of the dorsal cortex. These variables range from sensory and motor, to internal and cognitive. However, simultaneously representing many pieces of information in neural activity can also pose computational challenges for neural systems to overcome. We focus on two such challenges. One, how are multiple variables represented together? Two, do these representations include temporal context, e.g. progression through a time-varying behavior, an important factor for episodic memory and behavior in general? To answer these questions, we examined the structure of neural population coding during a rich yet well-controlled task, where correlated, context-dependent sensory information guided a decision-making behavior.

We recorded from large neural populations across the posterior cortex as mice performed a navigation-based visual evidence accumulation task (Pinto et al. 2018; BRAIN CoGS Collaboration, n.d.), which required subjects to comprehend time-varying relationships between multiple visual, motor, cognitive, and memory-related task variables. All these dorsal cortical areas were implicated in the mice’s performance of the task (Pinto et al. 2019), and here we wished to understand how the neurophysiology relates to behavior. Our analysis of neural data is based on conceptualizing the instantaneous population activity of neurons as a point in a high-dimensional neural state space, where each coordinate is the activity level of one neuron (Fig. 1A). The trajectory of the neural state through time is of interest from a dynamical systems perspective, and reflects how the neural circuit implements computations that may support a given behavior. Although the field has enjoyed remarkable success in describing decision-making behaviors in terms of high-level algorithms that an animal may employ, much less is known about which of many possible implementations of these algorithms are manifested in brain circuits. Here, we take a bottom-up approach of first visualizing and characterizing neural population dynamics in a decision-making behavior, which then inspired hypotheses at both computation and implementation levels (Marr and Poggio 1976).

**Figure 1.**
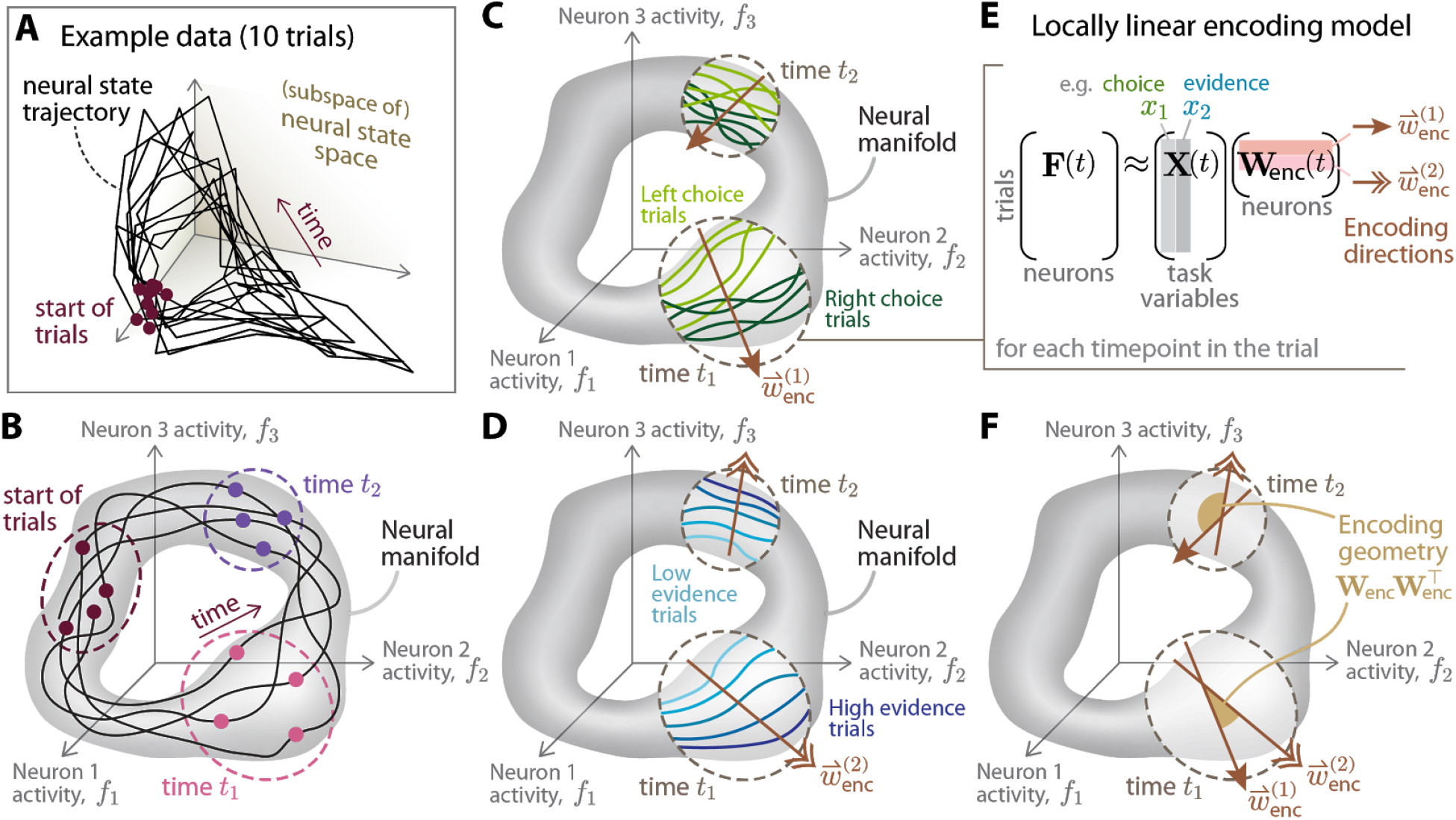
Conceptual framework for quantifying the time-dependent structure of neural-population coding of multiple task variables. **(A)** Visualization of 10 trials in one example imaging dataset (same data as in Fig. 3, 3rd column). The coordinated activity of neurons can be thought of as a point in a high-dimensional neural state space, where each axis is the activity level of one neuron. To visualize this, we linearly projected this neural state data down to 3 dimensions, as explained later for Fig. 3A. As time progresses, the neural state traces out a trajectory in the neural state space (black line, neural state evaluated at 11 time-bins in the trial and joined with a line to guide the eye). **(B)** Illustration of how neural states observed across time in the experiment appear confined to a lower-dimensional subset of the state space, termed the neural manifold (gray region). For a given timepoint in the trial, the neural states across trials (colored dots) appear furthermore confined to a local subregion of the neural manifold (dashed circles). **(C-D)** For each timepoint/local region of the manifold (dashed circles), we will show that the neural states across trials have substructure related to each of the trial-specific behavioral factors, e.g. navigational choice, illustrated in (C) and evidence levels in (D). **(E)** For a given timepoint, the substructure in (C-D) can be approximated using a linear regression model to describe how the neural state **F**(*t*) changes as task variables **X**(*t*) change. Each row of the regression weight matrix **W**_enc_(*t*) is a vector 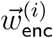 (“encoding direction”) of response strengths *across* neurons to a given task variable *x*_*i*_, and can be thought of as a gradient direction in the neural state space, e.g. arrows in (C-D). **(F)** We call the set of dot products between all pairs of encoding directions the “encoding geometry”; the dot product of two vectors is proportional to the cosine angle between the two vectors.

In many experimental scenarios, including ours (Fig. 1A), the observed neural states seemed confined to a lower-dimensional region of the neural state space, termed the “neural manifold” (Jazayeri and Afraz 2017; S. Ganguli and Sompolinsky 2012; Gallego et al. 2017; Sadtler et al. 2014; Tsodyks et al. 1999; Okun et al. 2015; Stopfer, Jayaraman, and Laurent 2003; Luczak, Barthó, and Harris 2009; Shenoy, Sahani, and Churchland 2013; Yu et al. 2009; Pang, Lansdell, and Fairhall 2016; Williamson et al. 2019; Hopfield and Tank 1986; Aksay et al. 2003). Given the large numbers of neurons that we can now record from, and the expectation that brain circuit dynamics are nonlinear, many recent efforts have focused on using nonlinear dimensionality reduction methods to describe neural manifold structure. These methods have the advantage of being able to approximate the geometrical structure of the neural manifold using a small and therefore easily examined number of coordinates. However, the often-complex transformation used to nonlinearly map neural data to low dimensions can make it difficult to interpret dimensionality-reduced observations in terms of specific candidate circuit mechanisms (see (Erem et al. 2016) for a counterexample). The alternative that we pursued was motivated by how the smoothness and continuity of physical systems often allow accurate approximations of the system’s behavior using simple (e.g. low-order polynomial) models, provided that the model is restricted to a sufficiently local region of the system’s parameter space. We show that locally-linear projections can be used to both clearly visualize and quantify intriguing structure in our neural data. In particular, we found that the neural states at a fixed timepoint in the trial, taken across trials, corresponded to local regions of the neural manifold (Fig. 1B). We thus used subsets of neural data, with each subset at a fixed timepoint in the trial, to construct locally linear approximations of how the neural state depended on various behavioral factors (Fig. 1C-D).

To understand how multiple variables were represented together, we considered “encoding directions” along which the neural state changes if the task variables change (Fig. 1E). These encoding directions can be thought of as defining a transformation of behavioral information into a neural code, which can also transform relationships stored in the neural representations. For example, using the same encoding directions for two different variables creates interference, in that these two pieces of information cannot then be distinguished, given only the neural state. However, such an encoding scheme could also support a cognitive function, generalization, by indicating that the two variables are equivalent in the process of computing successively more complex features of the world. In this way, relationships between neural representations can themselves contain extractable information about the expected structure of the world (H. B. Barlow 1989). We characterized these relationships by examining the dot products between all possible pairs of encoding directions, which we refer to as the “encoding geometry” (see Fig. 1F for a visual interpretation).

Our main finding is that at each timepoint and in all recorded posterior cortices, the encoding geometry across trials approximated the inverse of the task variable covariance matrix. This finding implies that at every timepoint, correlated task variables were encoded by approximately uncorrelated neural-population modes, which supports theories of efficient coding (Attneave 1954; Horace B. Barlow 1961). As the task variable covariances changed as a function of time in the trial, this raises a question of how a neural circuit may implement a time-dependent encoding function. Our observation that neural populations were sequentially active over the course of the trial suggests a simple neural circuit solution to the time-dependent encoding question, where the encoding geometry at a particular timepoint can be implemented via static task-input synapses onto only the subpopulation of neurons active at that timepoint. On the other hand, heterogeneous time-variations in the activities of neurons can be confounded for changes in the represented task variables, even when no such change has occurred. This brings up a second question of whether careful coordination of dynamics across the population is required for sequentially active neurons to implement encoding geometries that follow the relevant task-specific changes in time, but do *not* follow task-irrelevant temporal fluctuations of individual neurons. We show via simulations that given a sufficiently large neural population, the time-modulations of each neuron can be randomly designed, so long as they are temporally specific, and can implement an encoding geometry that varies smoothly on a timescale slower than that of individual neuron time-modulations. This is related to a mathematical property of random projections in high-dimensional spaces, which we conjecture may enable biologically plausible and robust circuit implementations where network dynamics do not have to be carefully coordinated across neurons.

Our findings have implications for longstanding theories of efficient coding. Much work on this subject has focused on how individual neurons in a population should exhibit statistically independent responses in order to represent sensory information with minimal redundancy (Rieke, Bodnar, and Bialek 1995; Laughlin 1981; Dan, Atick, and Reid 1996; Baddeley et al. 1997; Vinje and Gallant 2000; Olshausen and Field 1996; Marsat and Maler 2010; Onken et al. 2014; Weliky et al. 2003; E. P. Simoncelli and Olshausen 2001; Eero P. Simoncelli 2003), as well as how this function is modified by representational constraints and neural noise (Doi, Balcan, and Lewicki 2006; Diamantaras, Hornik, and Strintzis 1999; D. Ganguli and Simoncelli 2014, 2016; Beyeler et al. 2019; Brinkman et al. 2016; Averbeck and Lee 2006; Doi and Lewicki 2014). Our contribution is threefold. First, we examined the notion of efficient coding in terms of neural-state-level encoding directions, which capture how neural populations coordinate to represent information as a whole. Second, we discovered that efficient coding not only applies to early sensory information, but also to a set of external and internally-computed variables associated through a learned behavioral task, including in brain areas such as the retrosplenial cortex which is not traditionally considered a sensory area. Third, we report that despite dynamic task conditions and time-varying neural representations, the neural encoding geometry maintained efficient coding of task information through time. Our results thus link concepts of efficient coding with properties of computation in high-dimensional spaces, through an ethologically important question of how neocortical areas represent multiple interrelated variables to support a complex, dynamic behavior.

## Results

We performed cellular-resolution two-photon imaging of six posterior cortical regions of 11 mice trained in the Accumulating-Towers task (Fig. 2A). These mice were from transgenic lines that express the calcium-sensitive fluorescent indicator GCaMP6f in cortical excitatory neurons (Methods), and participated in previously detailed behavioral shaping (Pinto et al. 2018) and neural imaging procedures (Methods), as summarized below. Water-restricted mice were trained in a head-fixed virtual reality system (Dombeck et al. 2010) to navigate in a T-maze. As they ran down the stem of the maze, a series of transient, randomly located cues appeared along the right and left walls of the cue region corridor, followed by a delay region with no cues. Mice received a liquid reward for turning down the arm corresponding to the side with more cues, and experienced a longer time-out in the inter-trial-interval (ITI) otherwise. In agreement with previous work (Pinto et al. 2018), all mice utilized multiple cues to make decisions (Fig. 2B).

**Figure 2.**
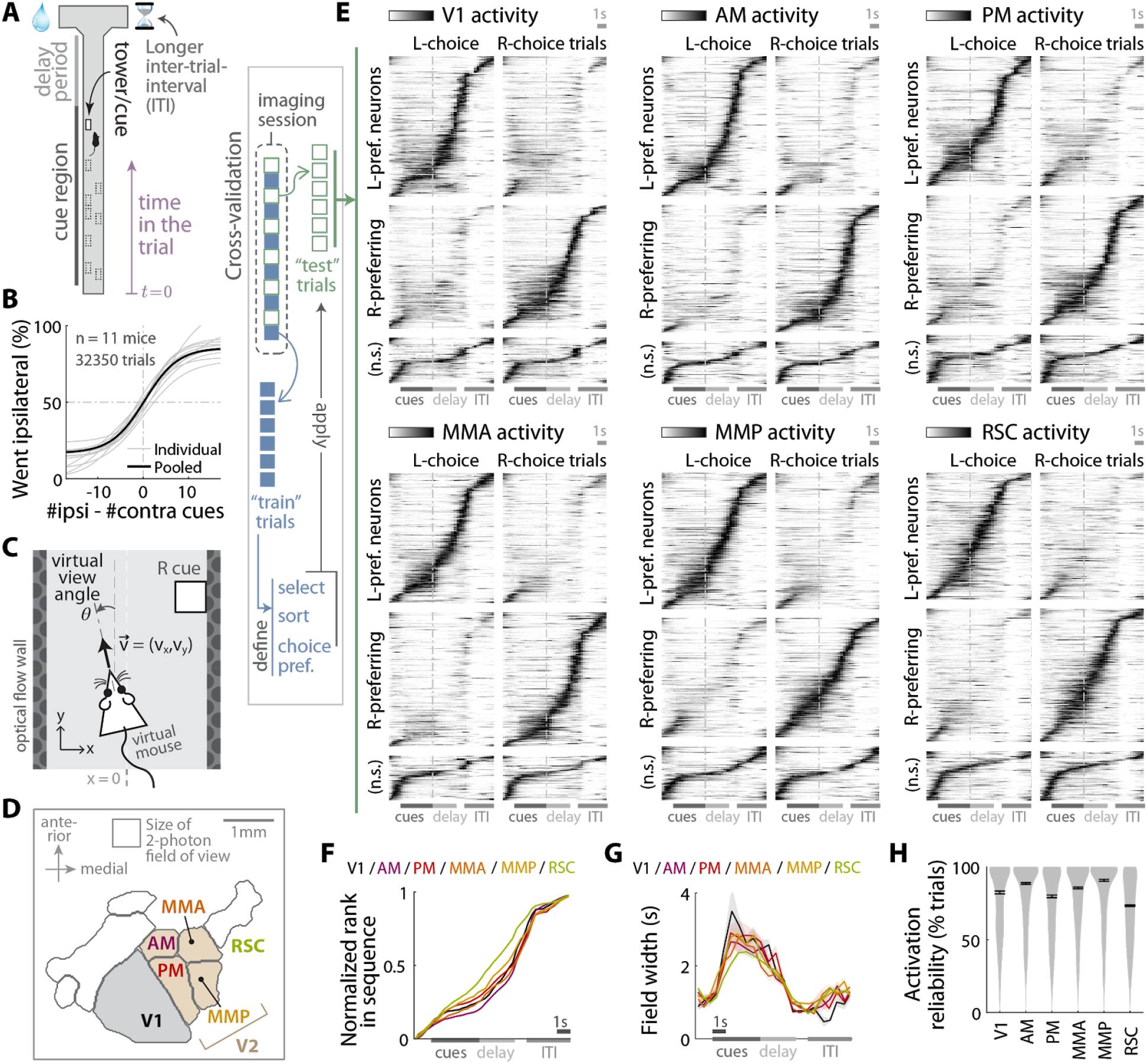
Neural populations across posterior cortex are sequentially active during the Accumulating-Towers task. **(A)** Layout of the virtual T-maze in an example left-rewarded trial. **(B)** Sigmoid curve fits to psychometric data for how frequently mice turned to the side ipsilateral to the recorded brain hemisphere, as a function of ipsilateral vs. contralateral cue counts. **(C)** Visual and motor task variables analyzed in this study. The virtual viewing angle θ determines the perspective of the virtual scene. 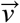 is the treadmill velocity. **(D)** Anatomical layout and labels for the six posterior cortical areas in this study. V2 refers to the set of four secondary visual areas, and RSC (retrosplenial cortex) was identified via absence of retinotopy. Visual area boundaries were functionally identified per mouse (Methods); shown here are average boundaries for *n* = 5 mice. **(E)** Normalized and trial-averaged activity of neurons, pooling data across sessions for the labeled cortical area. Neurons were divided into left-/right-choice preferring populations, and sorted by the peak activity times in correct preferred-choice trials. Each row corresponds to a single neuron, for which the left (right) column is the activity of that neuron averaged across left- (right-) choice trials. “n.s.”: neurons with no significant choice preference according to a t-test (sorted by peak activity averaged across all trials, see Methods). All sorting and normalization factors were computed using a set of training data, whereas these plots were made using the held-out set of testing data. Error trials were excluded from this analysis. **(F)** Rank (normalized to [0,1]) of sorted neurons vs. the peak activity time for that neuron. Data were pooled across sessions for a given area (colors). RSC is significantly different from other regions (*p* ≤ 10^−3^, K-S test). **(G)** Duration of activity fields vs. peak activity times. The activity field is defined as the span of time-points with activity at least half the height of the peak above baseline, in trial-averaged data. Lines: Mean across neurons, pooling data across sessions for a given area/layer. Bands: S.E.M. **(H)** Distribution (kernel density estimate) of activation reliabilities for neurons in a given area, defined as the fraction of trials in which the neuron is significantly active within its putative activity field. For (E-G), only neurons with ≥ 50% reliability were included (this criteria was not applied to any other analyses). See Fig. S1 for additional statistics. Error bars: S.E.M.

As illustrated in Fig. 2A, each trial corresponds to five conceptual (*not* explicitly cued) phases of the behavior, where the mouse traverses: a “start” region before cues appear, a spatially defined region in which cues can appear (“cue region”), the remaining length of the T-maze stem (“delay region”), a “turn region” up to the end of the trial; the last phase is the ITI. As trials could be of different durations, we resampled the behavioral and neural data according to a coordinate that measured progress through the trial (“time in the trial” which is linear w.r.t. experiment time within each behavioral phase; see Methods). In addition, we identified thirteen variables that spanned execution and psychophysics of the task: (1&2) the running tally #ipsi and #contra of cue-counts on the sides ipsilateral and contralateral to the recorded brain hemisphere; (3&4) the final tally of ipsi/contra cue counts from the previous trial; (5) the navigational choice to turn right or left; (6) the choice in the past trial; (7&8) whether the (past) trial was rewarded; (9) the virtual viewing angle θ (Fig. 2C); (10) the last value of θ in the past trial; and (11&12) treadmill velocities *v_x_* and *v_y_*; (13) *y* spatial position (along the stem) in the T-maze.

To obtain neurophysiological data, we first identified the locations of visual areas per mouse using a retinotopic visual stimulation protocol (Fig. 2D; Methods). Then, while mice performed the task, we used two-photon imaging to record from either layers 2/3 or 5 from one of six areas (Table S1): the primary visual cortex (V1), secondary visual areas (V2, including AM, PM, MMA, MMP (Zhuang et al. 2017)), or retrosplenial cortex (RSC). RSC imaging locations were chosen based on anatomy (alongside the midline) as well as the absence of retinotopic responses. After correcting for brain motion, putative single neurons were identified using a demixing and deconvolution procedure ((Pnevmatikakis et al. 2016); with modifications described in Methods). Neural activities were estimated using fluorescence-to-baseline ratios, and only neurons with ≥ 0.1 transients per trial were selected for analysis. In total, we analyzed 8,477 neurons from 143 imaging sessions. All population-level analyses were performed on datasets of simultaneously recorded neurons (average 59 neurons/session; see Fig. S1E for how this depends on cortical region and layer). All included sessions passed a behavioral performance criterion to gauge task engagement, and a small fraction of trials (0.4% overall, < 2.7% for individual sessions) were eliminated from these sessions that had extreme outlier values of task variables as these are known to cause large instabilities in fitting regression models (Methods).

### The neural state traverses an approximately time-ordered manifold in the course of a trial

Extending previous work (Harvey, Coen, and Tank 2012), we show in a cross-validated sense (Methods) that neurons in all recorded areas were sequentially active vs. place/time in the trial, and could be divided into left- vs. right-choice-preferring subpopulations (Fig. 2E; see Fig. S1 for statistics). Differences across areas were small, with RSC having more uniform tiling (Fig. 2F) and slightly more uniform field widths (Fig. 2G) of neuronal activities vs. time. Neurons were reliably active, i.e. in the majority of their preferred-choice trials (Fig. 2H), albeit a bit less so in RSC. These observations are compatible with previous findings of place/time-preferring (and choice-preferring) neurons in mouse cortex (Harvey, Coen, and Tank 2012; Morcos and Harvey 2016a; Saleem et al. 2018; Krumin et al. 2018; Runyan et al. 2017; Driscoll et al. 2017).

As individual neural activities could be ordered in time across the population, we wondered if a similar concept could be applied to the neural manifold. To visualize structure at the neural population level, we linearly projected the high-dimensional neural state data down to three dimensions, as shown in Fig. 3 for four representative imaging sessions (see Movie S1 for 3D-view animations of this figure). All visualizations were cross-validated by using half of the trials to define projection axes as explained below, and then displaying only the left-out half of trials. First, we chose a projection that reveals temporal structure in the data: given four points in the neural state space computed as the trial-average neural states evaluated at roughly equidistant timepoints across the trial, we defined the three projection axes so that they span the subspace containing these points. As seen in Fig. 3A, the trial-average neural state has a ring-like trajectory vs. time in the trial (black line). At each timepoint in the trial, the data *across* trials constitutes a cloud of points in the neural-state space, and four of these point clouds are shown in Fig. 3A including timepoints at the start of the trial, middle of the cue region, middle of the delay period, and start of the inter-trial-interval (ITI). Each of these time-specific point clouds appear in a local region of the neural state space that is distinct from the other clouds, and these clouds can be ordered along a single time coordinate in a ring-like structure (cf. the trial-average trajectory). This notion of time-specific subsets of the neural manifold being local and orderable along a time coordinate is what we call “global time order” for a manifold (illustrated in Fig. 4A), which we will quantify below. Note that due to the high dimensionality of the neural state space, global time order can be compatible with there being clear differences in neural activity patterns correlated to other behavioral factors, such as choice as already noted in Fig. 2E. Fig. 3B shows the same data in Fig. 3A but with neural states colored by the mouse’s navigational choice, from which we see that each point cloud i.e. local region of the neural manifold can have further substructure related to task variables such as choice. By using a different set of projection axes that includes a direction along which choice can best be predicted from the neural population activity (Fig. 3C), we see that a choice-related separation of neural states exists but along other directions of the neural state space that do not interfere with the temporal organization seen in Fig. 3A (see Fig. S2F for a geometrical illustration). In the last section, we explain how this kind of globally time ordered neural manifold structure can arise when neural responses to task variables have a form that factorizes from a neuron-specific time modulation function.

**Figure 3.**
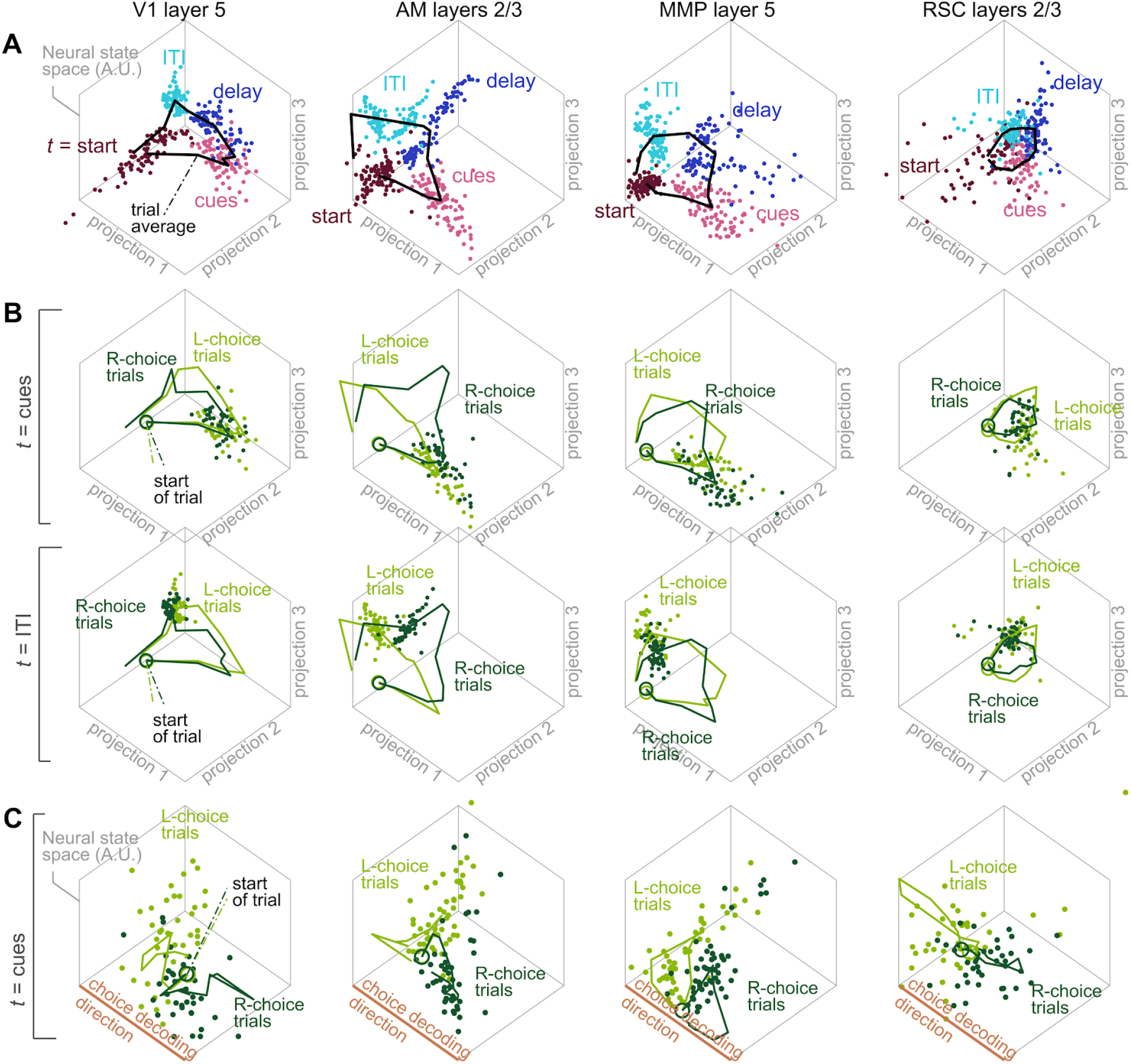
Neural states projected down to a 3-dimensional subspace (cross-validated) exhibit clear structure vs. time and navigational choice, in four typical imaging sessions. See Movie S1 for 3D view animations. **(A)** Neural states across time and trials, projected onto a 3D subspace of the neural state space as explained below. Each point corresponds to the neural state at one timepoint in one trial of a given session. Only a subset of data are shown, i.e. all correct-choice trials and including timepoints at the start of the trial (maroon), middle of the cue region(pink), middle of the delay period (blue), and beginning of the inter-trial-interval (ITI, cyan). Denoting the trial-average neural state at time *t* as *F* (*t*), the three vectors from *F* (*t_start_*) to *F* (*t_cues_*), *F* (*t_delay_*), and *F* (*t_ITI_*) define a parallelepiped in the neural state space, and the projection axes for this 3D display were chosen such that they span the subspace containing this parallelepiped (polar decomposition (Higham 1988)). Half of the trials were used to define the projection axes, and the data points in this plot include only the left-out half of trials. A standard viewing perspective (45° azimuth and 45° elevation angles) was used for all plots. Line: trial-average neural state vs. time, *F* (*t*). Columns: data from four example imaging sessions for different cortical regions, selected so as to have some active neurons at every timepoint in the trial. First, *F* (*t*) was summed across neurons in a given dataset, and then datasets were selected to have the minimum across time of this summed-activity value. **(B)** The same datasets (including identical projection axes and 3D viewing perspective) as in (A), but with data points restricted to one timepoint in the trial (rows) and colored by whether the mouse will make a left- or right-turn choice in that trial. **(C)** The same datasets as in (A), but projected onto a different 3-dimensional subspace chosen to best visualize the separation of neural states into left- vs. right-choice trials. Data points are restricted to the timepoint in the middle of the cue region (as in the first row of (B)). One projection axis is the direction along which choice can best be predicted (i.e. *decoded*) from the neural population activity (data from all timepoints were used to train a linear decoder). The two other projection axes are orthogonal to this choice-decoding direction, and chosen to maximize the variance in neural states projected onto these two axes (PCA in the hyperplane orthogonal to the choice-decoding direction). Half of the trials were used to compute all projection axes and the data points in this plot include only the left-out half of trials.

**Figure 4.**
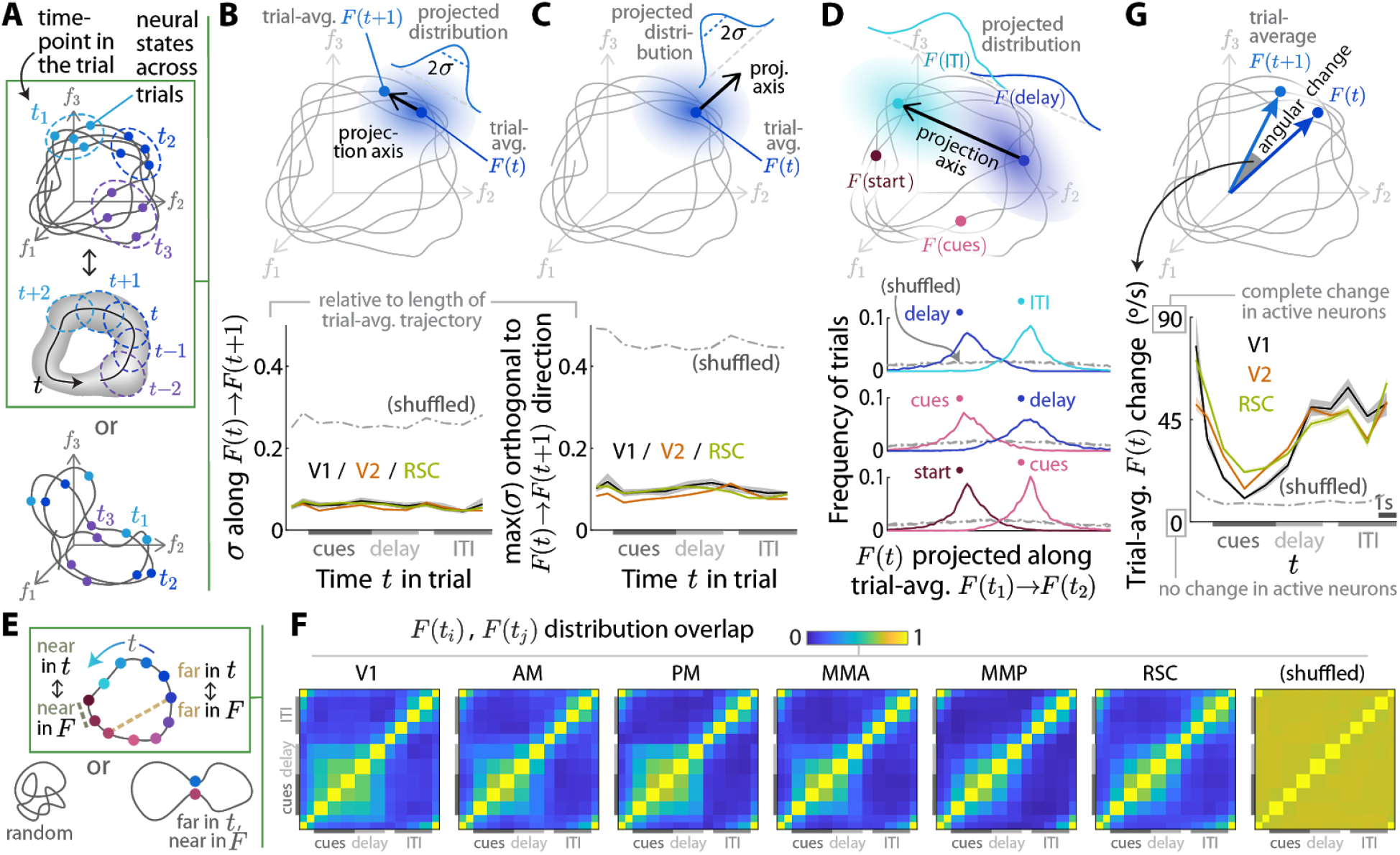
For each timepoint in the trial, the neural states across trials occupy a time-specific local region of the neural manifold. **(A)** Illustration of two possible forms of time structure for neural state trajectories across trials. Top and middle plots: a case where neural states at each timepoint in the trial occupies a different local region of the state space (dashed ellipses). This corresponds to a neural manifold where the time *t* in the trial is a coordinate along which local regions of the neural manifold can be ordered (middle plot). Bottom plot: a case where neural states at a fixed time in the trial (say, *t*_1_) occupy different locations of the neural state space depending on particulars of the trial, and these *t*_1_ neural states are interspersed with neural states at a different time in the trial (say, *t*_2_). In this case the neural state trajectory can still be confined to a low-dimensional manifold, but there is *no* one time coordinate along which local regions of this manifold can be ordered. **(B)** For each neural-state point cloud at time *t* in the trial, the normalized standard deviation of the projection of the point cloud onto the axis between the centers of two adjacent clouds. These centers are the trial-average neural states *F* (*t*) and *F* (*t* + 1), and the normalization factor is the total length of the trial-average trajectory. Projection axes were defined using one half of trials and the standard deviation computed using the left-out half of trials. Lines: average across imaging sessions. Bands: S.E.M. Shuffled: Pseudo-data with the neural state randomly cyclically permuted across imaging frames, preserving temporal and inter-neuron correlations, but destroying any relationships between neural activity and behavior (see text). **(C)** Similar to (B), but for the standard deviation of the projection of the neural state point cloud onto an axis orthogonal to the axis used in (B). This orthogonal axis was selected as the one (out of *m* − 1 possible dimensions, where *m* is the number of neurons) for which the projected standard deviation was maximal. Projection axes were defined using one half of trials and the standard deviation computed using the left-out half of trials. Lines: average across sessions. Bands: S.E.M. Shuffled: as in (B). **(D)** Distribution of neural-state point clouds for two distal timepoints, projected onto the axis between the centers of those two clouds. In the top illustration, the timepoints are at *t*_1_ in the middle of the delay period and *t*_2_ at the start of the inter-trial-interval (ITI). Projection axes were defined using one half of trials and the distribution uses the left-out half of trials. Lines: average across sessions. Bands: S.E.M. Shuffled: As in (B). **(G)** Angle between the centers of point-clouds at consecutive timepoints, divided by the time interval between timepoints. Lines: average across sessions. Bands: S.E.M. Shuffled: as in (B). **(E)** Illustration of how “time order” means that regions that are nearby (far) in time should be nearby (far) in neural-state space (box), vs. alternative possibilities (bottom diagrams). **(F)** Overlap (see Methods) between two projected distributions as in (D), for all possible pairs of timepoints. 0 means no overlap, and 1 means that the distributions are identical. Each plot was averaged across imaging sessions for the stated cortical areas. Projection axes were defined using one half of trials and overlap computed using the left-out half of trials. Shuffled: as in (B).

We devised two projection-based measures below to quantify the above concept that neural states at specific timepoints in the trial constitute (1) local regions of the neural manifold that are (2) ordered together in a ring-like structure. As a null hypothesis, we constructed pseudo-data (“shuffled” in the panels of Fig. 4) by randomly rotating the neural state across imaging frames in the session: given an *n* -by- *m* neural data matrix where *n* is the number of neurons and *m* the number of imaging frames, we randomly selected a frame *k* ∈ [1, *m*] and then constructed a pseudo-data matrix as the concatenation of columns [*k*…*m*, 1…*k* − 1] of the original matrix. This procedure preserves temporal and inter-neuron correlations while breaking relationships between the neural state and the behavioral trial structure, and exactly the same behavioral time-binning (Methods) and neural manifold structure computations were then applied to this pseudo-data as for the original data.

First, to quantify how localized the per-timepoint neural state point clouds are relative to the net displacement of these clouds vs. time in the trial, we projected the neural states onto axes related to the trial-average trajectory through time (in a cross-validated way, see Fig. 4B-C captions and Methods). Fig. 4B shows that the inter-trial spread projected along the trial-average trajectory was a small fraction of the total length of the trajectory. The maximum spread projected along axes orthogonal to this trial-average trajectory was also comparably small (Fig. 4C). These results indicate that the neural state at any one timepoint in the trial occupied a relatively small, local region of the neural manifold.

Second, two point clouds for two distal timepoints had little (cross-validated) overlap when projected along the axis between the clouds (Fig. 4D). If these clouds can be ordered in time, they can have higher overlap for nearby timepoints, but should have low overlap for any two distal timepoints (Fig. 4E). The matrix of overlap scores for all pairs of timepoints is shown in Fig. 4F. Entries near (far from) the diagonal of this matrix correspond to nearby (distal) timepoints, so we expect a time-ordered manifold to have an overlap matrix with high-valued entries close to the diagonal. Fig. 4F indeed shows such structure, albeit with less distinction between timepoints around the end of the cue period (see also Fig. 2G for how individual neurons that were active at these times tend to have the longest durations of activity). This could reflect reduced neural precision in keeping track of place/time along the stem of the T-maze away from boundaries (Tiganj et al. 2017; Singh, Tiganj, and Howard 2018), particularly since landmarks (cues) were randomly placed on every trial. Signatures of globally time-orderable structure were observed for neural manifolds in all surveyed posterior cortical regions and layers, and are far from that expected by chance of neural population activity that has the same temporal and inter-neuron correlations but no relationship to the behavioral trial structure (“shuffled” Fig. 4).

The above analyses examined how the neural manifold was geometrically structured vs. time, but for insight into circuit mechanisms it is also important to consider what this means in terms of single-neuron activity patterns. For example, if the activity levels of all neurons were to be scaled by the same amount between one timepoint and the next, this would also present as a change in state-space distance, yet there would be no change in the identities of active neurons. To quantify whether there is a change in active neurons vs. time at a neural-population level (as seen qualitatively in Fig. 2E), we examined the angular difference between the centers of the per-timepoint point clouds. As illustrated in Fig. S2A-B, an overall scaling of neural activities generates zero angular difference, whereas a 90° difference is interpretable as a complete change in active neurons given that activity levels are nonnegative. Fig. 4G shows that at all timepoints, there was an above-chance rate of change in active neurons (see Fig. S2C for all pairs of timepoints). Small differences were observed across cortical regions mostly during the cue region, where there was a V1-V2-RSC progression from more stable (near-chance) to more rapidly changing sets of active neurons.

To summarize, we found that neural manifolds across posterior cortex have ring-like geometries, where time in the trial can be used to index local subsets of these manifolds in an ordered way along the ring (Fig. 4A-F). This time-related structure is clearly visible even in the per-trial raw data (Fig. 3A-B), and arose from a systematic turnover in the active neurons vs. time in the trial (Fig. 4G). At a fixed timepoint in the trial the neural manifold can have further substructure related to behavioral factors such as choice (Fig. 3C), which we focus on in the next sections by examining how the neural state encodes multiple task variables at each timepoint. We then tie together the observations of time- and task-variable-related structure with a conceptual model in the last section.

### The neural encoding geometry approximates a whitening operation on correlated task variables

Using time in the trial to divide the neural data into local subsets of the neural manifold, we constructed locally linear models to examine how the neural population encodes multiple task variables. Specifically, we hypothesized that at a time *t* in the trial, **F**(*t*) ≈ **X**(*t*)**W**_enc_(*t*)(Fig. 1E) where the rows of **F** are neural states across trials, and each column of **X** corresponds to values of one task variable across trials. Each row of **W**_enc_ is an encoding direction for one task variable, which describes how the neural state changes if that task variable value changes (illustrated geometrically in Fig. 1C-D). Fig. S3A-B show that compared to linear encoding models, the cross-validated proportion of variance explained for neural activity is only slightly improved by including 2nd- and 3rd-order polynomial dependencies on task variables. Interestingly, even when we simulated neural responses with gaussian (i.e. nonlinear) tuning curves w.r.t. the behavioral data, these simulated neural activities could mostly be well-approximated by linear models (Fig. S3D-E). This happens because at any one timepoint in the trial, only a rather restricted range of task variable values were actually experienced by the mouse across trials in the experiment. If simulated neurons have broad tuning curves, then their task-variable dependencies can seem approximately linear within the range of the behavior; otherwise if neurons have very narrow tuning preferences to the many task variables, they tend to have near-zero signal responses and thus insufficient signal-to-noise to resolve nonlinear details of their responses. Given these empirical findings, we focused on the first-order (linear) structure of the neural code in the rest of this paper.

As explained in the Introduction, how the neural population encodes different task information *relative* to each other may have interesting implications on brain computation. We wished to quantify how the encoding directions (rows of **W**_enc_) are related to each other by examining the matrix of dot products between all pairs of encoding directions: the “encoding geometry” 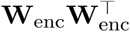 (illustrated in Fig. 1F; geometrically, the dot product between two vectors is proportional to the cosine angle between the two vectors, i.e. zero if the two vectors are orthogonal, positive if they are aligned, and negative if they are anti-aligned). However, the naıve method of first estimating **W**_enc_ and then computing 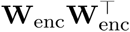 can produce noise-induced structure in the estimated encoding geometry that does not average to zero across experiments (a “noise offset”, see also (Cai et al., n.d.)). We developed a novel method for computing an estimate of the encoding geometry, which includes corrections for the noise offset as well as a procedure for selecting the form of regularization to reduce the expected variance of insofar as possible while balancing regularization-induced bias. Fig. S4B shows that in simulations, our method (1) returns an estimated encoding geometry with no significant structure when the simulated neural data is only noise; and (2) correctly interprets scenarios where choice and past-choice variables were *not* encoded (i.e. the columns and rows of corresponding to these variables are near zero) even while other, highly behaviorally correlated variables (view angle and past-view-angle) *were* encoded. Our method is thus robust against spurious, noise-induced structure across a variety of signal strength scenarios. The interested reader is referred to the Methods for details on the design of this encoding geometry estimator.

We found striking structure in the neural encoding geometry across posterior cortical areas that we could relate to the behavioral structure of task variable correlations (**C** ≡ **X**^⊤^**X** where each column of **X** corresponds to one task variable). As a high-level summary (statistical tests to follow), Fig. 5A shows that the encoding geometry averaged across time and imaging sessions has high-magnitude off-diagonal entries indicating that the encoding directions were anti-aligned (and to a lesser extent, aligned) for some pairs of task variables. Interestingly, these relationships between encoding directions tended to be *opposite* to the behavioral correlations between task variables (Fig. 5B), i.e. task variables such as view angle and choice that were positively correlated in terms of mouse behavior, tended to correspond to encoding directions that were anti-aligned (a.k.a. negatively correlated) to each other. For mathematical reasons, this observation led us to wonder if the encoding geometry may instead resemble the *inverse* of the task-variable covariance matrix (Fig. 5C). However, some rows/columns of the encoding geometry **U**, e.g. corresponding to past-trial variables and ipsilateral cue-counts, have overall reduced magnitudes (color intensity in Fig. 5A) compared to the same rows/columns of the inverse task-variable covariance matrix **C**^−1^ ≡ (**X**^⊤^**X**)^−1^ (Fig. 5C). We note that the magnitude of the encoding direction for a given task variable measures how strongly the neural state depends on that task variable, and therefore can have small magnitudes if a task variable is noisily encoded. On the other hand, the inverse task-variable covariance matrix **C**^−1^ assumes that perfect (noiseless) information about each task variable **X** is available for computing the inverse **X**^⊤^**X** of. To allow for noise in the neural code, we hypothesized that the neural encoding could be related to **X** + *ε*, where *ε* is a trial-by-variable matrix of noise fluctuations unrelated to the task variables. A noisy version of the task variable covariance matrix is **X**^⊤^**X** + **S** where **s** ≡ *ε*^⊤^ *ε* is a noise covariance matrix, and we wished to compare the neural encoding geometry to the behavioral hypothesis 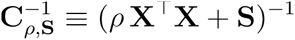 (the scalar *ρ* allows for an overall difference in scales). Since *ρ* and **S** are unknown hypothesized parameters of the brain, we empirically determined these per dataset by fitting for these parameters to produce the best agreement between **U** and 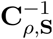 in the least-squares sense (Methods). To make this procedure tractable, we assumed that **S** is a diagonal matrix, i.e. that noise in the neural code is uncorrelated across task variables, and all fits were performed separately for each timepoint in the trial, so *ρ* and **S** depend on time. Fig. 5D shows excellent qualitative agreement between the encoding geometry and the best-fit inverse noisy task variable covariance matrix 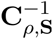, which we quantify below.

**Figure 5.**
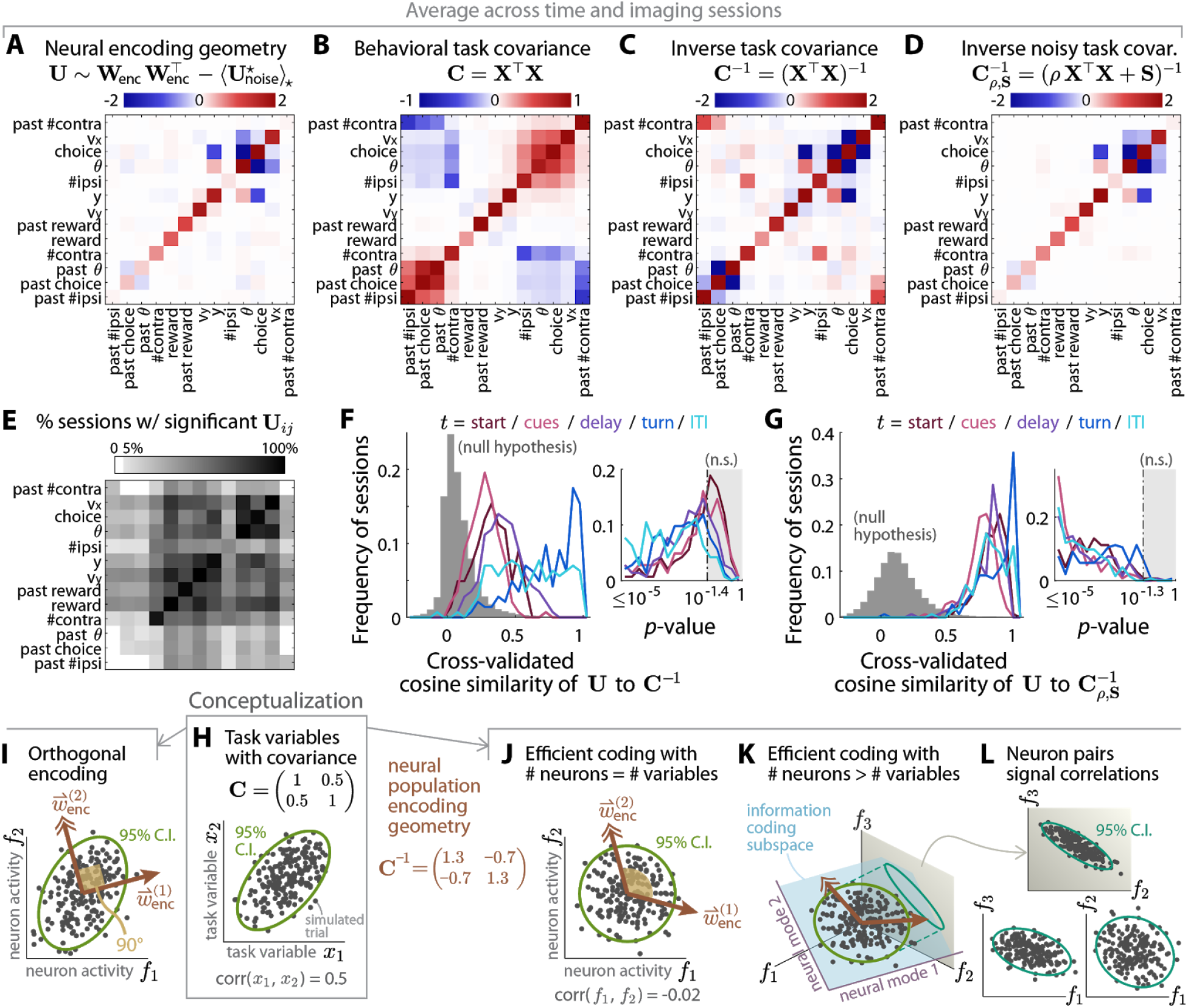
Pairwise dot products of encoding directions (“encoding geometry”) are compatible with a whitening operation, suggesting that correlated task variables are represented by uncorrelated neural modes. **(A)** Time- and session-average encoding geometry, corrected for noise-offset and regularized as explained in the text and Methods. Task variables (rows and columns) were ordered by performing hierarchical clustering on the task variable covariance matrix in (B). For timepoints where a particular task variable was not defined, e.g. the “reward” variable before the end of the trial, the row and column corresponding to that variable were zeroed out before averaging. **(B)** Time- and session-average covariance matrix of the behavioral task variable data. Task variables are in the same behaviorally-defined order as in (A), and timepoints where variables were not defined were zeroed out before averaging. **(C)** As in (B), but for the time- and session-average inverse of the task variable covariance matrix. For timepoints where only a subset of task variables were defined, the inverse covariance matrix was computed for this subset of variables, and the remaining (undefined) rows and columns were set to zero before averaging. **(D)** As in (C), but for a modified inverse task covariance matrix that accounts for different signal-to-noise levels per task variable: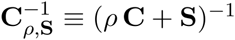. The scalar *ρ* and diagonal matrix **S** were fit per dataset so that 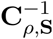 best matches the encoding geometry **U** in (A) (Methods). **(E)** Proportion of sessions for which various entries of the time-average encoding geometry matrix in (A) were significantly different from chance (*p* ≤ 0.022 after controlling for false discovery rate, see Methods). **(F)** Distribution of cross-validated similarity scores for how well the encoding geometry **U** matched the behavioral **C**^−1^. One score was computed per imaging session and timepoint in the trial, being the cosine similarity between **U**(*t*) and **C**^−1^(*t*) where entries of the two matrices were treated as two vectors. Left plot: histograms of the similarity score across imaging datasets, for various phases of the behavior (colored lines; each dataset contributes one score to each phase-specific histogram, which is the score averaged across timepoints within that behavioral phase). Right plot: *p*-value for how likely the similarity score for data is to exceed that of null hypotheses. Gray region: *p* > 0.036 not significant after controlling for false discovery rate. **(G)** Same as (F), but for the cross-validated cosine similarity between **U**(*t*) and the best-match regularized inverse 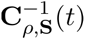 (Methods). Right plot: *p* > 0.046 not significant after controlling for false discovery rate. **(H-L)** Illustration of how different neural encoding schemes can modify signal correlations in the encoded task variables, as follows. **(H)** Simulated distribution of two correlated task variables. **(I)** Simulated responses of two neurons that linearly encode the task variables in (H), with a small amount of noise and encoding direction for *x*_1_ orthogonal to that for *x*_2_. **(J)** Simulated responses of two neurons that linearly encode the task variables in (H), with a small amount of noise and encoding geometry proportional to **C**^−1^. **(K)** As in (J), but with three neurons encoding the two task variables in (H). The neural activities lie within a 2-dimensional information-coding subspace (blue plane) spanned by the encoding directions (brown arrows), and the neural modes that define this subspace are uncorrelated (95% C.I. is a circle). **(L)** The same simulated data in (K), but plotted for various pairs of neural axes. These pairs of neurons have nonzero signal correlations (95% C.I. are ellipses).

We evaluated the significance of the above findings w.r.t. a null hypothesis that breaks *only* potential relationships between neural activity and behavior, while preserving the same time-dependent and temporally correlated neural activity distributions, inter-neuron covariance, and task-variable covariance as the actual experiment. Our procedure was inspired by (Elsayed and Cunningham 2017), but designed to preserve all the above data features. As detailed in the Methods and illustrated in Fig. S5A, our procedure takes as input the neural data **F**(*t*), which has a 3-dimensional structure where rows correspond to trials, columns correspond to neurons, and the third dimension corresponds to time within the trial *t*. To generate one instance of the pseudo-data null hypothesis **F**_∗_(*t*), we first randomly permuted the neuron-by-time “slabs” of **F**(*t*) across trials (rows). This trial permutation breaks correlations between the neural and behavioral data, while preserving neural correlations across the two other dimensions that define each slab, i.e. inter-neuron and within-trial temporal correlations. However, trial permutations removes long-timescale correlations across trials, i.e. across rows of **F**(*t*) assuming that they are ordered by time in the session. Inter-trial correlations can be signal correlations reflecting neural responses to the trial-specific behavioral conditions, which we indeed want the null hypothesis to break, but alternatively they could have been induced by slow, task-unspecific modulations of the neural state. We wish to retain the latter type of temporal structure in the null hypothesis, which we reasoned is not due to inter-trial correlation per se but rather structure in the *auto*correlation function. (The autocorrelation specifies how similar the neural state is between temporally adjacent trials, irrespective of i.e. averaged over the behavior-specific trial index.) We therefore in a second step constrained the row-autocorrelation function of **F**_∗_(*t*) to approximate that of **F**(*t*) by applying a convolution operation to the rows of **F**_∗_(*t*), or equivalently adding to each row a weighted sum of a small number of adjacent rows in order to impose similarities between the neural state in adjacent trials. Lastly to avoid sampling discreteness due to the limited number of trials per experiment, we then added a bit of noise to the entries of **F**_∗_(*t*) in a way that preserved the mean activity levels, inter-neuron covariance, and temporal correlations (Methods). In exactly the same way as we did for the data, **F**_∗_(*t*) was used for computing the null hypothesis encoding geometry w.r.t. the unmodified task variables **X**(*t*). By generating the pseudo-data **F**_∗_(*t*) from a time-specific data matrix **F**(*t*), this null hypothesis tests whether there is significant task-variable-related structure in the encoding geometry beyond trial-time dependence in neural activity levels, inter-neuron correlations and a trial history effect (autocorrelation). Since task variable responses could have produced all of these data features that were preserved in the null hypothesis, this is to the best of our ability the most conservative null hypothesis we can define.

For most pairs of task variables, the corresponding entry of the encoding geometry matrix was significantly different from the null hypothesis in a substantial fraction of imaging sessions (Fig. 5E, controlled for false discovery rate, see Methods). Given that the encoding geometry had structure beyond that expected from chance, we then proceeded to ask how well this neural structure matched either the noiseless inverse task covariance matrix **C**^−1^, or the noisy alternative 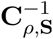. For cross-validation (details in Methods), we first randomly divided each dataset into two halves of trials, and optimized the free parameters *ρ* and **S** using one half of trials. Then keeping the free parameters fixed, we computed the cosine similarity between **U** and 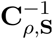 (or **C**^−1^ which has no free parameters) using only data from the left-out half of trials. The cosine similarity score is defined by converting each matrix to a vector of its entries and then computing the cosine angle between the two vectorized matrices. This score is insensitive to an overall scaling of either matrix, and can range from +1 if the matrices are identical (up to a scale), to −1 if their entries have opposite signs. Fig. 5F shows that although **U** can resemble **C**^−1^ for some datasets and at some timepoints in the trial, there is a much better correspondence between **U** and 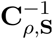 that is highly consistent across time and imaging sessions (Fig. 5E; this also holds for the Euclidean distance between **U** and 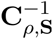, Fig. S5B).

The very high significance of the above comparisons w.r.t. the null hypothesis show that the relationship of the encoding geometry **U** to behavioral features is far beyond that expected from chance-level correlations of task variables with temporally structured, low-dimensional neural population activity. We additionally asked if the hypothesized relationship of **U** to 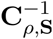 can be explained as arising from neurons that did respond to task variables, but in a random manner e.g. random mixed selectivity (Kennerley et al. 2009; Fusi, Miller, and Rigotti 2016; Rigotti et al. 2013; Raposo, Kaufman, and Churchland 2014). A random neural code corresponds to an encoding weight matrix **W**_enc_ with entries that are randomly drawn from some symmetric distribution, or equivalently encoding directions that are a set of randomly oriented vectors in the neural state space. In high dimensions (many neurons) random vectors are likely to be orthogonal to each other, i.e. the off-diagonal entries of the encoding geometry (dot products between different encoding directions) are likely to be near zero. However as seen qualitatively in Fig. 5A, the encoding geometry in our data has off-diagonal entries that are far from zero, and Fig. S5D-left shows that the cosine similarity of **U** to 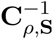 in Fig. 5E has substantial positive contributions from just the off-diagonal entries of the two matrices. This is qualitatively different than in simulations of random linear encoding of the behavioral data (for the same neural population sizes as the neural data), where off-diagonal contributions are close to and mostly symmetrically distributed around zero (Fig. S5D-right). In sum, the similarity of the encoding geometry **U** to the inverse noisy task variable covariance hypothesis 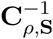 is clearly beyond that explainable by either chance or random linear codes (see also Fig. S4C-F for how this structure does not match some *non*linear codes that we may hypothesize).

The above observed structure of the encoding geometry **U** has an interpretation that relates to theories of efficient coding. We derive this mathematically in the Methods and sketch the idea here, starting from the conceptually simpler case of being proportional to the inverse (non-noisy) task variable covariance matrix **C**^−1^ ≡ (**X**^⊤^**X**)^−1^. Fig. 5H illustrates the joint distribution of two correlated task variables *x*_1_ and *x*_2_ in a simulated dataset, which we assume are linearly encoded by some neural population: **F** = **XW**_enc_. If this population consists of many neurons that encode each task variable with randomly distributed response weights, the encoding directions 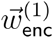 and 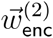 are likely to be orthogonal due to a mathematical property of high-dimensional spaces. In this case, we can think about the neural code as a veridical copy of the behavioral data (i.e. preserving correlations), where the behavioral *x*_1_ and *x*_2_ axes in Fig. 5H are mapped respectively onto the neural axes 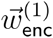 and 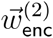 in Fig. 5I. If instead the encoding geometry is 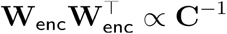, then given exactly as many neurons as encoded task variables we can design encoding directions to be 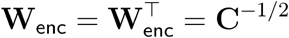, which results in the activities of the two neurons being uncorrelated as illustrated in Fig. 5J. This operation of encoding statistically *dependent* information with statistically *independent* channels (neurons) is called “whitening”, and (barring other constraints such as correlated noise) is optimal in the sense that it omits redundancy in the neural code (E. P. Simoncelli and Olshausen 2001). However, if there are more neurons than encoded variables, there is freedom in how the information-coding subspace spanned by encoding directions can be embedded in the neural state space (Fig. 5K). Even though the neural modes are mutually uncorrelated, pairs of individual neurons can still have a variety of nonzero signal correlations depending on the orientation of the information-coding subspace (Fig. 5L), and we indeed found nonzero correlations between pairs of neurons in our data (Fig. S5E-G). Our findings thus deviate from previous reports of efficient coding in two interesting ways: (1) whitening is only evident at the level of neural-population modes, and not pairs of neurons; (2) the whitening operation is imperfect, in that instead **U** ∝ **C**^−1^ of we observed 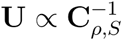, the inverse of a noise-corrupted behavioral covariance matrix that has different signal-to-noise levels for each task variable.

### The neural encoding geometry follows changes in inverse task-variable covariances vs. time in the trial

So far we have performed all analyses independently per timepoint in the trial, but which findings are actually time-dependent and which are not? First, the scale of some task variables depended on time in the trial, and the corresponding encoding weights of neurons across the population *inversely* followed this scale (Fig. S6A-C). This is as expected of the hypothesis that the neural encoding approximates a whitening operation at all timepoints, since whitening requires equalization of the scales of all encoded variables. Second, correlations between task variables changed slowly as a function of time in the trial (Fig. 6B), and the encoding geometry **U**(*t*) tracked the corresponding time-variations in the inverse task-variable covariance matrix 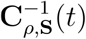 well (Fig. 6A). The similarity of **U**(*t*) to 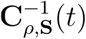 was comparably high for all timepoints *t*, as previously quantified in Fig. 5G.

**Figure 6.**
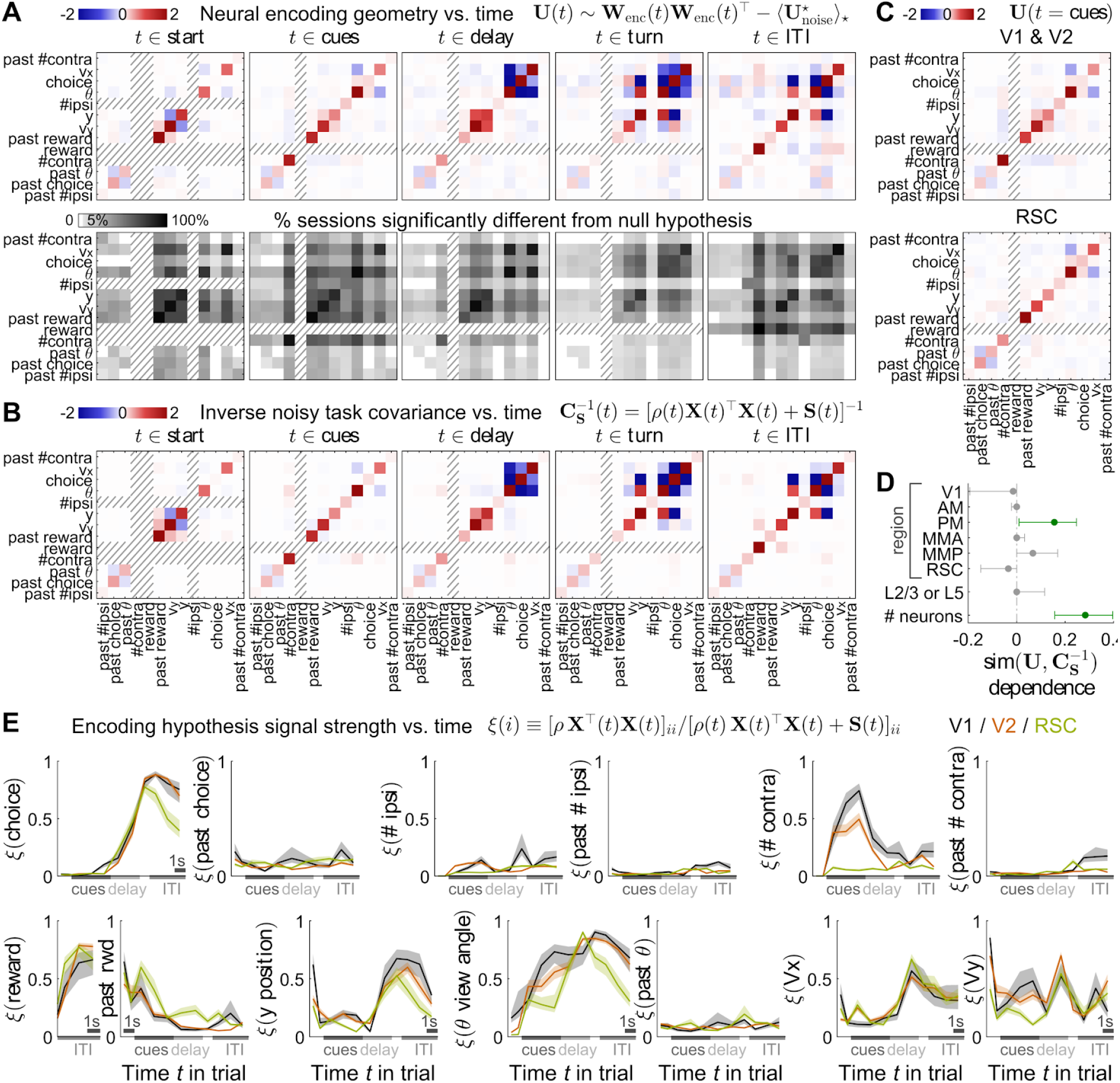
Encoding geometry follows time-variations in the inverse task-variable covariance matrix, with inter-region differences explainable as differences in task-variable-specific signal strength. **(A)** Session-average encoding geometry as a function of time in the trial (columns). Top row: Noise-corrected and regularized encoding geometry estimate as in Fig. 5A, but each plot was only averaged across a subset of timepoints within the stated behavioral phase of the trial. Bottom row: Analogous to Fig. 5E, proportion of sessions for which various entries of the encoding geometry in the corresponding top plot are significantly different from chance (*p* ≤ 0.011 after controlling for false discovery rate). Hatched rows/columns: task variable not defined in this behavioral phase. **(B)** Inverse noisy task variable covariance matrix, with signal strength *ρ*(*t*) and noise parameters **S**(*t*) fitted to the estimated encoding geometry for each timepoint and recording session as in Fig. 5D. **(C)** Encoding geometry during the cue period, averaged across visual area datasets for the top plot, vs. average across RSC datasets in the bottom plot. The reward variable (hatched row and column) is not defined in the cue period. **(D)** Coefficients from an L1-regularized linear regression model for how the similarity of (A) to (B) depended on factors like cortical region/layer and number of recorded neurons. Each imaging session contributed one data point to this model, the predicted variable being the time-average cross-validated cosine similarity of the encoding geometry to the inverse noisy task covariance hypothesis (as histogrammed in Fig. 5G). For each session, cortical region regressors were defined as indicator variables that were 1 if and only if that session was recorded from the stated region, and the layer regressor was 1 for recordings from layer 5 and 0 for layers 2/3. Error bars: 95% C.I. of 1000 bootstrap experiments. Significant factors are indicated in green. **(E)** Estimated proportion of signal variance ξ(*i*) for each task variable *i* in the best-fit inverse noisy task covariance matrix (B). Since **X**^⊤^**X** is the task variable covariance matrix, the *i* ^th^ diagonal entry (*ρ***X**^⊤^**X**)_*ii*_ can be thought of as the signal variance for task variable *i*, which is plotted here as a function of time and relative to the total variance (*ρ***X**^⊤^**X** + **S**)_*ii*_ for that variable. Lines: average across sessions. Bands: S.E.M.

In addition to the above time-dependence, differences in the encoding geometry across cortical regions and/or layers may hint at anatomical trends in cortical processing. For example during the cue period, the encoding geometry averaged across V1 and V2 datasets (Fig. 6C-top) exhibits some small qualitative differences with respect to the encoding geometry averaged across RSC datasets (Fig. 6C-bottom). Despite these differences in the *structure* of the encoding geometry **U**(*t*), we wondered if our hypothesis about the *function* of **U**(*t*), i.e. that it resembles the inverse of a noisy task-variable covariance 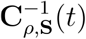, holds across cortical regions. To evaluate this, we used linear regression to predict the cosine similarity of **U**(*t*) to 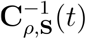, with covariates being the set of indicator variables (values 0 or 1) for whether a given dataset was recorded in a particular cortical region or layer (e.g. a dataset recorded from layer 5 of V1 will have regressor values “V1”=1, “layer”=1, and value 0 for all other regressors). We additionally included the number of simultaneously recorded neurons in the datasets as a covariate, since per-neuron noise can affect the accuracy to which we can estimate the encoding geometry (see Fig. S1E for how neuron count depends on cortical region and layers). Fig. 6D shows that **U**(*t*) is significantly more similar to 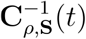 in recordings of larger neural populations, and in fact the number of neurons is the largest factor in predicting the goodness of the 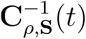 hypothesis fit. Beyond this population size effect, the similarity of **U**(*t*) to 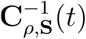 is comparable across cortical regions, with slightly better agreement in the secondary visual area PM, and no significant effect of cortical layer.

We can understand both the time dependence and inter-region differences in encoding geometries **U**(*t*) by examining the best-fit parameters *ρ*(*t*) and **S**(*t*) in the hypothesized structure 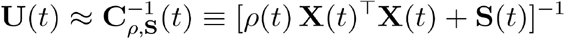. As explained in the previous section, we can think of *ρ***X**^⊤^**X** as a task-signal covariance and as a noise covariance that determines the neural code. To visualize these parameters, we defined a measure of “signal strength” ξ(*i*) for task variable *i* as the proportion of signal over total variance (diagonal entries of the covariance matrices) for that variable: ξ(*i*; *t*) ≡ [*ρ*(*t*)**X**(*t*)^⊤^**X**(*t*)]_*ii*_/[*ρ*(*t*)**X**(*t*)^⊤^**X**(*t*) + **S**(*t*)]_*ii*_. This signal strength is plotted in Fig. 6E for each task variable, and exhibits variable-specific time dependencies as well as some inter-region differences. The largest differences are for visual-related task variables such as the running tally of contralateral cues and the virtual viewing angle, for which the signal strengths are highest in V1 and progressively lower in V2 and RSC. The neural coding of past-trial reward is also significantly higher and persists for a longer duration of the trial in RSC compared to the visual areas. In sum, posterior cortical regions are all comparably well-modeled as having encoding geometries **U**(*t*) that imply a time-specific whitening of noisy task variable information 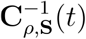, but the signal-to-noise parameters of the 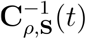 hypothesis varies with time as well as across cortical regions for some task variables.

### Neural responses are parsimoniously explainable by a multiplicative time-dependence model

How can a neural circuit implement an encoding geometry that adjusts at every timepoint in the trial (Fig. 6A), so as to cancel out the different structure of behavioral correlations at different timepoints (Fig. 6B)? While recurrently connected neural networks are theoretically powerful enough dynamical systems to produce such dynamic encoding functions, we point out a particularly simple and robust candidate implementation inspired by how the encoding of task variables occurred in conjunction with sequential dynamics across the neural population (Fig. 2E, Fig. 4G). Here, we first demonstrate that neural sequences in posterior cortex can be parsimoniously described as having time-modulations of neural activity levels that approximately factorizes from their task-variable dependencies. As hypothesized in the next (theory) section, this factorization allows time-specific encoding geometries to be stored in static task-input synapses, and moreover in a robust way that does *not* require careful coordination with neural dynamics across the network.

In the previous sections, we fitted encoding models separately to the data for each timepoint *t* in the trial. This means that the activity of neuron *i* in trial *j* was hypothesized to have the form 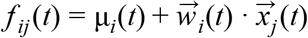, where μ_*i*_(*t*) is the neuron’s trial-average activity, 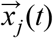 are task variable values in trial *j*, and the encoding weights 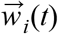 can potentially be different at every timepoint *t*. We alternatively proposed a multiplicative time-modulation model, where the neuron’s activity has the form 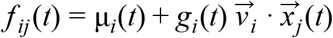. In the latter model, the time-dependence of a neuron’s activity is completely described by functions that do not depend on task conditions: a trial-average baseline μ_*i*_ (*t*), and a neuron-specific time-modulation function *g_i_*(*t*) that applies to all trials. The task-variable dependence of such a neuron’s activity arises from the scaling of *g_i_*(*t*) by 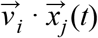, where 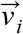 are time-independent encoding weights for the task variables. As opposed to the per-timepoint encoding model which has (13 task variables)⨉(11 timepoints)=143 free parameters, the multiplicative time-modulation model has only 13+11=24 free parameters. Nevertheless, the cross-validated variance explained in Fig. 7A shows that the much simpler multiplicative time-modulation model predicted all single-neuron activities almost as well as the per-timepoint encoding models. Furthermore, an even simpler model with constant *g_i_*(*t*) could *not* model neural activities well: neural responses varied more in time than could be explained by dependencies on time-varying task variables on top of a time-varying baseline (Fig. 7B). In fact, equivalent to the *g_i_*(*t*) functions of most neurons being near-zero (inactive) for at least some portion of the trial, we previously showed in Fig. 4G and Fig. S2C that the subset of active neurons changed substantially across time according to a neural-population-level measure. The multiplicative time-modulation model thus provides a parsimonious description of single-neuron activities, and we refer to this pattern of activity that we observed across posterior cortex as “multiplicative neural sequences”.

**Figure 7.**
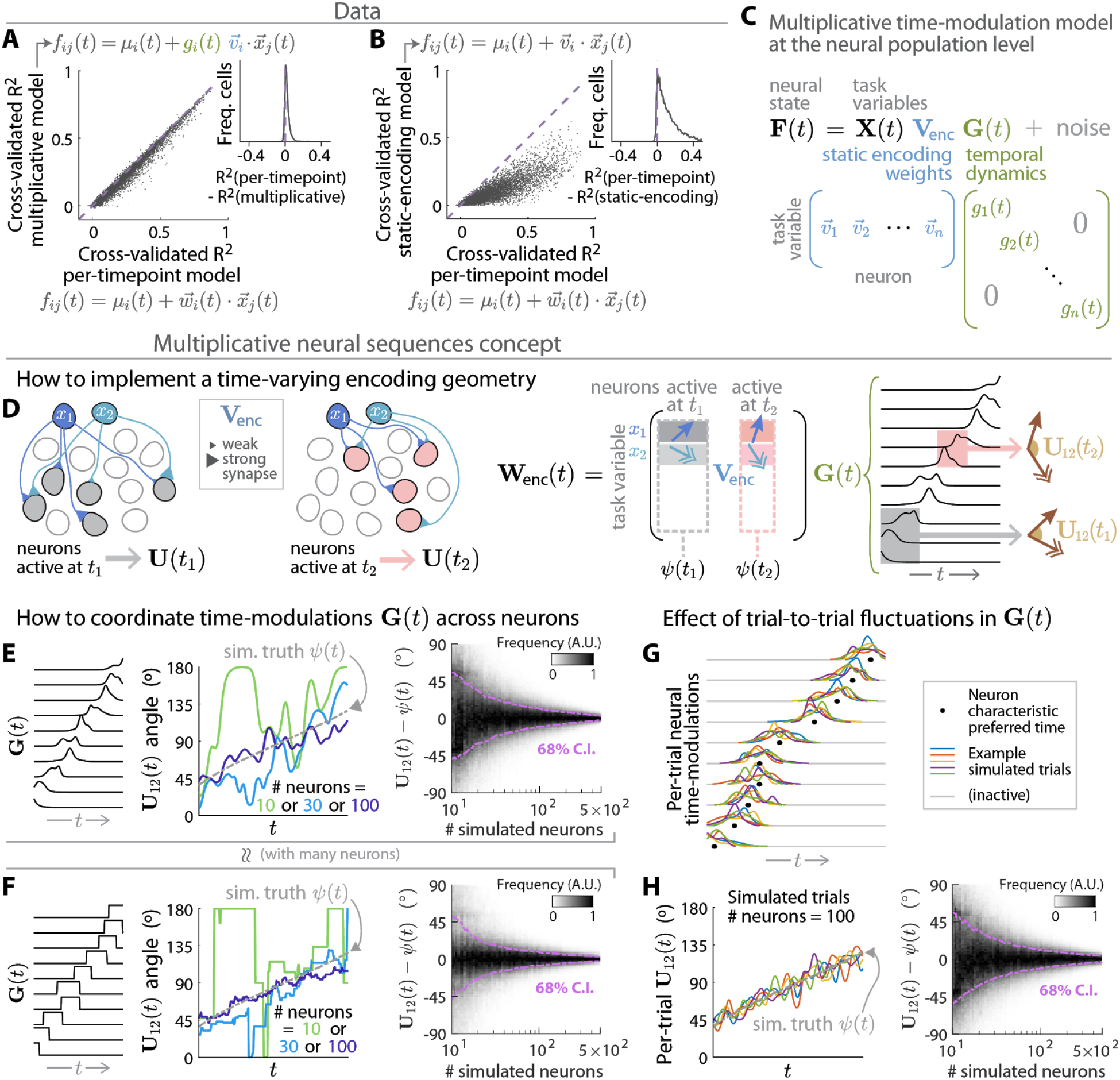
Neural activity is well explained by a multiplicative sequential encoding model, which suggests a simple implementation of time-specific encoding geometries by a neural population with static task-variable encoding weights. **(A)** Cross-validated variance explained for a multiplicative time-dependence model vs. a fully flexible model with task-variable encoding weights that can be arbitrarily different per timepoint (ridge regression per neuron, see Methods). *f_ij_*(*t*) is the activity of neuron *i* vs. time *t* in trial *j*, and μ*i*(*t*) is the empirical mean of *f_ij_*(*t*) across trials. In the multiplicative time-dependence model *g_i_*(*t*) is a 11-parameter time-dependence function and 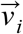 is a vector of constant encoding weights for the 13 task variables, totaling 24 free parameters. In the per-timepoint model 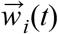 has 13 × 11 = 143 free parameters. Each point in the plot corresponds to a single neuron. Inset: distribution across neurons of differences in variances explained for the two models (*x* minus *y* coordinates of the left scatter plot). **(B)** Same as (A), but for a model with constant encoding weights per neuron (13 free parameters) vs. the per-timepoint encoding model. **(C)** Correspondence between the single-neuron multiplicative time-modulation model parameters in (A) and the notation for the same model at the neural-population level. Each column of the time-independent encoding matrix **V**_enc_ corresponds to the encoding weights 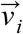 of one neuron *i*, and the time-modulation matrix **G**(*t*) is diagonal with each entry being the activation level *g_i_*(*t*) of neuron *i* at time *t*. **(D)** Conceptualization of how a multiplicative neural sequence (C) can have an encoding geometry that changes with time in the trial. At a given time *t*, the encoding directions are the rows of the matrix **W**_enc_(*t*) = **V**_enc_ **G**(*t*). For a sequentially active neural population, **W**_enc_(*t*) has contributions from only those columns of corresponding to neurons that are active at time *t* (dashed rectangles). Thus, columns **V**_enc_ of corresponding to different subsets of neurons active in (say) times *t*_1_ vs. *t*_2_ can produce different encoding geometries **U**(*t*_1_) ≡ **W**_enc_(*t*_1_)**W**_enc_(*t*_1_)^⊤^ vs. **U**(*t*_2_). **(E)** Middle plot: angle between two encoding directions, in three example simulations where a population of *n* neurons responds to two task variables according to the multiplicative sequential encoding model in (C). **G**(*t*) functions were randomly generated such that neurons each have a preferred activation time that is uniformly distributed across the population (Methods), thus forming a sequence as illustrated in the left plot. The 2 × *n* matrix **V**_enc_ was constructed so that the columns of **V**_enc_ corresponding to neurons with maximal **G**(*t*) at time *t* have rows that form an angle ψ(*t*) (dash-dotted line). The angle between the two encoding directions (colored lines) have fluctuations away from ψ(*t*) due to the randomly simulated **G**(*t*). Right plot: distribution across simulation experiments and time of differences in the angle between encoding directions vs. ψ(*t*), as a function of neural population sizes in the simulations. **(F)** Same as (E), but the time-modulations **G**(*t*) in these simulations were converted to binary functions via thresholding, i.e. each neuron is only either “on” or “off”. **(G)** Simulated time-modulation functions **G**(*t*) for 10 (out of a population of 200) sequentially active neurons, each of which had time-modulations that were localized around a characteristic time preference for that cell (dots), but otherwise randomly drawn per trial (colored lines; shown as gray when the neuron is inactive). **(H)** Left plot: angle between encoding directions vs. time for the 5 simulated trials in (G), where in each trial *k* the neural time-modulations **G**^(*k*)^(*t*) multiply the *same* time- (and trial-) independent encoding weight matrix **V**_enc_. **V**_enc_ was constructed so that the columns of **V**_enc_ corresponding to neurons with time preference 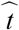 have rows that form an angle 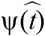 as shown in the plot (dash-dotted line). Note that unlike (E), **V**_enc_ depends only on the generative time preference of neurons, and not the trial-specific instance of **G**^(*k*)^(*t*). Right plot: same as (E-right), but for these simulations where **V**_enc_ depends only on the generative time preference of neurons.

### Multiplicative neural sequences as a robust design for neural implementations of time-specific encoding geometries

From a neural circuit design standpoint, multiplicative neural sequences have two interesting theoretical properties that we can show via simulations and mathematically explain. One, because different timepoints in the trial correspond to different subsets of active neurons, *time-specific* encoding geometries can be implemented via *neuron-specific* (but time-independent) task-input weights. Two, given a sufficiently large neural population, multiplicative time-modulations act essentially like binary on/off functions that determine the subset of active neurons, i.e. details of individual neural activation levels have net little effect. We argue that these two properties make for a biologically plausible design where task- and time-specific encoding geometries can be learned with minimal requirements on inter-neuron coordination and little sensitivity to the exact form of neural dynamics across the population. We demonstrate this quantitatively by describing how to design encoding geometries that vary systematically with time in the trial (as seen in our data Fig. 6A,D), using the above two properties of multiplicative neural sequences.

First to recap the multiplicative time-modulation model at the neural population level (Fig. 7C), the neural state **F**(*t*) at time *t* in the trial arises from responses to task variables: **X**(*t*)**V**_enc_ **G**(*t*). Here **X**(*t*) is a trial-by-variable matrix of task variable values **V**_enc_, is a variable-by-neuron matrix of static encoding weights, and the time-modulation matrix **G**(*t*) is diagonal with entries being the activation level of each neuron. We explore circuit design in this context by specifying how to generate **V**_enc_ (interpretable as strengths of task-input synapses, Fig. 7D-left) as well as **G**(*t*) (e.g. sequential activations illustrated in Fig. 7D-right), so as to produce encoding geometries with various properties discussed below.

Assuming that each neuron is only active at a few timepoints in the trial, how can we design the time-*independent* **V**_enc_ to produce a time-*dependent* encoding geometry? Because the encoding directions at time *t* are given by **W**_enc_(*t*) = **V**_enc_ **G**(*t*), inactive neurons (zeros in **G**(*t*)) are zeroed out in **W**_enc_(*t*) and thus do not contribute to the population-level encoding geometry **U**(*t*) ≡ **W**_enc_(*t*)**W**_enc_(*t*)^⊤^. As illustrated in Fig. 7D, we can thus use the subset of columns of **V**_enc_ corresponding to only active neurons to specify the desired **U**(*t*). If two timepoints *t*_1_ and *t*_2_ have non-overlapping subsets of active neurons, then the encoding geometries at these two timepoints can be completely different because they are determined by two disjoint subsets of columns of **V**_enc_. In the extreme (but pedagogically illustrative) case where each neuron is active at exactly one timepoint in the trial, every timepoint *t* corresponds to a unique subset of columns of **V**_enc_ that can be freely chosen to produce any desired encoding geometry at time *t*. In the more realistic case where neurons have some range of time in which they are active, the duration of their activities constrain how quickly the encoding geometry can change vs. time. In this way, the continuously changing subsets of active neurons in a multiplicative neural sequence (cf. neural state change measures in Fig. 4G,F) enables different encoding geometries vs. time to be implemented via different columns (neurons) of the static “synaptic weights” matrix **V**_enc_.

Although the encoding matrix **V**_enc_ has no explicit time dependence, in the above design **V**_enc_ depends implicitly on the time-modulation functions **G**(*t*) through which columns of **V**_enc_ contribute at which times. How exactly should the designs of **V**_enc_ and **G**(*t*) be coordinated to produce a particular encoding geometry **U**(*t*)? Mathematically, because the encoding geometry is 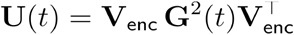, we may in principle need to (a) adjust the entries of **V**_enc_ to the specific neural activation levels in **G**(*t*); and/or to (b) coordinate the time-modulations across neurons (entries of **G**(*t*)). Conceptually, the influence of **G**(*t*) reflects how if neural activity levels constitute information in the brain about particular task variables, then any unrelated time-modulations of these activities can be confused as changes in task variable values even when there was no such change. We might thus wonder if careful tuning of the circuit is required to coordinate encoding and neural dynamics in a way that ensures reliable representation of task information through time. Remarkably, we can show via simulations that for multiplicative neural sequences, precise coordination is *not* required (i.e. neither (a) nor (b)). Instead, random time-modulations across neurons will average out in a large enough neural population, enabling the encoding geometry to vary smoothly according to the structure of **V**_enc_ on timescales longer than that of individual neural time-modulations.

Per the definition of multiplicative neural sequences, we simulated neural states of the form **X**(*t*)**V**_enc_ **G**(*t*) in response to two randomly generated task variables **X**(*t*). As illustrated in Fig. 7D-right (see Methods for details), we first generated **G**(*t*) functions such that each neuron had randomly shaped time-modulations roughly confined to a preferred time window (also randomly determined per neuron), which then formed a sequential activation pattern across the population. We then constructed **V**_enc_ so that the subset of its columns corresponding to neurons with maximal **G**(*t*) at time *t* had an angle ψ(*t*) between their rows (encoding directions). Fig. 7E-middle shows the encoding geometry in three example simulation experiments with small to large neural population sizes. Here, to make comparisons across neural populations of different sizes more intuitive, we measure the encoding geometry in terms of the *angle* between encoding directions (the unnormalized dot product depends on the total number of neurons). Since the **G**(*t*) were *not* coordinated across neurons, the encoding geometry randomly deviated from ψ(*t*) across time (difference between solid colored and dashed gray lines in Fig. 7E-middle). However, these deviations became progressively smaller for simulations with larger neural-population sizes, and with a few hundred neurons different simulations i.e. different instantiations of **G**(*t*) all produced encoding geometries that closely approximated ψ(*t*) (Fig. 7E-right). The convergence of **U**(*t*) to ψ(*t*) at large population sizes is insensitive to details of **G**(*t*) such as the shapes of time-modulation functions. For example, Fig. 7F shows qualitatively similar results when the simulated **G**(*t*) were converted to binary on/off functions via thresholding.

The above random design for **G**(*t*) has a limitation and an extension that we now discuss, before lastly providing a theoretical explanation for why it works. First, without fine-tuning **V**_enc_ to exploit the specific neural timecourses in a given **G**(*t*), the timescale of single-neuron activities limits how quickly the encoding geometry **U**(*t*) can track changes in ψ(*t*). For the simulations in Fig. S7A-F, we designed ψ(*t*) to be constant except for two abrupt switches between low and high values, and used ψ(*t*) to define **V**_enc_ in the same way as before. If we then generated **G**(*t*) such that each neuron was active for longer (shorter) periods, **U**(*t*) transitioned slowly (quickly) vs. time between the low and high values of ψ(*t*) (Fig. S7A vs. Fig. S7D). The ability of populations of fast-timescale neurons to implement fast changes in ψ(*t*) trades off with a need for larger population sizes to average out random deviations from ψ(*t*) (Fig. S7B-C vs. Fig. S7E-F). Within these limitations, that the designs of **V**_enc_ and **G**(*t*) do not need to be carefully coordinated has an interesting extension to how the encoding geometry can be robust to variability in **G**(*t*), e.g. trial-to-trial stochasticity present in our data and often reported by others. Fig. 7G illustrates five simulated trials *k* with different generated **G**^(*k*)^(*t*), but where each neuron *i* has a fixed, characteristic preferred time 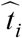 (black dots) around which they are active. We then designed a *single* **V**_enc_ with structure across columns set by 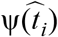, i.e. **V**_enc_ depends on the characteristic 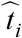 of neurons and *not* on the trial-specific instances of **G**^(*k*)^(*t*). For large enough neural sequences, Fig. 7H shows that the encoding geometry still tracks ψ(*t*) up to small trial-to-trial fluctuations, and in fact the rate of convergence vs. population size is essentially the same as before when **V**_enc_ was designed using the precise time order of neural activations in a given **G**(*t*) (Fig. 7E-right).

The reason why we did not need to carefully select the **G**(*t*) functions or coordinate them with details of **V**_enc_ can be understood via a mathematical property of high-dimensional spaces. We can think of multiplicative neural sequences as characterized by an underlying set of time-independent encoding directions **V**_enc_ in the high-dimensional neural state space (Fig. 8b-top), with geometry 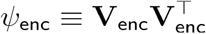. At each timepoint, random time-modulation functions **G**(*t*) approximately act to randomly project **V**_enc_ onto a low-dimensional subspace of active neurons, producing the observed encoding directions **W**_enc_(*t*) = **V**_enc_ **G**(*t*) (Fig. 8b-bottom). If the underlying **V**_enc_ directions are randomly oriented in the neural state space, the encoding geometry **U**(*t*) ≡ **W**_enc_(*t*) **W**_enc_(*t*)^⊤^ approximates the constant *ψ*_enc_ at all times *t* (Fig. S7G-H). This is because random projections in a high-dimensional space are likely to preserve the relative distances between points, and thus the relative geometry of encoding directions (Fig. S7I; a constructive proof of the Johnson-Lindenstrauss lemma (Dasgupta and Gupta 2003); see also extensions to entire manifolds (Baraniuk and Wakin 2009; Clarkson 2008; Yap, Wakin, and Rozell 2013)). Our observation that **U**(*t*) in the neural data varies gradually across time (Fig. 6A) can be thought of as originating from **V**_enc_ directions that are *not* fully randomly oriented, but instead have systematic structure w.r.t. **G**(*t*). Assuming without loss of generality that the rows of **G**(*t*) (and thus columns of **V**_enc_) are ordered by the activation times of neurons, this means that the first few columns of **V**_enc_ should have rows with a different geometry than the next few columns, and so forth. The Johnson-Lindenstrauss lemma then applies to each submatrix of **V**_enc_ defined by columns corresponding to an interval of time over which the encoding geometry **U**(*t*) is approximately constant. Within such a submatrix **V**_enc_[•; *i* … *j*], the sub-sequence of fast-timescale activations of neurons *i* to *j* produce a near-constant approximation of **V**_enc_[•; *i* … *j*], whereas the systematic change of **V**_enc_ across submatrices produces a gradual change in **U**(*t*) over longer timescales.

**Figure 8.**
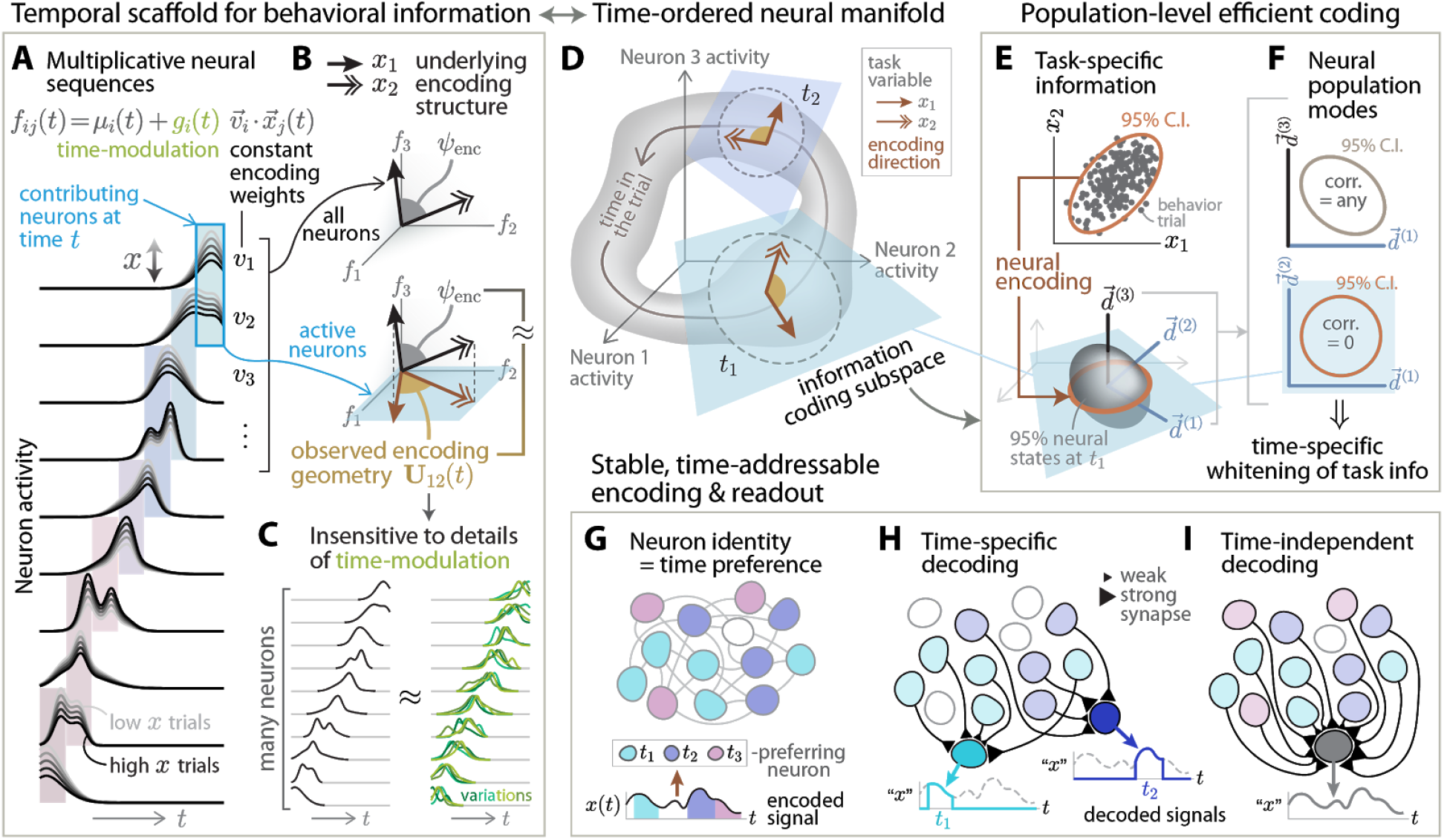
Conceptual summary: multiplicative neural sequences hypothesized to implement time-specific efficient neural-population coding of task-specific variables. **(A)** Neurons were sequentially active, with the response of each neuron *i* well-described by static task-variable encoding weights 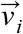 multiplied by a behavior-independent time-modulation function *g_i_* (*t*). Vertical colored bands indicate neurons that contribute significantly to the *x* encoding direction at various timepoints in the trial, each band corresponding to a different illustrated timepoint. **(B)** At each timepoint *t*, the observed encoding geometry **U**(*t*) approximates a projection of a hypothesized underlying encoding structure *ψ*_enc_ onto a low-dimensional subspace of active neurons. The underlying encoding directions (black arrows) are specified by the set of constant encoding weights 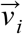 in (A), and *ψ*_enc_ is the matrix of dot products between all pairs of these encoding directions. In high dimensions (many neurons), the observed encoding geometry **U**(*t*) is likely to be nearly equal to the underlying *ψ*_enc_ up to an overall scale. See text for an explanation of how systematic structure in *ψ*_enc_ w.r.t. the preferred activation times of neurons (time sequence of projections) can generate a time-dependent **U**(*t*). **(C)** With many sequentially active neurons, the projection effect in (B) can be insensitive to details of single-neuron time-modulation functions. Specifically, for a *fixed* set of static task-encoding weights 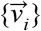 i.e. underlying *ψ*_enc_, the observed encoding geometry still approximates *ψ*_enc_ for different instances of the time-modulation functions: left traces same as (B), vs. right traces which were different randomly generated instances but maintaining a characteristic activity time window per neuron (e.g. stochasticity across trials). **(D)** Local regions of the neural manifold could be ordered by time in the trial, and at each timepoint/local region there was a different information-coding subspace spanned by encoding directions (brown arrows). **(E)** At each timepoint, task variables were correlated across trials (top plot), and part of the variance in neural states could be explained as dependence on these task variables (bottom plot). **(F)** The distribution of neural states in (E) was approximately uncorrelated within the information-coding subspace (bottom plot), but there can be signal correlations between pairs of neurons and w.r.t. other directions orthogonal to this subspace (top plot). **(G)** Since the time-modulations of neurons in (A) do not depend on task conditions, neuron identities can be used to select neurons with particular time preferences (colors). The subset of neurons with time preference *t* can then be used to encode a task variable *x* (as well as other task variables and the behavioral relationships between task variables), specific to time *t*. **(H)** Neuron identities in (G) can be used to selectively and stably read out task information at specific timepoints in the trial, i.e. a simple weighted sum of upstream neural activities (static synapses) can produce a read-out signal that is undistorted by individual neural time-modulations unrelated to task variable changes. **(I)** The union of synaptic weights for the time-specific readouts in (H) can be used to read out task information stably through time.

To summarize our proposal for how multiplicative neural sequences can implement time-dependent encoding geometries, the key requirement for the time-modulation functions **G**(*t*) is that each neuron should have a characteristic preferred activation time (cf. high reliability within their activity fields, Fig. 2H), but can otherwise have stochastic fluctuations in activity levels whether across neurons or across trials per neuron. At time *t* in the trial, a hypothetical neural circuit may then slowly adjust the strengths of task-input synapses for the subset of neurons active at time *t*, which corresponds to storing time-specific task information in a subset of columns of the time-independent **V**_enc_. This kind of “writing” into **V**_enc_ at different times *t*_1_ vs. *t*_2_ will not mutually interfere if there is no overlap in the subpopulations of neurons active at *t*_1_ vs. *t*_2_, which argues for why **G**(*t*) should have a sequential design i.e. systematic turnover in active neurons vs. time in the trial (as seen in our data, Fig. 4G,F). Multiplicative neural sequences may thus suggest a particularly robust way for neural circuits to encode the time structure of dynamic behaviors that exploits mathematical properties of high-dimensional, random designs: updates to synaptic connectivity can be made independently for subsets of neurons active at different times in the trial, and the resulting encoding geometry is insensitive to details of the neural dynamics beyond its role in determining which neurons are active at a given time.

## Discussion

In this work, we described some geometrical structures of neural-population activity across posterior cortical areas as mice performed a complex, dynamic task. How were neural representations of the many task-related variables organized relative to each other and maintained/updated through time? We answered in three parts. First, neurons were sequentially active vs. place/time in the trial (Fig. 2E), and in fact had time modulations that did not depend on task conditions (Fig. 7A), corresponding to a neural manifold with global time order (Fig. 4). Second, neural populations across the posterior cortex represented different task variables with *related* encoding directions, in particular with encoding geometries that approximated the inverse task variable covariance matrices. This supports the hypothesis that the brain’s encoding scheme approximately whitens the correlated task information, i.e. representing task variables with less correlated neural modes. Third, this encoding function was not static but reliably followed changes in task-variable correlations across time in the trial (Fig. 6A-B), which we propose can be implemented in a simple way by multiplicative neural sequences (Fig. 7C-H). Below, we discuss some implications of our findings in regards to the two questions posed in the Introduction: how neural populations simultaneously encode multiple variables, and how this neural code is dynamically coordinated and represents temporal context.

How *should* the brain encode information? Theories of efficient coding propose that to minimize resource usage (e.g. signal bandwidth), an efficient code should utilize statistically independent neural representations (Attneave 1954; Horace B. Barlow 1961). Our results support these theories, but with three distinctions:

1. We did not observe that individual neurons have statistically independent responses (Fig. S5E-G), so in a strict sense our findings differ from being a fully efficient code, as well as the focus of a large body of related work (Rieke, Bodnar, and Bialek 1995; Laughlin 1981; Dan, Atick, and Reid 1996; Baddeley et al. 1997; Vinje and Gallant 2000; Olshausen and Field 1996; Marsat and Maler 2010; Onken et al. 2014; Weliky et al. 2003; Atick and Redlich 1992). Rather, the population-level encoding directions can be thought of as spanning an information-coding subspace of the neural state space (Fig. 8D). We showed mathematically that the observed geometry of these encoding directions implies that task variables were represented by approximately uncorrelated modes within the information-coding subspace (Fig. 8E-F). In a high-dimensional neural state space, this information-coding subspace can be oriented in such a way that we can observe a spectrum of signal correlations between pairs of neurons (Fig. 8F-top; see Fig. 5L, Fig. S5F,I).
2. Although the encoding geometry was similar to the inverse task variable covariance matrix **C**^−1^ ≡ (**X**^⊤^**X**)^−1^ (Fig. 5C,F), it much better matched an alternate hypothesis 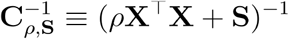 (Fig. 5D,G). The distinction is that **C**^−1^ assumes that perfect information about the task variables **X** is available for computing the inverse covariance matrix, whereas 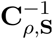 allows for different task variables to have different signal-to-noise ratios (SNR) as specified by the free parameters *ρ* and **S**. Intriguingly, all posterior cortical regions were well-characterized as having encoding geometries that matched 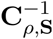, but with systematic differences in variable-specific SNR such as reduced sensitivity to visual-related task variables in higher-order cortical regions compared to V1 (Fig. 6E).
3. As opposed to purely sensory responses, we report a correspondence between the encoding geometry and the inverse covariance of a set of external and internally computed task variables, which were (only) interrelated through a learned behavioral task.

One possibility is that our three findings above arose from a learning process that optimized computational utility of the neural code, more so than low-level resource constraints. For example, (H. B. Barlow 1989; H. Barlow 2001) pointed out that utilizing a neural representation that has canceled out *expected* statistical regularities of the world permits easy detection of *unexpected* coincidences, as is relevant for survival.

As emphasized in the design of our task, another major computational challenge brains face is that the information they represent and process is not static, but changes in time depending on the environment and the animal’s own behavior. We accounted for the dynamic and nonlinear nature of neural responses by computing the encoding geometry as a function of time in the trial, equivalent to a locally-linear approximation of the structure of the neural manifold (Fig. 1). We found that locally-linear encoding models could predict neural activities nearly as well as models with higher-order (nonlinear) dependencies on task variables (Fig. S3A-B). Complementary to this, we could accurately predict the majority of task variables from neural-population activity using linear *decoders* trained on seconds-long temporal phases of the task, i.e. with accuracies comparable to per-timepoint decoders (Fig. S8B, blue vs. black traces). Accuracy comparable to per-timepoint decoders also holds when a single linear decoder per task variable was trained using data from all timepoints (Fig. S8B, red vs. black traces), albeit not for all variables because some variables such as position are highly *non*linearly coded when viewed at a global as opposed to local level. Our results are compatible with previous reports of long-timescale structure in neural state transitions in mouse posterior parietal cortex (Morcos and Harvey 2016b), long-timescale order in single-trial neural state trajectories in mouse premotor cortex (Wei et al. 2019), and timescales of stability of neural representations in monkey orbitofrontal cortex (Kimmel et al. 2020) (discussed further below). Strikingly, we observed that the efficient-coding-like function of the neural encoding geometry changed in time to approximately cancel out task variable correlations that were specific to each timepoint in the trial (Fig. 6A-B). This dynamic encoding function was not implemented by dynamical changes in neuron-neuron correlations, but instead involved a sequence of different neurons across time. We introduced a model-agnostic measure of sequentiality at the population level, i.e. the rate of angular change in the neural state, which has the intuitive interpretation that a 90° change corresponds to a complete change in active neurons. This is complementary to model-based methods in the literature that aim to fit low-dimensional dynamical systems to the neural-population data (e.g. (Churchland et al. 2012), but see also (Lebedev et al. 2019)). The angular change in neural state showed that the subset of active neurons changed continuously vs. time in the trial (Fig. 4G, Fig. S2C), and provides a quantitative measure of the sequential neural activation patterns seen in Fig. 2E.

Our observations of sequential dynamics in all areas are amongst a growing number of such reports for the mouse neocortex (Harvey, Coen, and Tank 2012; Saleem et al. 2018; Krumin et al. 2018; Morcos and Harvey 2016a; Driscoll et al. 2017; Runyan et al. 2017). These phenomena are reminiscent of place (O’keefe and Nadel 1978) or time (Pastalkova et al. 2008; MacDonald et al. 2011, 2013) cells in the hippocampus, which are also known to jointly encode a variety of other spatial and nonspatial factors. An interesting idea that has arisen in the field concerns how sequential activity could act as a temporal scaffold upon which other information can be imprinted, i.e. multiplexing this information with “timestamps” to indicate *when* they occurred (Eichenbaum 2017; Lisman 1999; Pastalkova et al. 2008; Howard et al. 2014; Jin, Fujii, and Graybiel 2009). How can such multiplexing be designed so that information can be read out without confounding the timestamp with the imprinted information? We point out a simple, robust, and interpretable design inspired by what we call multiplicative neural sequences in our data (Fig. 8A), where the response of each neuron *i* to task variables was well described as a product of two functions, 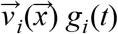. Here 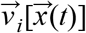 is a behavioral response function that depends on the time-dependent task variables 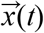 but does not otherwise depend explicitly on time, and *g_i_*(*t*) is a time-modulation function that depends only on time *t* in the trial. In other words, the nominally high-dimensional neural population activity in all surveyed cortical regions could be parsimoniously described by a low-dimensional set of multiplicative factors. This type of factorizable neural responses seems intriguingly ubiquitous, as similar findings have been reported for mouse prefrontal cortex and nonhuman-primate motor cortex (Williams et al. 2018).

Perhaps contrarily to intuition, the activity of individual neurons can be time-modulated on fast timescales and yet not preclude stable encoding and readouts on longer timescales. For neuron *i* in a multiplicative neural sequence, the proportion of activity variance explained by each task variable in 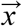 is determined by the task-variable response function 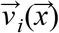, which has no explicit time dependence. This is consistent with how the selectivity of neural representations can appear stable on longer timescales than the individual neural time-variations *g_i_*(*t*) (Kimmel et al. 2020) (assuming that the measure of selectivity used is insensitive to overall activity scales as set by *g_i_*(*t*)). Population-level information about one task variable *relative* to another, e.g. the encoding geometry, can be stable on even longer timescales. This is because neural population outputs sum over the contributions of individual neurons, so contributions from one subset of neurons can be replaced by contributions from another subset, even if the subsets of active neurons change. The timescale of single-neuron activities can therefore be thought of more as limiting how quickly the neural code can change rather than how stable it can remain, subject to the trade-off that a shorter activity time window per neuron requires a larger population size in order to sustain stable output (Fig. S7A-F). In fact as we next discuss, random and heterogeneous *g_i_*(*t*) will for mathematical reasons average out in sufficiently large neural populations, and can thus in a simple, neurobiologically plausible way be used to implement neural representations that vary smoothly over long timescales.

As each neuron in a multiplicative neural sequence has a characteristic time-preference that does not depend on task conditions, static task-input synapses onto a subset of neurons with a given time-preference can be used to encode task information at just that specific time in the trial (Fig. 8G; i.e. near-independently for distal timepoints). From a circuit design perspective, the use of different neural subpopulations to encode information at different times allows for the neural code at any *one* timepoint to be learned via modifying the synapses of the fewest number of neurons. This is a special choice out of the many ways that low-dimensional (relative to the number of neurons) representations can be constructed, which we speculate reflects the special, universal role of time in behavior. While the restricted subset of active (i.e. coding) neurons mirrors ideas in sparse coding (e.g. see (Olshausen and Field 1996; Vinje and Gallant 2000; Willmore and Tolhurst 2001)), unlike previous work on sparse codes, the phenomenon here does not appear optimized to produce particular representations of high-dimensional information such as visual stimuli. Instead, we think of multiplicative neural sequences as implementing a general temporal scaffold that permits the time structure of task variable relationships to be encoded in mechanistically the same way across a variety of behaviors.

Also of particular interest to circuit design is that with large enough neural populations, the encoding geometry can be robustly constructed to vary slowly according to the time structure of the behavior, i.e. without careful tuning to the details of more rapid time-modulations of individual neurons. We demonstrated this using simulations where neural time-modulations were randomly generated, up to requiring each neuron to only be active within one (randomly determined) time window in the trial. In these simulations, the encoding geometry approximated the design truth even though the time-modulations were not coordinated across neurons (Fig. 7E-F), and even if activity levels fluctuated stochastically across trials per neuron (Fig. 7G-H). Since details of the neural time-modulations do not matter, we can think of them as functionally equivalent to simple on/off functions per neuron (Fig. 7E vs. F). Mathematically, this means that the time-modulations approximately act to randomly project the underlying encoding structure determined by static input synapses (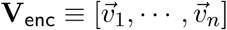, Fig. 8B-top) onto a low-dimensional subspace corresponding to the active neurons (Fig. 8B-bottom). The relevance of the projection effect is that according to the Johnson-Lindenstrauss lemma (Dasgupta and Gupta 2003), random projections are likely to preserve the relative geometry of the underlying **V**_enc_. Due to this mathematical property, the observed encoding geometry **U**(*t*) at each timepoint *t* can closely follow the structure of the static 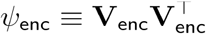, with little dependence on the fast time-modulations of individual neurons (within a band in the schematic Fig. 8A) beyond how the slower turnover in active neurons (across bands in Fig. 8A) selects different subsets of **V**_enc_ to be reflected in **U**(*t*). Abrupt changes in **U**(*t*) can also be implemented via rapid changes in the subset of active neurons (Fig. S7D-F). Our point here about the theoretical utility of random sequential neural codes extends the literature on how neural circuits can exploit random designs to perform interesting computational functions, such as separable neural representations (Fusi, Miller, and Rigotti 2016; Rigotti et al. 2013; Babadi and Sompolinsky 2014; Lindsay et al. 2017), short term memory of input patterns and dynamics (S. Ganguli and Sompolinsky 2010; Charles, Yin, and Rozell 2017; Jaeger and Haas 2004; Bouchacourt and Buschman 2019), and unsupervised learning of the structure of input signals (Maoz et al. 2020).

To summarize, utilizing a complex mouse behavior that involved a set of interrelated sensorimotor, memory- and decision-related variables, we observed that the encoding geometry of neural populations across the posterior cortex implied a time-specific whitening of these correlated task information (Fig. 6A-B). The multiplicative neural sequences observed in our data suggest a form of time-behavior multiplexing that enables large neural populations to implement this kind of time-dependent encoding function in a mechanistically simple way (Fig. 8G). This multiplexing does not require careful coordination of neural dynamics to ensure that time modulations are not confounded for changes in task variables, but rather can exploit a random design to average out the time dependence given a sufficiently large neural population. For the same reason, task information contained in neural activity can be stable against trial-to-trial variations in activity levels, e.g. due to biophysical sources of stochasticity and/or other (task-unrelated) neural processes. Complementary to this encoding function, computing a weighted sum of activities of a subpopulation of neurons with time preferences around *t* can allow a readout circuit to detect that a behavioral signal has occurred specifically at time around *t* (Fig. 8H), or task information can also be read out in a time-independent way by sampling neurons with a range of time preferences (Fig. 8I). We propose that the above computational properties of multiplicative neural sequences underlie dynamic efficient coding by neural modes across posterior cortex, and may in general be a useful design principle for temporal scaffolds in the brain.

## Online Methods

### Animals

All procedures were approved by the Institutional Animal Care and Use Committee at Princeton University (protocol #1910) and were performed in accordance with the Guide for the Care and Use of Laboratory Animals (National Research Council et al. 2011). We used 11 mice aged 2-16 months of both genders, and from two transgenic strains that express the calcium-sensitive fluorescent indicator GCamp6f (Chen et al. 2013) in excitatory neurons of the neocortex. 6 mice were of the Thy1-GP5.3 (Dana et al. 2014) strain (Jackson Laboratories, stock #028280), and 5 were crosses of the Ai93-D;CaMKII α-tTA (Madisen et al. 2015) and Emx1-IRES-Cre (Gorski et al. 2002) strains (Jackson Laboratories, stocks #024108 and #005628). All the data analyzed in this work were from fully-trained mice as described in the following sections.

### Surgery

Young adult mice (2-3 months of age) underwent aseptic stereotaxic surgery to implant an optical cranial window and a custom lightweight titanium headplate under isoflurane anesthesia (2.5% for induction, 1-1.5% for maintenance). Mice received one pre-operative dose of meloxicam subcutaneously for analgesia (1 mg/kg) and another one 24 h later, as well as peri-operative intraperitoneal injection of sterile saline (0.5cc, body-temperature) and dexamethasone (2–5 mg/kg). Body temperature was maintained throughout the procedure using a homeothermic control system (Harvard Apparatus). After asepsis, the skull was exposed and the periosteum removed using sterile cotton swabs. A 5mm diameter craniotomy approximately centered over the parietal bone was made using a pneumatic drill. The cranial window implant consisted of a 5mm diameter round #1 thickness glass coverslip bonded to a steel ring (0.5mm thickness, 5mm diameter) using a UV-curing optical adhesive. The steel ring was glued to the skull with cyanoacrylate adhesive. Lastly, a titanium headplate was attached to the cranium using dental cement (Metabond, Parkell).

### Behavioral task

After at least three days of post-operative recovery, mice were started on water restriction and the Accumulating-Towers training protocol (Pinto et al. 2018), summarized here. Mice received 1-2mL of water per day, or more in case of clinical signs of dehydration or body mass falling below 80% of the pre-operative value. Behavioral training started with mice being head-fixed on an 8-inch Styrofoam® ball suspended by compressed air, and ball movements were measured with optical flow sensors. The VR environment was projected at 85Hz onto a custom-built Styrofoam® toroidal screen and the virtual environment was generated by a computer running the Matlab (Mathworks) based software ViRMEn (Aronov and Tank 2014), plus custom code.

For historical reasons, 3 out of 11 mice were trained on mazes that were slightly longer (30cm pre-cue region + 250cm cue region + 100-150cm delay region) than the rest of the cohort (30cm pre-cue region + 200cm cue region + 100cm delay region). No qualitative differences were observed in the results of this paper across these ranges of maze lengths, and the data were thus analyzed on equal footing as described in the “Time binning” section below. In VR, as the mouse navigated down the stem of the maze, tall, high-contrast visual cues appeared along either wall of the cue region when the mouse arrived within 10cm of a predetermined cue location. These locations were drawn randomly per trial according to a spatial Poisson process with 12cm refractory period between consecutive cues on the same wall side. Cues were made to disappear after 200ms. The mean number of majority:minority cues was 8.5:2.5 for the 250cm cue region maze and 7.7:2.3 for the 200cm cue region maze. Mice were rewarded with ≥ 4μ*L* of a sweet liquid reward (10% diluted condensed milk, or 15% sucrose) for turning down the arm on the side with the majority number of cues. Correct trials were followed by a 3s-long inter-trial-interval (ITI), whereas error trials were followed by an indication sound and an additional 9s time-out period. To discourage a tendency of mice to systematically turn to one side, we used a de-biasing algorithm that adjusts the probabilities of sampling right- vs. left-rewarded trials (Pinto et al. 2018).

### Functional identification of visual areas

We adapted methods (Garrett et al. 2014; Kalatsky and Stryker 2003; Zhuang et al. 2017) to functionally delineate the primary and secondary visual areas using widefield imaging of calcium activity paired with presentation of retinotopic stimuli to awake and passively running mice. We used custom-built, tandem-lens widefield macroscopes consisting of a back-to-back objective system (Ratzlaff and Grinvald 1991) connected through a filter box holding a dichroic mirror and emission filter. One-photon excitation was provided using a blue (470nm) LED (Luxeon star) and the returning green fluorescence was bandpass-filtered at 525 nm (Semrock) before reaching a sCMOS camera (Qimaging, or Hamamatsu). The LED delivered about 2-2.5mW/cm^2^ of power at the focal plane, while the camera was configured for 20-30Hz frame rate and about 5-10μm spatial resolution. Visual stimuli were displayed on either a 32″ AMVA LED monitor (BenQ BL3200PT), or the same custom Styrofoam® toroidal screen as for the VR rigs. The screens were placed to span most of the visual hemifield on the side contralateral to the mouse’s optical window implant. The space between the headplate and the objective was covered using a custom made cone of opaque material.

The software used to generate the retinotopic stimuli and coordinate the stimulus with the widefield imaging acquisition was a customized version of the ISI package (Juavinett et al. 2017) and utilized the Psychophysics Toolbox (Brainard 1997). Mice were presented with a 20° wide bar with a full-contrast checkerboard texture (25° squares) that inverted in polarity at 12 Hz, and drifted slowly (9° /s) across the extent of the screen in either of four cardinal directions (Zhuang et al. 2017). Each sweep direction was repeated 15 times, totaling four consecutive blocks with a pause in between. Retinotopic maps were computed similarly to previous work (Kalatsky and Stryker 2003) with some customization that improved the robustness of the algorithms for preparations with low signal-to-noise ratios (SNR). Boundaries between the primary and secondary visual areas were detected using a gradient-inversion-based algorithm (Garrett et al. 2014), again with some changes to improve stability for a diverse range of SNR.

### Two-photon imaging during VR-based behavior

The virtual reality plus two-photon scanning microscopy rig used in these experiments follow a previous design (Dombeck et al. 2010). The microscope was designed to minimally obscure the ~ 270° horizontal and ~ 80° vertical span of the toroidal VR screen, and also to isolate the collection of fluorescence photons from the brain from the VR visual display. Two-photon illumination was provided by a Ti:Sapphire laser (Chameleon Vision II, Coherent) operating at 920nm wavelength, and fluorescence signals were acquired using a 40x 0.8 NA objective (Nikon) and GaAsP PMTs (Hamamatsu) after passing through a bandpass filter (542/50, Semrock). The amount of laser power at the objective used ranged from ~40-150mW. The region between the base of the objective lens and the headplate was shielded from external sources of light using a black rubber tube. Horizontal scans of the laser were performed using a resonant galvanometer (Thorlabs), resulting in a frame acquisition rate of 30Hz and configured for a field of view of approximately 500 × 500μ*m* in size. Microscope control and image acquisition were performed using the ScanImage software (Pologruto, Sabatini, and Svoboda 2003). Data related to the VR-based behavior were recorded using custom Matlab-based software embedded in the ViRMEn engine loop, and synchronized with the fluorescence imaging frames using the I2C digital serial bus communication capabilities of ScanImage. A single field of view at a fixed cortical depth and location relative to the functional visual area maps was continuously imaged throughout the 1-1.5 hour behavioral session. The vasculature pattern at the surface of the brain was used to locate a two-photon imaging field of view (FOV) of interest.

### Identification of putative neurons

All imaging data were first corrected for rigid brain motion by using the Open Source Computer Vision (OpenCV) software library function `cv`::`matchTemplate`(). Fluorescence timecourses corresponding to individual neurons were then extracted using a deconvolution and demixing procedure that utilizes the Constrained Non-negative Matrix Factorization algorithm (CNMF (Pnevmatikakis et al. 2016)). A custom, Matlab Image Processing Toolbox (Mathworks) based algorithm was used to construct initial hypotheses for the neuron shapes in a data-driven way. In brief, the 3D fluorescence movie was binarized to mark significantly active pixels, then connected components of this binary movie were found. Each of these components arose from a hypothetical neuron, but a neuron could have contributed to multiple components. A shape-based matching procedure was used to remove duplicates before using these as input to CNMF. The “finalized” components from CNMF were then selected post-hoc to identify those that resembled neural somata, using a multivariate classifier with a manual vetting step.

### Dataset selection

Per session, we computed the percent of correct choices using a sliding window of 100 trials and included the dataset for analysis if the maximum performance was ≥ 65%. All trials where the mouse performed the Accumulating-Towers task at the abovementioned difficulty levels were used for the results in this paper, excluding trials where the mouse’s navigational trajectory grossly deviated from running down the maze (view angle magnitude > π/2 anywhere in the cue region, or if the mouse backtracked in *y* position and re-entered the delay region after first crossing the midpoint of the delay region).

For stability of regression models, we excluded a further 0.4% of trials that had outlying leverage scores according to the following algorithm. Leverage measures how much the regressor values of a single trial can influence a linear regression prediction 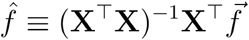, where 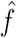 are the predicted values of the observations 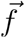 across trials, and **X** is a trial-by-variable matrix of regressors (task variables). The leverage score for trial *i* is defined as 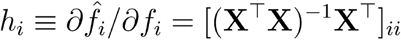, and notably depends only on the regressor values and not the observations. We therefore eliminate high-leverage trials because they cause instabilities i.e. a large change in predictions depending on whether a single trial was included or omitted from the dataset. Importantly, selecting trials based on leverage does *not* bias the distribution of observations (neural responses). To detect outliers in leverage scores for a given set of trials, we first computed a histogram of the scores (using the Freedman-Diaconis rule for selecting bin widths (Freedman and Diaconis 1981)). We used this histogram density estimate to find the modal (maximum density) leverage score 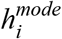, and estimated the range of typical values as the score 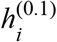 at which the density falls to 0.1 × the modal density (with 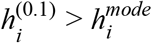). All trials with score 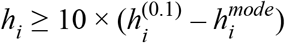 or *h_i_* ≥ 0.8 were then eliminated from the dataset. Because these criteria depend on the current set of trials used to evaluate them, the entire procedure was then iterated until no additional trials were eliminated.

### Time binning and task variable definitions

To compare data across mice and trials/mazes of uneven durations, we resampled the neural and behavioral data according to a time-like coordinate that measured progression through spatially defined epochs of the task (as indicated in the T-maze in Fig. 2A). These five epochs of the trial are: (1) the pre-cue period from the start of the trial until the mouse reaches the cue region; (2) the cue period which then lasts until the mouse exits the cue region; (3) the delay period until the mouse reaches the end of the T-maze stem; (4) the “turn” period up to the end of the trial; and (5) the first 3s of the inter-trial interval (i.e. ignoring the additional 9s time-out for incorrect trials). For each trial, the subset of the (neural and behavioral) time-series data within a given epoch was then divided into equally-sized time bins, and averaged within each of these disjoint bins. Different numbers of equally-spaced time bins were used for the five different epochs, so as to have approximately the same duration for all time bins throughout the trial. For Fig. 2E and Fig. S8 the time-bins were ~200ms (72 bins) and for all other analyses ~1.1s (11 bins) in duration.

Because it is possible for neural responses to have laterality preferences, we consistently expressed all variables relative to the brain hemisphere that was recorded from for a given mouse. That is, we defined choice, view angle, and treadmill velocity variables such that a positive sign corresponds to the mouse turning to the side ipsilateral to the recorded hemisphere. View angle is a circular variable that we represented in the range [−2π, +2π] radians, where 0 corresponds to straight down the stem of the T-maze and values with magnitude > π means that the mouse has rotated the VR view around (this essentially only happens after the mouse makes a left/right turn and has entered the T-maze arms, where further rotations of the VR view have little behavioral consequences).

### Cross-validated sequences

These analyses utilized only correct trials and followed previous work (Harvey, Coen, and Tank 2012), but with cross-validation, i.e. the following neuron categorization and sorting criteria were performed using half the trials in a given imaging session. First, for a given neuron, we identified all timepoints during which its trial-average activity was ≥ 25% of the trial-average maximum for a minimum duration of ~ 400*ms*. The neuron was defined as choice-specific if the distribution across trials of its activity in these active periods was significantly different in right- vs. left-choice trials (two-sample t-test, two tailed *p* < 0.05). A ridge-to-background excess was then defined using the activity averaged over only preferred-choice trials (or all trials, for non-choice-specific neurons), as the maximum minus the modal value across time. A neuron was determined to have significantly task-localized activity if no more than 5% of 1000 null hypothesis pseudo-datasets have a larger ridge-to-background excess. Each of these pseudo-datasets was generated by selecting a random imaging frame *k* in the session, and then defining a cyclically permuted pseudo activity time-series as [*f _i_*(*k*), *f _i_*(*k* + 1),…, *f _i_*(*n*), *f _i_*(1),…, *f _i_*(*k* − 1)] where *f _i_* (1…*n*) is the original time-series data for neuron *i*.

Using half of the trials in a given session, the preferred time of a neuron was defined as the time in the trial when its trial-average activity was maximal, and its activity field was defined as all contiguous time-points around this preferred time that have trial-average activity ≥ 50% of the maximum. For the sequence display Fig. 2E, neurons were sorted by preferred time to determine the order of rows, but the displayed trial-average activity was computed using only the left-out half of trials. Also using left-out trials, the reliability index for neuron *i* was defined as the fraction of preferred-choice trials in which the activity averaged in the neuron’s activity field is ≥ 3 times noise 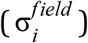. For this, the average noise level 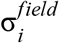 was estimated first by histogramming the distribution of the neuron’s activity *f _i_*, computing a per-imaging-frame noise estimate 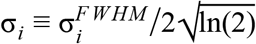 where 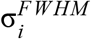 is the full-width-at-half-max of the *f_i_* distribution, and then scaling σ_*i*_ to account for averaging within the time-binned neural activity field: 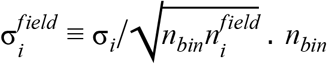. is the average number of imaging frames per time-bin (see “Time binning” section), and 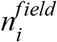 is the neuron-specific number of time bins in the identified activity field. Only significantly task-localized neurons with reliability ≥ 50% were included in Fig. 2E-G, and only significantly task-localized neurons were included in Fig. 2H. See Fig. S1 for additional statistical summaries of these measures.

### Cross-validated distribution of projected neural states

Let 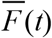 be the trial-average neural state at time *t* in the trial, computed using half of the trials for a given dataset. We defined the projection axis between two timepoints *t*_1_ and *t*_2_ as the unit vector 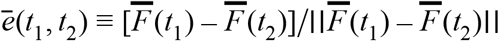. Using the other (left-out) half of trials, the neural state *F* (*t*) projected onto this axis was defined as 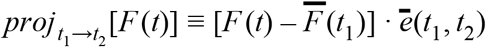, i.e. with the origin of this projection at 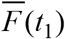. The distance along this projection axis depends on the number and activity scale of neurons, which we do not attempt to interpret. Thus for Fig. 4D we scaled the projected distributions per dataset such that 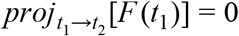 and 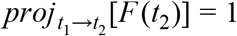, before pooling data across sessions.

As a measure of overlap between the above projected distributions, we compute the Bhattacharyya coefficient (Comaniciu, Ramesh, and Meer, n.d.) 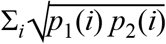, where *p*_1_(*i*) is the probability density of 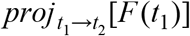 in bin *i*, and analogously *p*_2_(*i*) is the probability density of 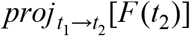 in bin *i*. 101 bins in the range [−5, 5] were used for evaluating the density histogram for this metric.

### Correction for false discovery rate (FDR)

In all cases, we used the Benjamini-Hochberg procedure (Benjamini and Hochberg 1995) to control FDR at an α = 0.05 level, as follows. We sorted the *p*-values of a given set of hypothesis tests in ascending order, [*p*_1_, *p*_2_,…, *p_n_*], and found the first rank *i*_α_ such that 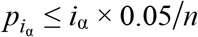. Tests were then considered to be significantly above chance (rejecting null hypotheses) for all 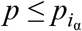.

### Per-timepoint encoding models

All encoding models used neural and behavioral data that were z-scored per timepoint in the trial, i.e. the time-dependent mean was subtracted and then the data divided by the time-dependent standard deviation. Models were fitted separately per timepoint, and all utilize the same linear regression framework that we will first explain, but with variations on regularization and choice of regressors that we will explain next. The basic linear regression model is **F** = **XW** + noise, for which the point estimate of **W** is **W**_enc_ = (**X**^⊤^**X**)^−1^**X**^⊤^**F**. For each timepoint *t* in the trial, we computed the Singular Value Decomposition (SVD; e.g. see (William H. Press et al. 2007)) of the trial-by-variable matrix of task variable values **X**(*t*) = **USV**^⊤^, then used the pseudo-inverse to solve for encoding directions **W**_enc_(*t*) ≡ **VS**^−1^ **U**^⊤^ **F**(*t*) where **F**(*t*) is a trial-by-neuron matrix of neural activity levels. Singular values ≤ 10^−5^ of the largest singular value were set to 0 in **S**^−1^.

For assessing how well encoding models could predict the activity of individual neurons (Fig. S3, Fig. 7A-B), we regularized the above linear regression solution independently per neuron using ridge regression. Specifically, the ridge-regularized encoding weights for neuron *i* is 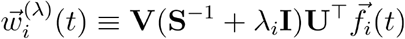 where 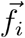 is the *i* ^th^ column of **F**. The regularization hyperparameter was chosen per neuron to maximize the 10-fold cross-validated variance explained, 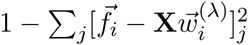. Here 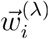 was computed using a randomly selected 1/10^th^ of trials, and the sum is over all trials *j* in the left-out 9/10^th^ of trials. The average variance explained across 2 resamplings of 10 cross-validation folds (i.e. 20 computations in total) was used to select λ_*i*_. After λ_*i*_ had been chosen, another (i.e. independent) draw of 2 × 10 cross-validation folds were used to compute all reported goodness-of-fit scores for the model.

For models with higher-order dependencies on task variables, we augmented the regressor matrix with 2^nd^-order terms [**X X**^[2]^] (Fig. S3A) or up to 3rd-order terms [**X X**^[2]^ **X**^[3]^] (Fig. S3B-E). Here 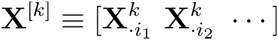, where 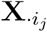 are the columns of **X** corresponding to non-binary variables, and 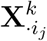 were computed by taking the *k* ^th^ power of the entries of 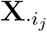, then orthogonalizing 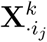 w.r.t. 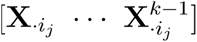 (modified Gram-Schmidt procedure).

### Encoding geometry estimates

To robustly estimate the geometrical relationships between encoding directions, we developed a procedure that optimally determines a geometry-level regularization, as well as corrects for a finite-sample noise offset which would not otherwise average to zero across experiments. Suppose that in a given experiment, the trial-by-neuron population activity data matrix **F** came from a generative process such that **F** = **XW** + *ε*, where **X** is a trial-by-variable matrix of task variable values, **W** is a variable-by-neuron matrix of true encoding weights, and *ε* is a random noise matrix unrelated to **X**. The unregularized regression weights **W**_enc_ ≡ (**X**^⊤^**X**)^−1^ **X**^⊤^ **F** is an asymptotically unbiased estimator of **W**, but the variance of **W**_enc_ across experiments (i.e. different instances of *ε*) can be large in the realistic case where the number of trials is *not* order(s) of magnitude greater than the number of free parameters. Regularization is a standard way of reducing variability at the cost of introducing some bias into estimators. However, if we independently regularize the encoding weights for each neuron, this leads to a net bias on the naıve population-level encoding geometry estimator 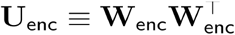 that is difficult to intuit, and likely suboptimal compared to optimizing a geometry-specific regularizer (“no free lunch” theorem (Wolpert and Macready 1997)). In the following, we thus describe an algorithm for selecting an encoding geometry regularizer based on explicitly modeling the bias-variance tradeoff to be optimized. Furthermore, the naıve estimator has contributions from second-order statistics of the neural noise fluctuations, *εε*^⊤^, which we must also correct for. These noise contributions are also a form of bias in **U**_enc_, but for clarity we refer to them as a “noise offset” to distinguish them from the (unrelated) regularization-induced bias. The following procedure mitigates both of the above issues with **U**_enc_, and we present some simulation tests at the end of this section.

We first address the noise offset, which arises from neural noise *ε* contributions to the estimated encoding directions **W**_enc_ = **W** + **C**^−1^ **X**^⊤^ *ε*, where for brevity, **C**^−1^ ≡ (**X**^⊤^**X**)^−1^. The noise offset in the naıve encoding geometry estimator is the average deviation of **U**_enc_ from the true encoding geometry **WW**^⊤^ over many experiments (indicated by 〈 … 〉):

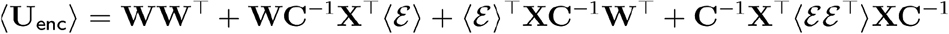

We assume that the neural noise comes from a distribution with zero mean, 〈*ε*〉 = 0 (recall that the neural and behavioral data were z-scored so that we did not need to include a constant term in the encoding models). This gives:

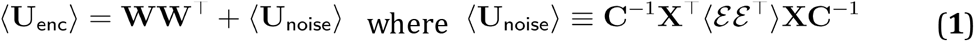

The noise offset 〈**U**_noise_〉 can be thought of as arising from random coincidences of the neural noise *ε* with the task variable values **X**. Although we do not know what *ε* is for a particular experiment, we can nevertheless estimate the experiment average 〈*εε*^⊤^〉. The idea is that instead of taking **X** to be fixed and calculating random coincidences with different possible *ε*, we can keep the unknown *ε* fixed and calculate random coincidences with different randomly generated **X** that preserves the structure of the original problem.

To do this, we generated an ensemble of 100 random matrices **X**_∗_ with normally distributed entries subject to two constraints: 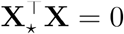, i.e. **X**_∗_ are unrelated to the original task variables; and 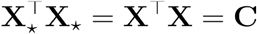, i.e. **X**_∗_ has the same covariance as the original task variables. Using the neural data, we then computed 100 linear regression estimates for the encoding of, which theoretically have the noise-only form:

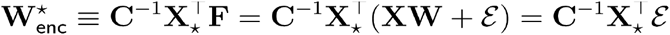

We used 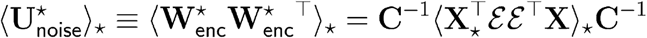, where 〈…〉_∗_ indicates averaging across the randomly generated **X**_∗_, as an estimator of the experiment-average **U**_noise_ ≡ **C**^−1^ **X**^⊤^ 〈*εε*〉 **XC**^−1^. That is, we defined a noise-offset-corrected encoding geometry estimator 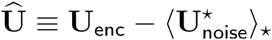, which in simulations where the “neural” data consisted solely of noise produces 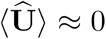.

Beyond subtracting the experiment-average noise offset, we also wished to reduce noise-induced contributions to the estimated encoding geometry in any one experiment. We achieved this in analogy to Tikhonov-Miller regularization (Tikhonov et al. 1977), where instead of using the inverse task covariance matrix to estimate **W**_enc_ ≡ **C**^−1^ **X**^⊤^ **F**, a regularized inverse is used to estimate regression weights as 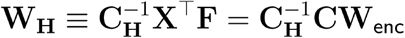, where **C**_**H**_ ≡ **X**^⊤^ **X** + **H** for some user-specified choice of the symmetric regularizer matrix. Similarly, we define a regularized and noise-offset-subtracted estimate of the encoding geometry as:

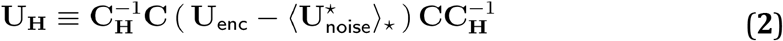

In the ideal world where we could repeat the same experiment many times, we would select the hyperparameters to minimize fluctuations of these per-experiment estimates from the true encoding geometry: 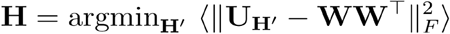 where ||**A**_*F*_|| is the Frobenius norm of matrix **A**. To obtain **H** in practice, we need to make two approximations. First, we approximate the unknown true encoding geometry **WW**^⊤^ with our best guess so far, 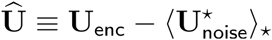. Second, we approximate the distribution across *experiments* of, with the distribution across *noise fluctuations* 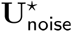 of a noise-contaminated encoding geometry. Specifically, we optimized:

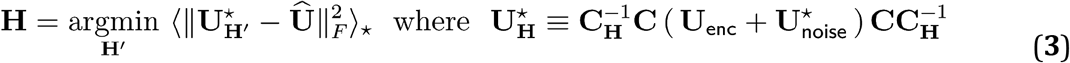

In order to make this optimization program well-defined, we have to constrain the form of **H**. We tried three fairly standard forms:

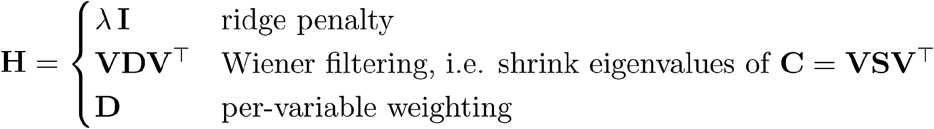

where *λ* is a scalar and **D** a diagonal matrix of free parameters. For each dataset, we selected the best form of **H** as that which produced the smallest 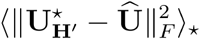 after minimization. We note that Eq. 3 was used only to determine the hyperparameters **H**, which was then plugged into Eq. 2 for the actual encoding geometry estimate. In our simulations, the final results were actually fairly robust to how we chose to model the distribution of **U**_**H**_ − **WW**^⊤^, e.g. bootstrapping worked comparably well.

Lastly, the estimator Eq. 2 is by design biased, and we wished to improve it by correcting for an approximate *expected* bias, as inspired by (Shen, Xu, and Li 2012). Bias is defined as the experiment-average deviation of a given estimator from the true encoding geometry, **B**_**H**_(**WW**^⊤^) ≡ 〈**U**_**H**_ − **WW**^⊤^〉 for a particular choice of **H** as explained above. Plugging in Eq. 1 and Eq. 2, we get:

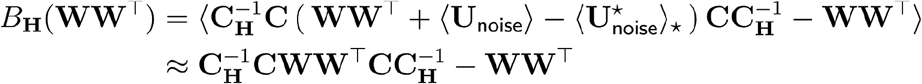

Although (as suggested by the notation) the bias is a function of the unknown truth **WW**^⊤^, we can estimate it (Shen, Xu, and Li 2012) by replacing **WW**^⊤^ with our best guess so far, **U_H_**. This means that our final, bias-reduced estimator for the encoding geometry (as used throughout the text) is:

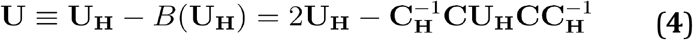

For the interested reader who is looking for structure in very few datasets, it can be preferable to use more aggressive regularization so that results averaged across the small number of datasets are less likely to deviate from the truth; this of course comes at a cost of not being able to resolve weakly encoded signals. For this purpose, the above procedure of determining the regularizer matrix **H** can be iterated by replacing the initial guess 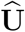 for the “true” encoding geometry in Eq. 3, with the best estimate **U** at the end of the previous iteration (Eq. 4). Here, use of the bias-corrected estimate in Eq. 4 is crucial because if the pre-correction Eq. 2 (**U**_**H**_) is used instead, the bias-induced shrinkage will accumulate with more and more iterations, eventually resulting in a near-zero estimated **U**. In contrast, in our simulation tests iterating the optimization in Eq. 3 while setting 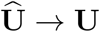 after each iteration converges to a nonzero encoding geometry estimate for all but the most weakly encoded task variables, with low variance across repeated simulations as desired. To detect convergence in this iterative procedure we checked for 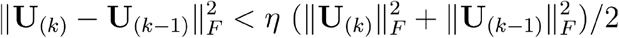, where **U**_(*k*)_ is the estimated encoding geometry at the end of iteration *k*, and η is a small tolerance threshold (a smaller value of η will result in more aggressive regularization). For simplicity and in light of the large number of datasets in our hands, we did *not* use an iterated procedure for the results described in the text.

In Fig. S4, we illustrate how well the above encoding geometry estimation algorithm works to recover the simulated ground truth in scenarios where (1) neurons linearly encoded task variables, and also explore what we might have found if (2) neurons randomly but *nonlinearly* encoded task variables. To stay close to conditions in our data, for these simulations we used the same behavioral data **X**(*t*) and neural population sizes as in our actual experiments. Specifically, for each experimental dataset of *n* simultaneously recorded neurons across *m* trials, we defined 11 simulated experiments where in each simulation, we generated a *n* -by- *m* “neural” matrix **F** (in ways described next) and asked what its encoding geometry is w.r.t. **X**(*t*) from the data (where *t* is one of 11 timepoints that define the data). The total number of simulations is thus 143 × 11 (datasets × timepoints, because each timepoint was treated like a different dataset). For the linear encoding scenario, we simulated **F** = *ρ*_sim_**XW**_sim_ + *ε*, where each entry of **W**_sim_ was drawn from the normal distribution except that the columns of **W**_sim_ corresponding to choice and past-choice task variables were set to 0, and each entry of the noise matrix *ε* was also drawn from the normal distribution. An example of such a simulated **W**_sim_ is shown in the top row of Fig. S4A, where each column corresponds to a different setting of the scalar parameter *ρ*_sim_ so that the neuron-average signal-to-noise (SNR) variance has a prespecified value (10% vs. 40%). The bottom row of Fig. S4A shows the estimated encoding geometry **U** (Eq. 4) in these examples, from which we see that **U** can qualitatively capture the true encoding geometry 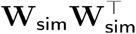, albeit low-magnitude entries of 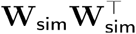 (particularly if they correspond to highly correlated task variables) are harder to resolve at lower SNR. Importantly, the near-zero columns and rows of **U** corresponding to choice and past-choice variables illustrate that our procedure correctly estimates near-zero encoding directions for non-encoded variables. This is the case even though choice (past-choice) is highly correlated to view angle (past-view-angle), the latter of which are encoded and are nonzero in **U**. Fig. S4B shows the estimated encoding geometry averaged across 143 × 11 simulations, for various choices of *ρ*_sim_ including a pure noise scenario. Because were randomly generated for each simulation, we expect the off-diagonal entries of **U** to average to zero across enough simulations, and this is what we see in Fig. S4B. Importantly, the zero-signal scenario (Fig. S4B-left) produces **U** ≈ 0, with none of the entries being statistically significantly different from the null hypothesis (see next section). At higher SNR (other columns of Fig. S4B), we see that there are no significantly nonzero choice and past-choice encoding directions, as per simulation truth. Lastly, Fig. S4C-F show simulation-average encoding geometries for scenarios where we generated neurons that were tuned to random combinations of task variables, in the sense of having gaussian-shaped tuning curves for their responses to task variables (see caption of Fig. S4C-F for a more detailed explanation). Notably, as seen in the nonzero off-diagonal entries of Fig. S4D,F, this kind of nonlinear neural code can exhibit structure in the linear encoding geometry **U** that is consistent across random instantiations (simulations), due to the structure in the encoded task variables **X**. However insofar as we can tell, this depends on details of the nonlinear code (Fig. S4C vs. Fig. S4E), and does not in some evident way explain our observations in data that the encoding geometry (Fig. 5A) resembles the inverse of a noisy task-variable covariance matrix (Fig. 5D).

### Null hypothesis for the encoding geometry

For each timepoint *t* in the trial and given a trial-by-neuron neural state data matrix **F**(*t*), we wished to generate null hypothesis pseudo-experiments that preserved (1) the time-specific distribution across trials of neural activity levels (which were positively skewed in our data); (2) inter-neuron activity correlations; (3) neural state correlations across timepoints in the trial; and (4) inter-trial temporal autocorrelations (observed to fall to zero after ~5 trials in the data). Although the typical procedure of cyclically permuting the neural state across trials can satisfy all of these three requirements, given only about 70-350 trials per session we could not use it to generate ~ 10^5^ distinct pseudo-datasets as required to evaluate *p*-values reported in the text. Furthermore, cyclic permutations applied in this way are overly conservative in that when we cyclically permuted the *behavioral* data, it preserves some level of correlations between the permuted vs. original behavioral data, and thus could not break the relationship between behavior and neural activity as required for a null hypothesis. Instead we devised a procedure (Algorithm 1), sketched conceptually below, that uses random row (trial) permutations to completely break relationships between the neural state and behavior, and then restores the trial-autocorrelations (4) by applying a row-convolution operation. To mitigate any possible finite-sample effects due to generating null hypotheses from a limited number of data trials, we further add a small amount of stochasticity to the generated pseudo-data in a way that preserves (1), (2), and (3).

Algorithm 1 assumes that the rows of **F**(*t*) are ordered according to the sequence of trials in a given experimental session. First we randomly permute the rows (trials) of **F**(*t*) to generate the pseudo-data matrix **F**_∗_(*t*), using the same permutation for every timepoint *t*. This preserves inter-neuron and cross-timepoint correlations, but destroys any autocorrelations across trials. However, we can restore the row-autocorrelations in **F**(*t*) to **F**_∗_(*t*), by convolving the rows of **F**_∗_(*t*) with a symmetric kernel or equivalently left-multiplying **F**_∗_(*t*) by a band-diagonal matrix **M**. Assuming that we seek a solution with minimal amount of mixing of the rows of **F**_∗_(*t*), i.e. **M** with small off-diagonal entries, we can obtain an optimal **M** to equate the autocorrelation of **F**_∗_(*t*) to that of **F**(*t*) by solving a system of linear equations as given in Algorithm 3. This algorithm requires specification of how many lags *L* up to which we should constrain the autocorrelation function, which we chose to be *L* = 20 based on the empirical observation that the trial autocorrelation of the data was nonzero for only up to ~5-10 trials. We also wish to add a small amount of randomness to the pseudo-data **F**_∗_(*t*), analogous to “smoothed bootstrap” procedures (Shakhnarovich, El-Yaniv, and Baram 2001) which can be thought of as drawing **F**_∗_(*t*) from a kernel density estimate of the empirical distribution of neural activity levels specified by **F**(*t*). In order to preserve temporal correlations under this procedure, the noise added to **F**_∗_(*t*) must have the same temporal correlations as the neural data. This is achieved by drawing the noise from a multivariate gaussian distribution with covariance matrix specified by the data (Algorithm 2), after which the generated noise for time *t* was provided for use in Algorithm 1 to produce a noise-added version **F**_∗,*ε*_(*t*). Lastly in order to exactly preserve the inter-neuron covariance structure, we whiten **F**_∗,*ε*_(*t*) using polar decomposition (Higham 1988), and then multiplying by the polar factor of **F**(*t*) to impose the desired covariance structure. Because modifying the column covariance of a matrix does not generally leave unchanged its row autocorrelations, we then iterate the adjustments for row-autocorrelation and column-covariances twice, which in our data are sufficient for the procedure to converge to the target structure.

Our algorithm is a variant of the Corrected Fisher Randomization method in (Elsayed and Cunningham 2017), which in our hands did not seem to preserve the distribution across trials of neural activity levels, particularly if strongly skewed or approximately bounded from below (since fluorescence contrast cannot fall far below baseline for sparsely active neurons). We empirically found that using our procedure, performing the noise addition step on a logarithmic scale (see Algorithm 1) is sufficient to qualitatively preserve the shape of activity distributions, including the bounded support.

**Algorithm 1.**
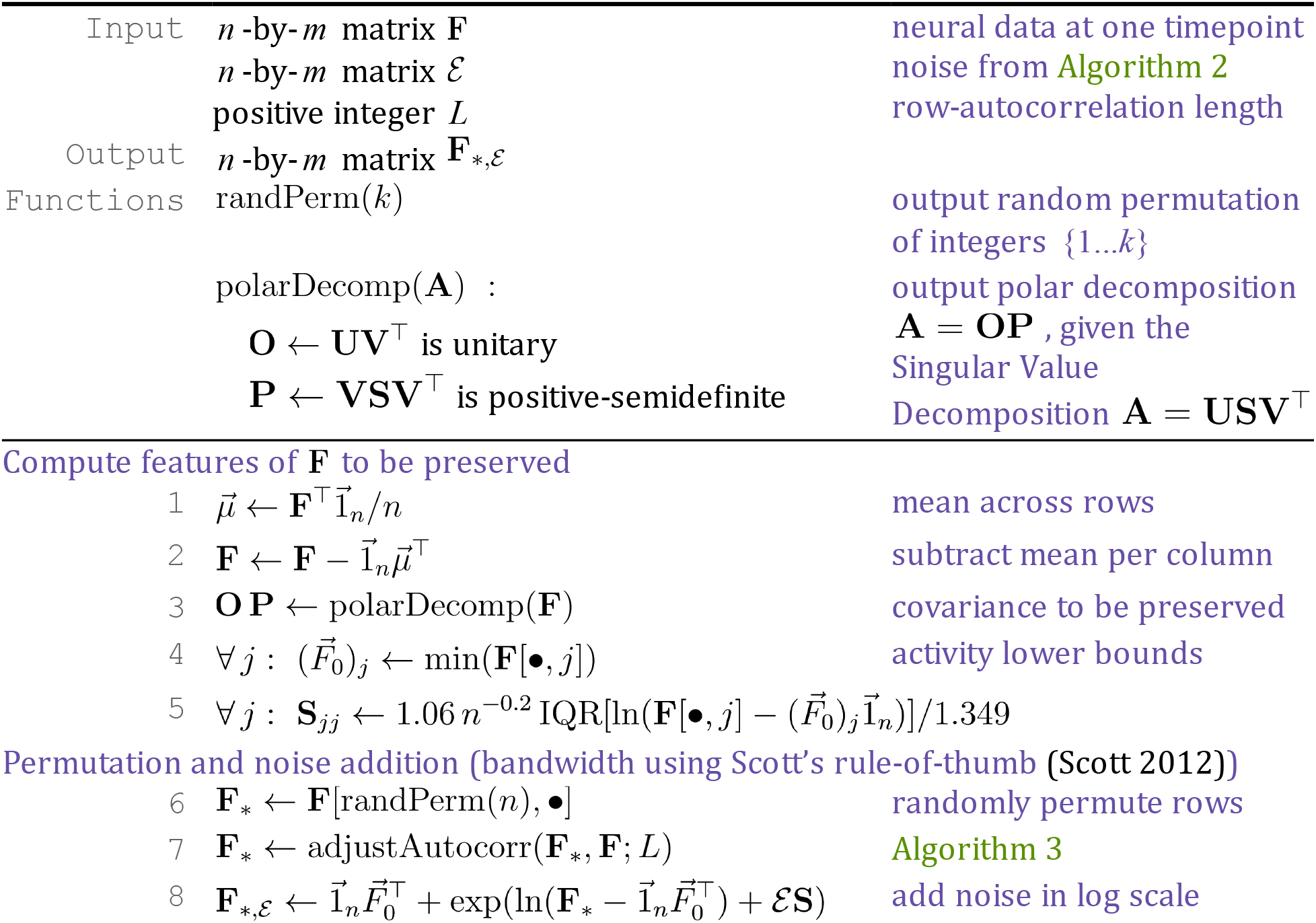
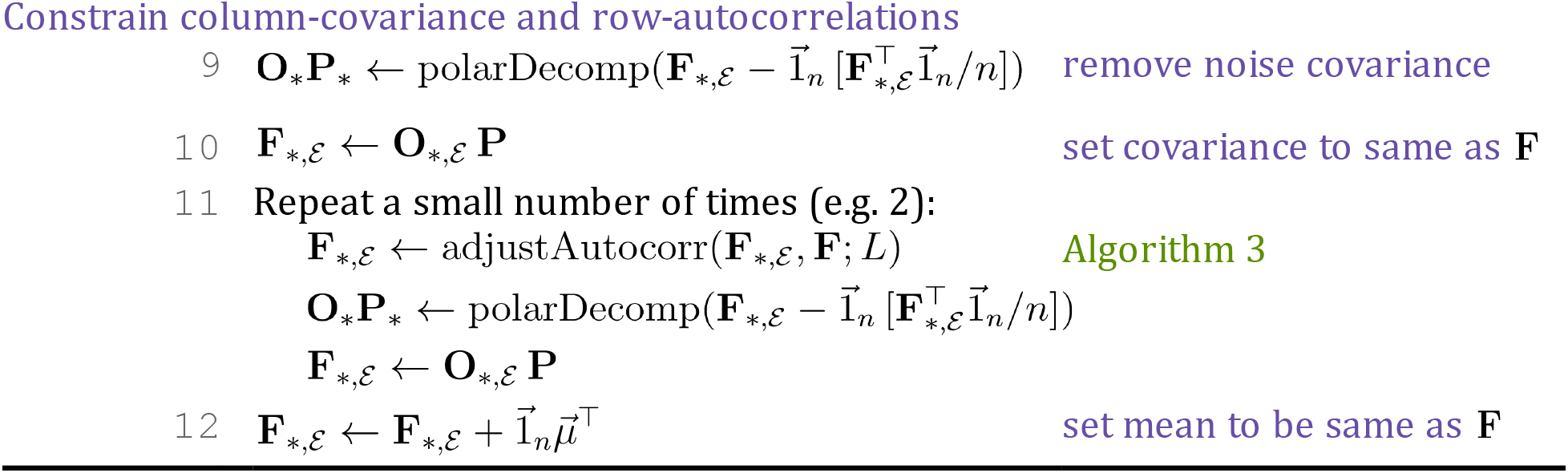
Generate one instance of null hypothesis pseudo-neural-data by randomly permuting the neural state across trials, with an added small amount of noise. 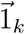 is a *k* -by-1 vector with all entries being 1. IQR(*S*) is the interquartile range of set *S*.

**Algorithm 2.**
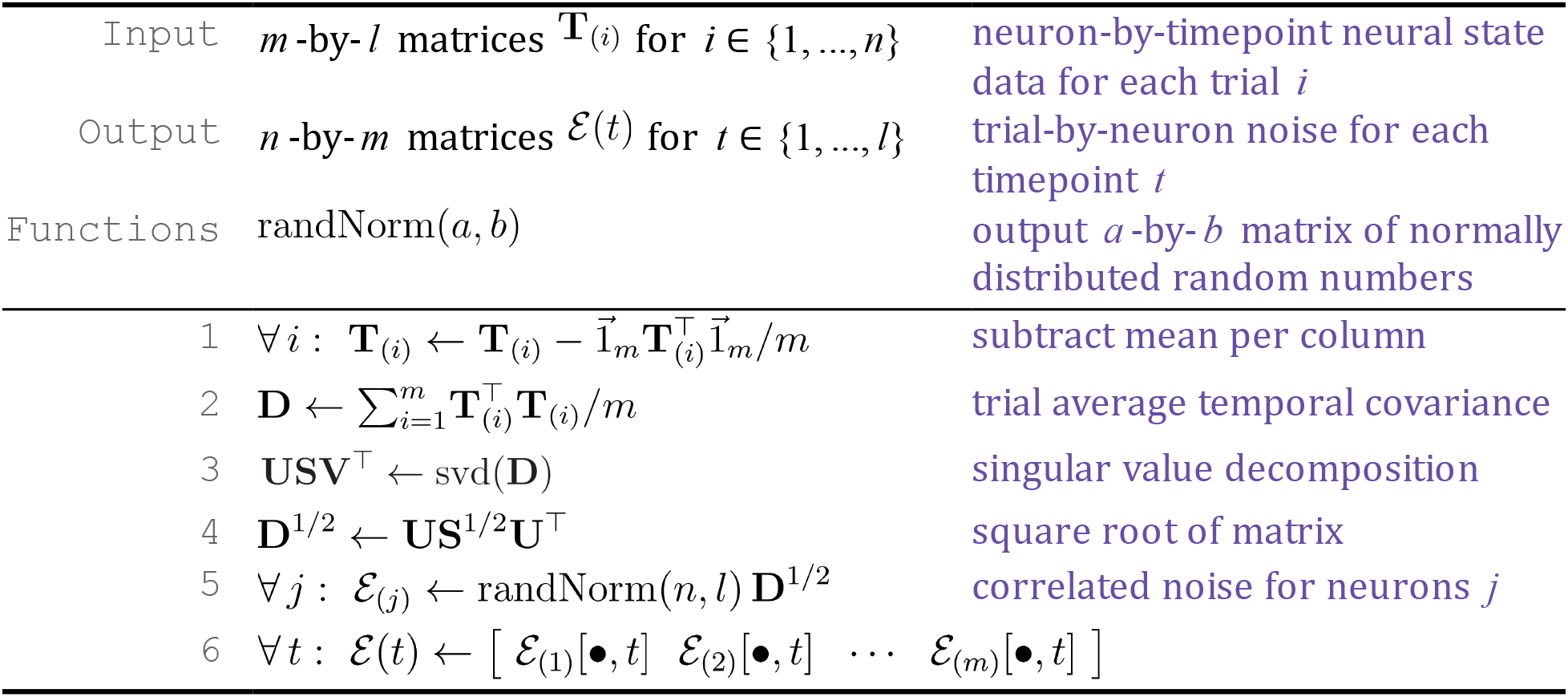
Generate temporally correlated noise.

**Algorithm 3.**
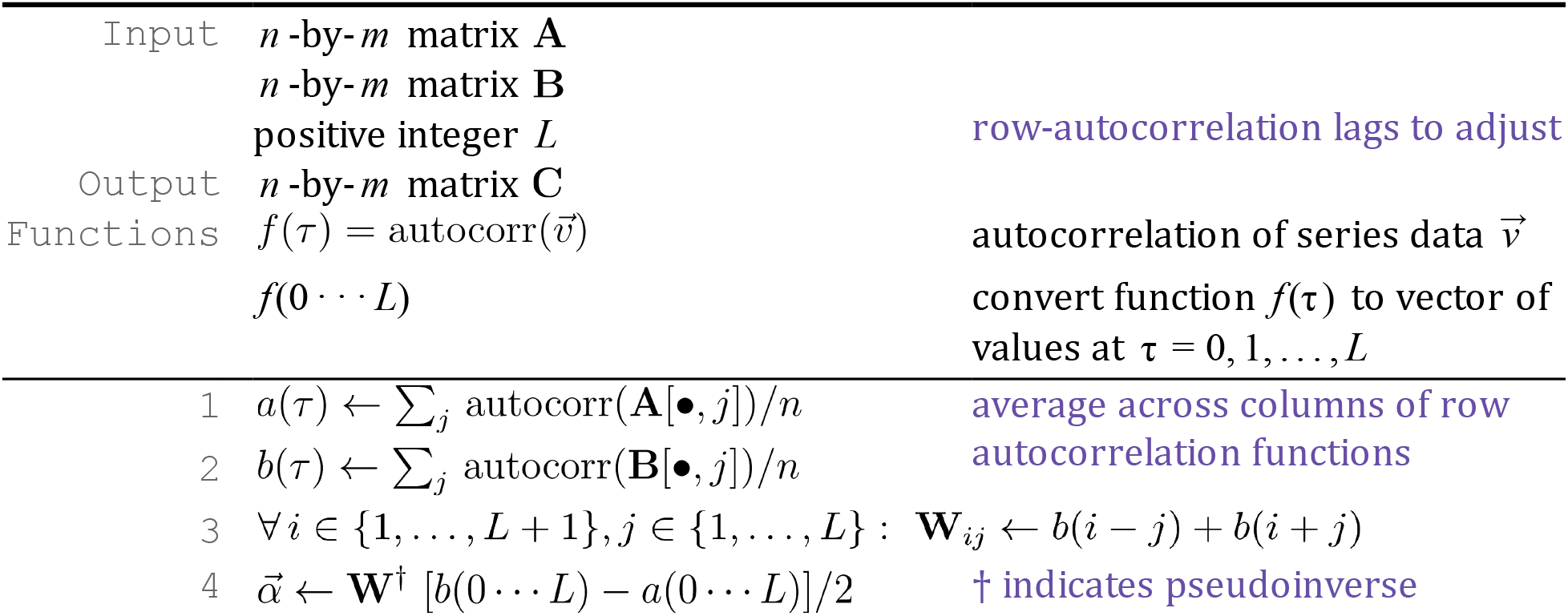
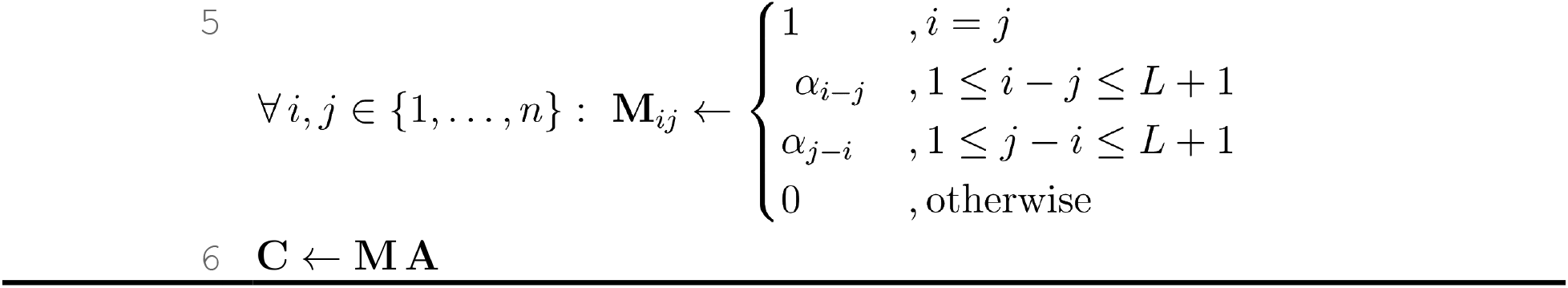
Adjust the row autocorrelation (up to *L* lags) of matrix A to match that of B.

### Statistical significance for encoding geometry

To compute a 2-sided p-value for each entry *U_ij_* of the encoding geometry matrix (Fig. 5E, Fig. 6A-bottom, Fig. S4), we accounted for both the finite number of permutation samples as well as the asymmetric distribution of the corresponding null hypotheses 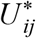. First, we estimated the 1-sided p-value for the observed *U_ij_* to be larger than null hypotheses: 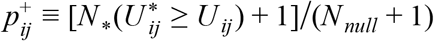, where 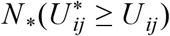 is the number of sampled null hypotheses experiments where 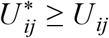, and *N_null_* = 10^5^ is the total number of sampled null hypotheses experiments. The addition of the pseudo-count 1 to both the numerator and denominator of this p-value estimate prevents the reported p-value from being 0 when insufficient permutations were performed, and produces a conservative estimate because it can be thought of as including *U_ij_* from the actual experiment as part of the distribution of null-hypotheses 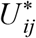 (Phipson and Smyth 2010). Similarly, we defined another 1-sided p-value for the observed *U_ij_* being smaller than null hypotheses: 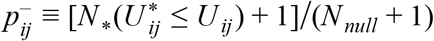. Lastly, we computed the 2-sided p-value for the observed *U_ij_* being more extreme than null hypotheses as 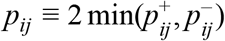, i.e. performing two 1-sided tests and then combining the p-values according to Bonferroni’s correction for multiple comparisons (Abdi 2007).

### Multiplicative time-modulation and static-encoding models

For the *i* ^th^ neuron with activity *f_ij_*(*t*) in trial *j*, we fit two alternative encoding models with constrained changes in encoding weights vs. time *t* in the trial. Here 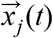 are the values of the task variables at time *t* in trial *j*, and have their time-dependent means across trials subtracted, but only scaled such that the standard deviation of each variable, computed across all timepoints as well as trials, are 1. This is in contrast to the per-timepoint encoding/decoding models, where the task variables were scaled differently per timepoint. We note that according to these models, the sensitivity of the neural-population encoding to the *time-dependent* scales of task variables (Fig. S6A-C) should be ascribed to differences in static task-variable encoding weights *across* individual neurons in the population.

The first, multiplicative time-modulation model (Fig. 7A) predicts that a neuron’s activity has the form 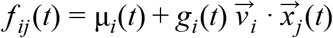, where μ_*i*_ (*t*) is a time-dependent baseline that does not depend on task variables, *g_i_*(*t*) is a piecewise-constant function with 11 free parameters for the 11 time-bins, and 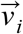 are 13 free parameters for linear dependencies on each of the 13 task variables. Since 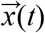 has zero mean across trials for a fixed time *t*, without loss of generality μ_*i*_(*t*) is just the trial-average mean activity level of the neuron. We estimated the 11+13 free parameters for the model by minimizing an L2-regularized least-squares cost function, where the regularization hyperparameter was selected using 10-fold cross-validation, same as for the per-timepoint encoding models. After selecting hyperparameters, a different draw of 10 cross-validation folds were used to evaluate the goodness-of-fit for this model.

The second, static-encoding model (Fig. 7B) is a special case of the first model where we constrained *g_i_*(*t*) = 1 for all timepoints, i.e. predicts neural activity of the form 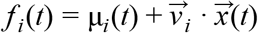. The fitting procedure for both models are otherwise identical.

### Why we call U ∝ C^−1^ an (effectively) whitening operation

Given a trial-by-variable data matrix **Y** with non-singular covariance matrix, and the transformed data **Z** = **YT**, we call **T** a whitening transformation if the transformed covariance is **Z**^⊤^ **Z** = **I**. Below, we show that constraining the encoding geometry to be U ∝ C^−1^ corresponds to such a whitening transformation for **Z** = **FD**, where **D** is an orthonormal basis (**D**^⊤^ **D** = **I**) for the brain’s encoding matrix **W**^⊤^. **Z** is the projection of the neural state **F** onto the information-coding subspace (spanned by basis vectors **D**) as discussed in the text.

For the following derivations, we will need the fact that applying the projection operator **DD**^⊤^ to **W**^⊤^ does nothing: **DD**^⊤^ **W**^⊤^ = **W**^⊤^, because the columns of **W**^⊤^ are by construction vectors that live in the subspace spanned by **D**. No other properties of **D** are required, i.e. **D** can be any orthonormal basis for the information-coding subspace. We call the projected coordinates of encoding directions in the information-coding subspace **E**^⊤^ ≡ **D**^⊤^ **W**^⊤^ (a variable-by-variable matrix).

Assuming that the neural activity depends on task variables **X** in the form **F** = **XW** + *ε*, the covariance of the neural data is **F**^⊤^ **F** ≈ **W**^⊤^ **CW** + **S** where **S** ≡ *ε*^⊤^ *ε* is the noise covariance. In the presumably typical case where there are more neurons than encoded variables, specifying the variable-by-variable matrix U ∝ C^−1^ will not fully constrain the variable-by-neuron matrix **W**, and is therefore insufficient to completely whiten the neural data in the sense of achieving **F**^⊤^ **F** ∝ **I** even with neural noise covariance **S** ∝ **I**. This is the reason why there can still be signal correlations between pairs of neurons, as explained in the text regarding Fig. 5K-L.

To understand how the encoding structure affects the covariance of **FD**, we first note that although the encoding directions **W**^⊤^ are vectors in the high-dimensional neural state space, the encoding geometry **U** depend only on the projected coordinates of these encoding directions in the information-coding subspace, **E**^⊤^ ≡ **D**^⊤^ **W**^⊤^. This is because plugging in **DD**^⊤^ **W**^⊤^ = **W**^⊤^ to **U** ∝ **WW**^⊤^ gives **U** ∝ **WDD**^⊤^ **W**^⊤^ = **EE**^⊤^. Our claim is that *any* invertible **E** that satisfies **U** ∝ **C**^−1^ will whiten **FD** (up to an overall scale, which we ignore). To see this, start from **U**^−1^ = (**E**^−1^)^⊤^ **E**^−1^ = **C**, left-multiply by **E**^⊤^ and right-multiply by **E** to get **I** = **E**^⊤^ **CE**. The covariance of the projected neural state is (**FD**)^⊤^ (**FD**) = **D**^⊤^ (**W**^⊤^ **CW** + **S**)**D**. Again assuming **S** = σ^2^**I**, we get (**FD**)^⊤^ (**FD**) = **E**^⊤^ **CE** + σ^2^**D**^⊤^ **D** ∝ **I**, as claimed. In sum, we call **U** ∝ **C**^−1^ a whitening operation because it is the constraint that if exactly satisfied, will whiten **FD** (Fig. 5K).

### Decoding models

All decoding models used neural and behavioral data that were z-scored per timepoint in the trial, i.e. the time-dependent mean was subtracted and then the data divided by the time-dependent standard deviation. Unless otherwise specified, models were fitted separately per timepoint. The exceptions are in Fig. S8B, where phase-specific decoders included data from all timepoints within the stated phases of the trial as if they were additional trials, and where time-independent decoders included data from all timepoints as if they were additional trials.

We trained an L2-regularized Support Vector Machine classifier (Fan et al. 2008) (or L2-regularized Support Vector Regression for continuous variables) to predict a given task variable from the neural population state. For categorical variables, performance was defined as the proportion of correct classifications of test trials, averaged across categories (i.e. to balance the influence of all categories). For continuous variables, performance was defined as the Pearson’s correlation coefficient between predicted and actual variable values. 10-fold cross-validation was used to optimize the regularization and support vector hyperparameters, and another set of 10-fold cross-validation folds was used to evaluate decoder performances. Null hypotheses were the same as for the encoding models (see above section), and *p*-values were defined as the fraction of null hypothesis pseudo-datasets for which the decoding performance was greater or equal than the actual experiment, with an added pseudocount to conservatively prevent zero empirical *p*-values (Phipson and Smyth 2010).

### Simulations of multiplicative neural sequences

For each simulated neuron *i*, we generated a time-modulation function *g_i_*(*t*) by first drawing a uniformly random time preference within 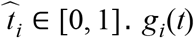. *g_i_*(*t*) was then defined as the sum of 5 gaussian bumps with peaks 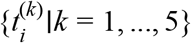 randomly distributed around this time preference 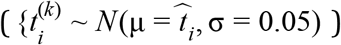. The width 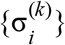 of each bump was drawn randomly: 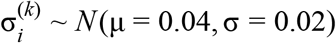 with minimum value 0.02. The neural population **G**(*t*) was then composed as a diagonal matrix the entries of which are the *g_i_*(*t*) of individual neurons.

To construct **V**_enc_ given a simulation truth ψ(*t*) as described in the text, we first ordered the simulated neurons in ascending order of either the empirical maximal response time 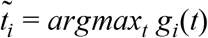 (Fig. 7E-F), or the generative time preference 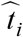 (Fig. 7G-H). This means that the rows/columns of **G**(*t*) are in temporal order, as are by implication the columns of the to-be-defined **V**_enc_. As a procedurally simple (albeit approximate) way of producing structure ψ(*t*) across the columns of **V**_enc_, we first initialized the entries of the 2-by- *n* matrix **V**_enc_ to normally distributed random numbers, i.e. generating two randomly oriented encoding directions (rows) for a population of *n* neurons. Then for each pair of adjacent columns, i.e. the submatrices **V**_enc_[•, *i* … *i* + 1] for *i* = 1, 3,…, *n* − 1, let *t*_(*i*)_ and *t*_(*i*+1)_ be the respective nominal activation times for the two associated neurons (either 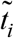 or 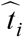 as defined above, after sorting the neurons). We orthogonalize the two rows of **V**_enc_[•, *i* … *i* + 1], producing unit vectors 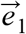 and 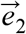, and then set them to have the desired cosine angle via linear combination: 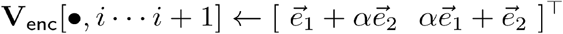, where 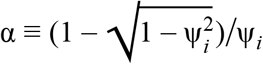 and *ψ_i_* ≡ *ψ*([*t*_(*i*)_ + *t*_(*i*+1)_]/2). Note that although by construction ψ(*t*) specifies the *cosine* angle between encoding directions, for clarity we convert it to the angle in degrees for display in Fig. 7E-H and Fig. S7.

## Acknowledgements

B. Engelhard and L. Pinto built rigs for high-throughput training of mice, and S. Stein helped in the training of mice in this study. We thank S. Lewallen, R. Low, and M. Schottdorf for impetus to dive into the strange new world of high-dimensional geometry, and all members of the BRAIN COGS team, and present and past members of the Tank and Brody labs for guidance, feedback, and science. This work was supported by the NIH grants 5U01NS090541 and 1U19NS104648, and the Simons Collaboration on the Global Brain (SCGB).

## Author Contributions

SAK performed the experiments, data analysis and conceptualization/modeling. ASC consulted on math and computational methods. SYT and DWT designed the experimental setups. SAK wrote the manuscript with input from DWT, CDB, and ASC. SAK, DWT, and CDB conceived the project.

## Data availability

The data that support the findings of this study are available from the corresponding author upon reasonable request.

## Code availability

Custom code (Matlab and C++) used to perform all analyses are available from the corresponding author upon reasonable request.

## Supplemental Information

**Table S1.** Number of imaging sessions and mice that contributed imaging sessions, for recordings of various cortical regions and layers.

|  | Layers 2/3 |  |  |  |  |  | Layer 5 |  |  |  |  |  |
| --- | --- | --- | --- | --- | --- | --- | --- | --- | --- | --- | --- | --- |
|  | V1 | AM | PM | MMA | MMP | RSC | V1 | AM | PM | MMA | MMP | RSC |
| # sessions | 9 | 18 | 11 | 15 | 8 | 30 | 8 | 12 | 5 | 8 | 7 | 12 |
| # mice imaged | 4 | 8 | 6 | 9 | 4 | 8 | 4 | 7 | 3 | 8 | 4 | 6 |

https://drive.google.com/open?id=1Oux0uhnq9e4UOX_NFMKUVqmEDzoyxe-g

**Movie S1.** Animation of Fig. 3 to show 3D layout of data points and alternative viewing perspectives.

---

**Figure S1.**
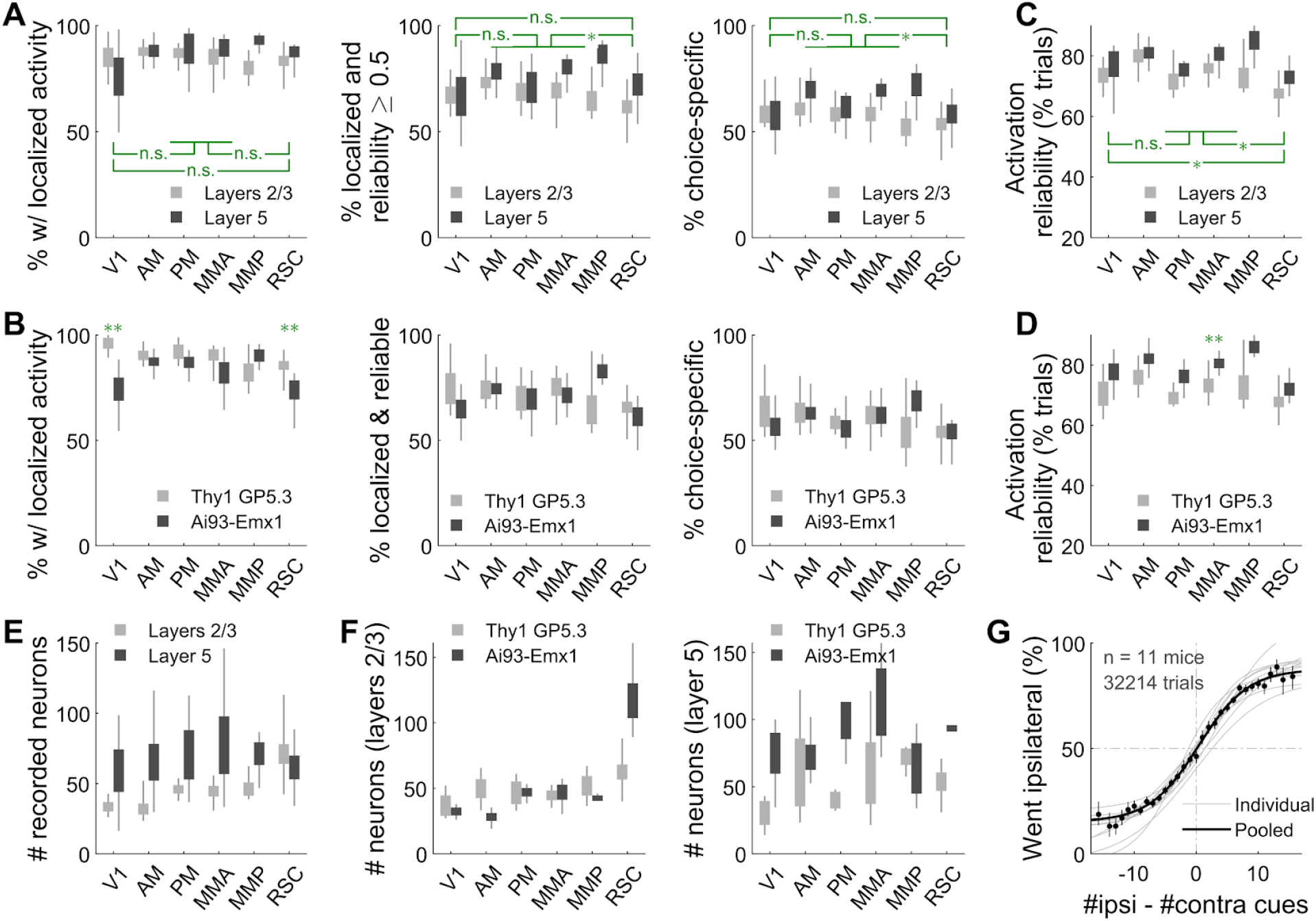
Statistics for choice-specific sequences vs. cortical regions and layers, and cross-strain comparisons. **(A)** Percents of neurons that had significant ridge-to-background excess vs. a permutation test (left plot), and additionally were active within their firing fields in ≥ 50% of their (preferred-choice, if any) trials (middle plot), and additionally had different activity levels in right- vs. left-choice trials (right plot). Error bars: std. dev. across imaging sessions. Rectangles: S.E.M. Stars: significant differences in means (Wilcoxon rank-sum test). **(B)** Like (A), but comparing two strains of mice. Data were pooled across layers. Double-stars indicate areas for which there was a significant difference in means (Wilcoxon rank-sum test). **(C)** Average reliability of choice-specific neurons in a given area/layer, defined as the fraction of trials in which the neuron was significantly active within its putative firing field. Error bars as in (A). **(D)** Like (C), but comparing two strains of mice. Data were pooled across layers. Double-stars indicate areas for which there was a significant difference in means (Wilcoxon rank-sum test). **(E)** Distribution of the numbers of simultaneously imaged neurons, for various brain regions and layers. Error bars: std. dev. across imaging sessions. Rectangles: S.E.M. **(F)** Like (E), but comparing datasets for two strains of mice. **(G)** Sigmoid curve fits to behavioral data for how frequently mice turned to the side ipsilateral to the recording hemisphere, for a given difference in total ipsilateral vs. total contralateral cue counts for the trial. Points: Percent of trials in which mice turned to the ipsilateral side, for trials with various tallies of ipsilateral vs. contralateral cue counts (data pooled across all mice). Error bars: 95% binomial C.I.

**Figure S2.**
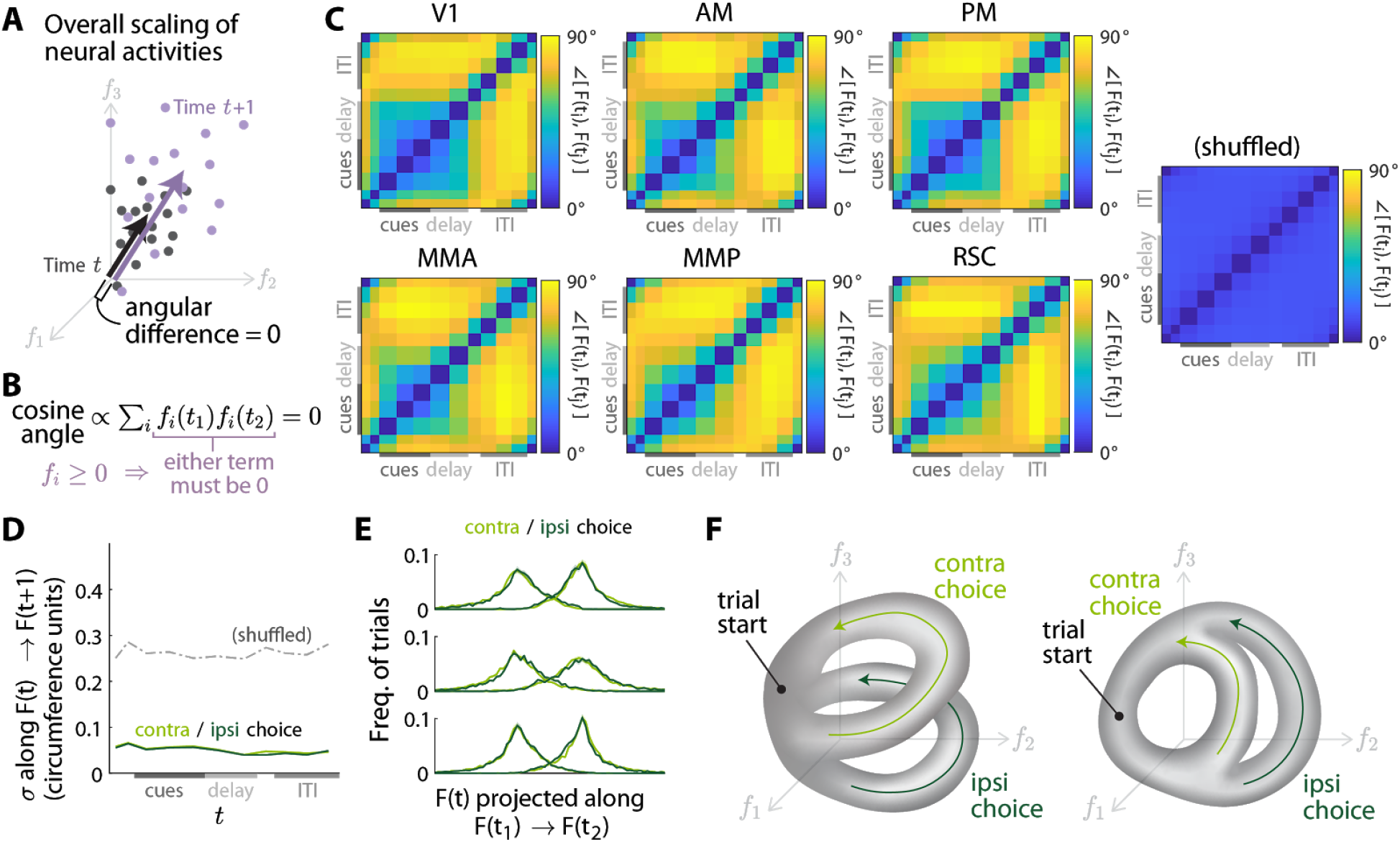
Neural manifold geometry metrics for all pairs of timepoints, and differences vs. navigational choice. **(A)** Illustration of a case where the neural states across trials at time *t* + 1 are an overall scaling of the neural states at time *t* (i.e. by the same scale factor for the activity of each neuron). This results in zero angular difference between the centers of the two point clouds (black vs. purple arrows), because the vector to the center of the *t* + 1 point cloud is also just a scaling of the vector to the center of the *t* point cloud. **(B)** Two vectors 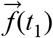 and 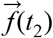 with nonnegative entries are orthogonal (zero cosine angle between vectors) if and only if they have *no* nonzero coordinates in common. This is because the cosine angle between these vectors is proportional to the sum over coordinates *i* of the product of their entries, *f_i_*(*t*_1_) *f_i_*(*t*_2_). If any *f_i_*(*t*_1_) *f_i_*(*t*_2_) ≠, then the nonnegativity of the vectors means that *f_i_*(*t*_1_) *f_i_*(*t*_2_) > 0. In this case, since all terms in the sum are nonnegative and cannot cancel each other out, the cosine angle is nonzero. **(C)** Angles between time-average neural state vectors *F*(*t*_1_) and *F*(*t*_2_), for all possible pairs of timepoints *t*_1_ and *t*_2_. Data were averaged across all imaging sessions for each stated cortical region. Shuffled: Pseudo-data with the neural state randomly cyclically permuted across imaging frames in the session. **(D-E)** Same as Fig. 4B,D, but plotted separately for trials where the mouse makes a choice to turn to the direction contra- or ipsi-lateral to the recording brain hemisphere. The same projection axes (cross-validated and defined as for Fig. 4B,D) were used for data of both trial types. **(F)** Illustration of how manifolds with global time order can have sub-structure that depends on trial-specific details, in this case the mouse’s navigational choice. In these two examples, trials of different choices approximately follow trajectories that bifurcate mid-trial, remain well-separated until the end of the trial, and then gradually converge in the ITI. See Fig. 3B-C for data examples.

**Figure S3.**
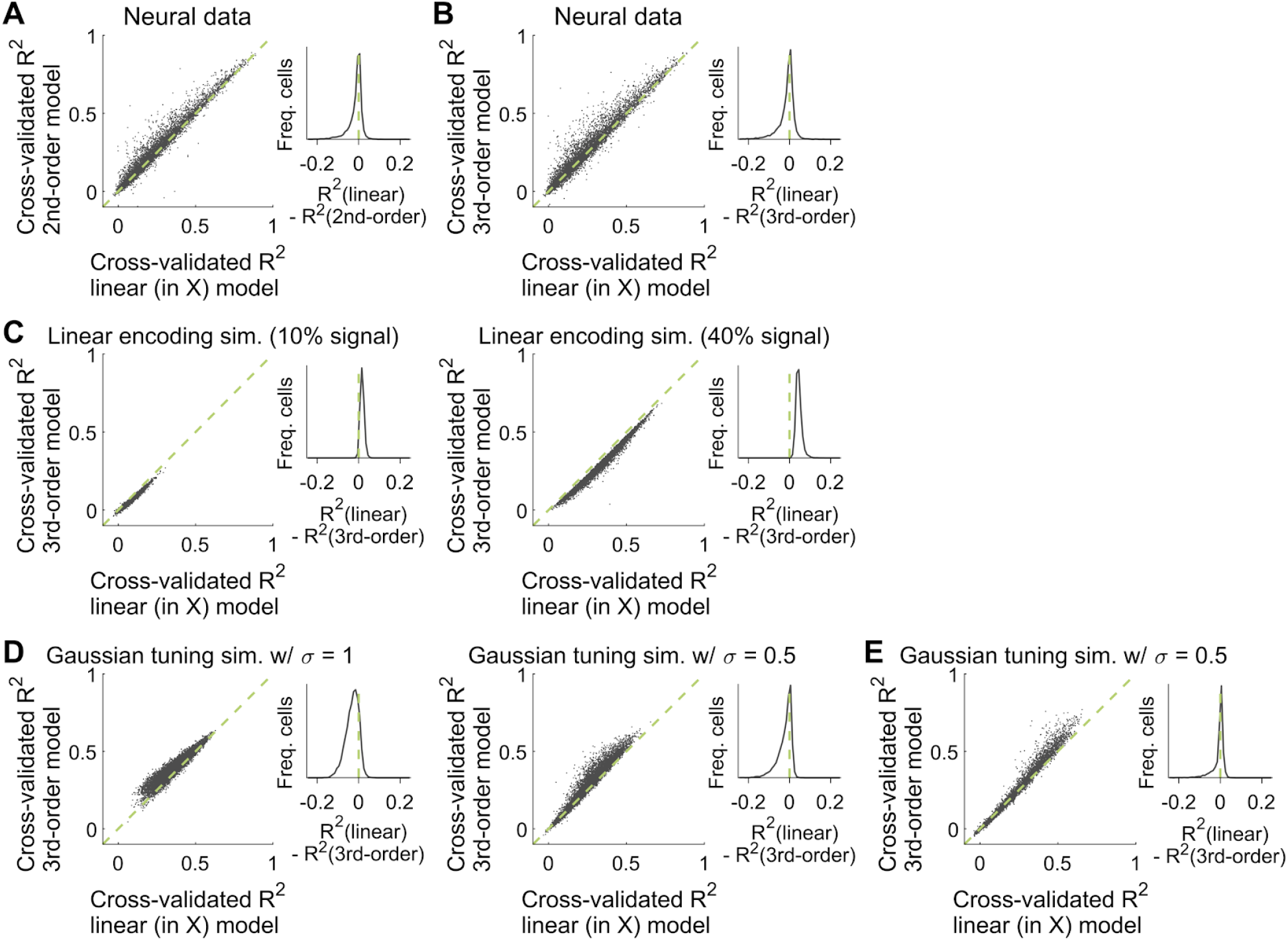
Nonlinear encoding models are only a little better at predicting neural activity compared to linear models. **(A)** Cross-validated variance explained for regression models with up to 2^nd^-order dependencies on task variables, vs. models with linear dependence on task variables (as used throughout the text). For each neuron and timepoint in the trial, ridge regression was performed to predict the activity of that neuron as a weighted sum of task variable values, and the nonlinear vs. linear models differ only in the set of considered regressors. For the linear model, the trial-by- *k* matrix for *k* = 13 task variables **X** was used as regressors, and for the 2^nd^-order model the regressors were 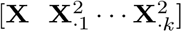 where **X**_.*i*_ is the *i* ^th^ column of the **X** matrix. Each point in this plot corresponds to the score for one neuron (including all imaging sessions), evaluated using all timepoints and trials. Inset: Distribution of differences in variances explained for the linear vs. 2^nd^-order regression model. **(B)** Same as (A) but for a 3^rd^-order regression model with regressors 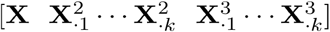. **(C)** Same as (B) but for simulated neurons that respond linearly to the task variables in the behavioral data. See Fig. S4A-B for how these simulations were constructed. Note that because the true response model is linear, cross-validation penalizes the nonlinear model for having excess parameters and thus the nonlinear model almost never outperforms the linear model. **(D-E)** Same as (B) but for simulated neurons that have multivariate gaussian tuning curves for their responses to the task variables in the behavioral data. See Fig. S4C,E for how the corresponding two types of gaussian-tuning simulations were constructed.

**Figure S4.**
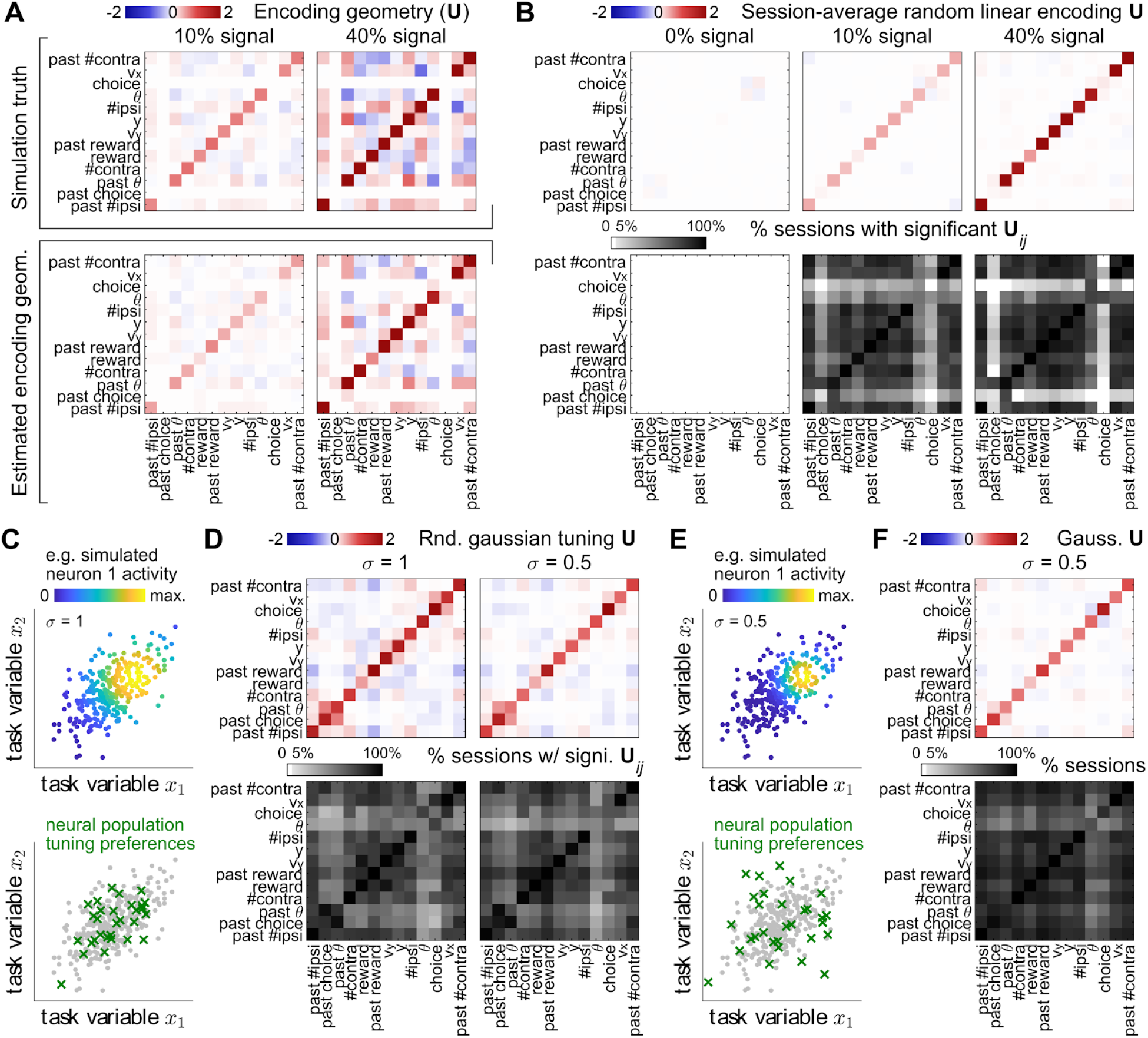
Encoding geometry in simulations of linear and nonlinear neural coding schemes for the correlated task variables in actual experiments. **(A)** Encoding geometry for a simulated neural population (63 neurons) that responded linearly to task variables, **F**(*t*) = **X**(*t*) **W**_sim_ + i.i.d.noise where **X**(*t*) is the task variable data vs. trials and time from an experimental session, and **W**_sim_ is a randomly generated (time-independent) encoding matrix for that session (228 trials). The neural population size and number of trials were selected to be around average for the data in our hands. Top row: the simulated encoding geometry 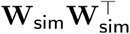. Each entry of **W**_sim_ was drawn from a random gaussian distribution centered around 0, except for choice and past-choice variables which were set to zero for all simulated neurons. Only the overall scale of **W**_sim_ differs between the two scenarios that have low (left column) vs. higher (right column) signal variance. Bottom row: time-average encoding geometry (see text and Methods) for the corresponding two simulated datasets in the top row. **(B)** Top row: Estimated encoding geometry averaged across time and simulated sessions, where neural data were simulated as described in (A) using the behavioral data in all 143 experimental sessions. Because the simulated encoding matrix **W**_sim_ was randomly drawn for each dataset, the session-average encoding geometry should theoretically tend towards a multiple of the identity matrix, except for choice and past-choice variables, which should be zero. Note that the reward variable is also partially zero in this time-average report, because reward was not defined until the end of the trial (and the corresponding encoding geometry entries were set to zero in undefined periods). Bottom row: Proportion of simulated sessions for which various encoding geometry entries were significantly different from chance (controlled for false discovery rate per simulation signal strength scenario). **(C)** Illustration for how simulations with gaussian-tuned neural responses were constructed. Top plot: example trials from a simulation with two correlated task variables *x*_1_ and *x*_2_, with points colored according to the activity level of one simulated neuron that has a 2D gaussian tuning curve for its responses to these task variables. This neuron has maximum response at (*x*_1_, *x*_2_) = (1, 0.5) and the tuning curve has width σ = 1 for both variables. Bottom plot: the same example simulated trials as in the top plot, but overlaid with the locations of maximum response for a population of simulated neurons (green crosses). These maximum-response locations were randomly drawn from a 2D gaussian distribution with covariance matrix being the covariance matrix of the simulated task variables, i.e. so that the neural tuning preferences tend to fall within parameter combinations that are actually sampled in the “behavior”. **(D)** Time- and session-average encoding geometry (top row) and proportion of sessions in which the encoding geometry was significantly different from chance (bottom row), same as in (B) but for simulations of gaussian-tuned neural responses as described in (C). The two columns correspond to **(E)** Illustration of an alternative construction of gaussian-tuned neural responses, where the maximum-response locations of the simulated neural population were randomly drawn from a 2D gaussian distribution with variance being twice the variance of the behavioral data (and zero off-diagonal terms in the covariance matrix). This means that neural tuning preferences for each task variable was independent of the tuning preferences for other task variables. **(F)** As in (D), but for the alternative construction of gaussian-tuned simulated responses as described in (E).

**Figure S5.**
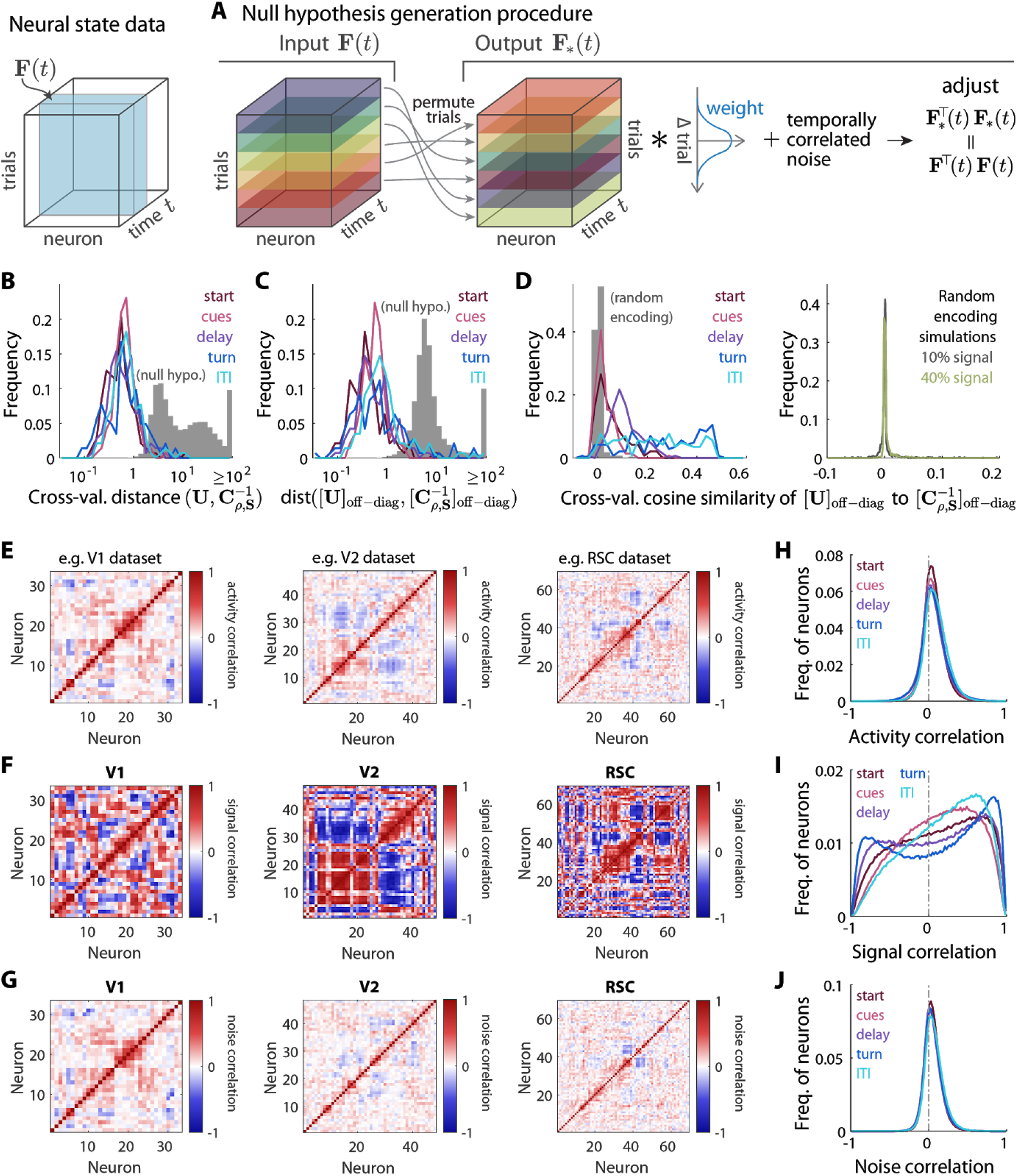
Additional metrics for comparing the encoding geometry to an inverse noisy task covariance, and correlations between pairs of neurons. **(A)** Sketch of the null hypothesis generation procedure (see Methods for details) starting from a given trial-by-neuron-by-time neural dataset {**F**(*t*)|*t* = 1, 2, …, 11}. First we permute each neuron-by-time “slab” of the neural data across trials, i.e. using the same trial permutations for each neuron and timepoint in the trial. This permuted pseudo-data **F**_∗_(*t*) is then row-convolved with a symmetric kernel designed to make the row-autocorrelation function of **F**_∗_(*t*) the same as that of **F**(*t*). A small amount of noise is then added with the same temporal correlations as the data, and on a logarithmic scale so as to preserve the bounded domain of the data. Lastly, a small number of iterative residual adjustments are made to constrain both the column-covariance structure and row-autocorrelation function of **F**_∗_(*t*) to be the same as **F**(*t*). **(B)** Analogous to Fig. 5F, but for the cross-validated Euclidean distance between the encoding geometry **U** and the inverse noisy task variable covariance matrix 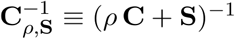. The Euclidean distance between matrices **A** and **B** is defined as 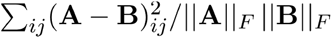, where the Frobenius norm of matrix **A** is 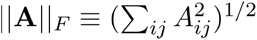. Colored lines: distribution across sessions for data in various phases of the trial (see caption of Fig. 5F). Gray histogram: distribution across sessions and phases, for the null hypothesis as explained in the text. **(C)** Same as (B), but where the Euclidean distance was computed using only off-diagonal entries of the two matrices: 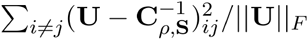. **(D)** Left plot: same as Fig. 5F, but for the cross-validated cosine similarity between **U** and 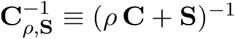 computed using only off-diagonal entries of the two matrices: 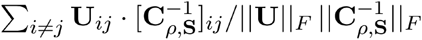. Colored lines: distribution across sessions for data in various phases of the trial (see caption of Fig. 5F). Gray histogram: distribution across sessions and phases for simulations where neurons randomly linearly encode task variables (with 10% signal variance, as explained in the Methods and caption of Fig. S4A). Right plot: distribution across sessions and phases for the random linear encoding simulations, with 10% vs. 40% signal variance (gray vs. green). **(E)** Pairwise correlation coefficients for the activities of neurons across trials, evaluated at a fixed timepoint at the end of the cue region. The various plots are for three example imaging sessions in the stated brain areas, selected to have the median number of neurons across all imaging sessions for that region. Neurons (i.e. the displayed order of rows and columns) were sorted using hierarchical clustering of this matrix. **(F)** Same format and order of neurons as (E), but for signal correlations between neurons, defined as correlations between the predicted activities of neurons according to the per-timepoint behavioral encoding models. **(G)** Same format and order of neurons as (E), but for estimated noise correlations between neurons, where “noise” was estimated as the residual activity of neurons after subtracting the behavior-based signal prediction in (F). **(H-J)** Distributions of correlation coefficients as in (E-G), for neurons pooled across all sessions but restricted to the stated time periods in the trial.

**Figure S6.**
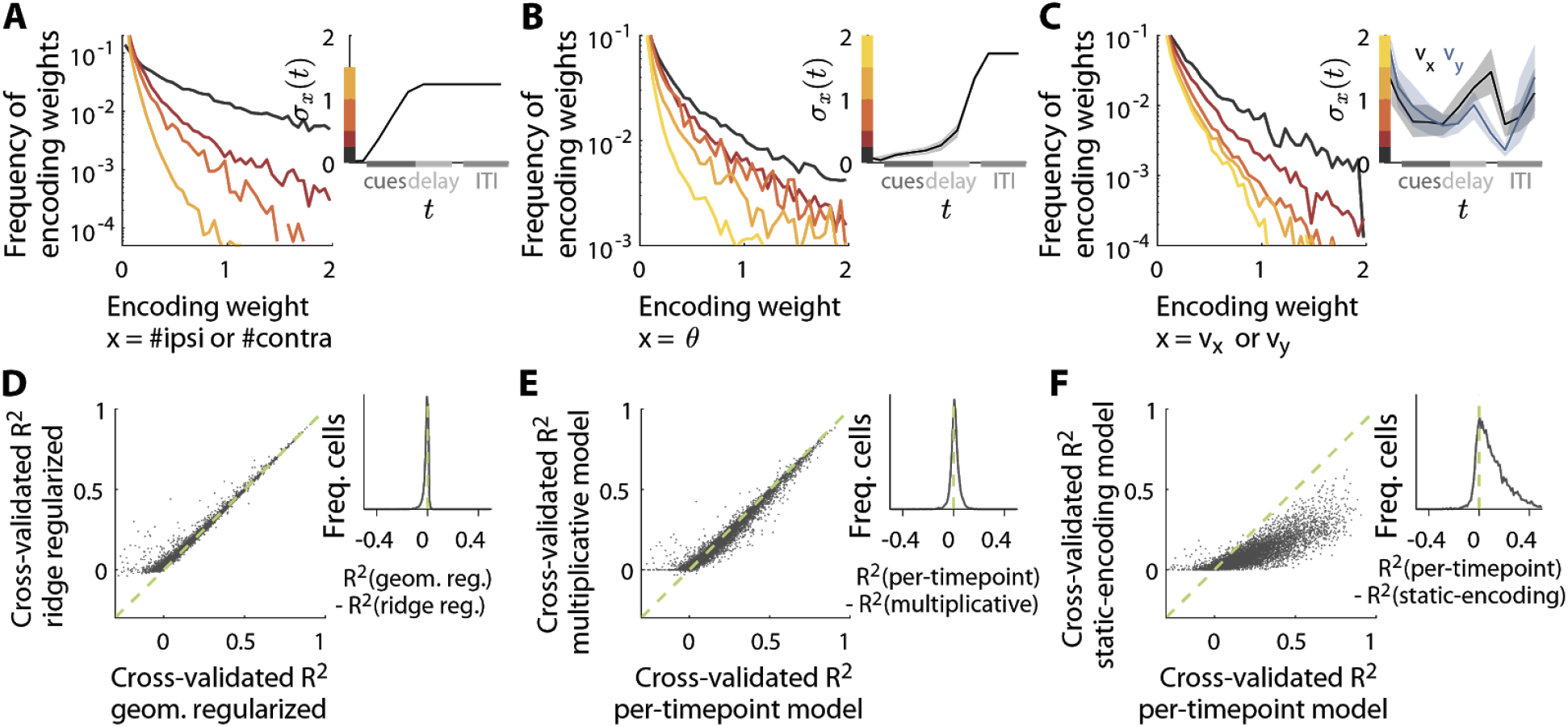
Time-dependence of task-variable encoding weights, and comparisons between various types of encoding models. **(A-C)** Dependence of encoding weights on the time-dependent scale of the respective task variables (insets). 11 encoding models were fit separately per timepoint for each neuron, which yields 11 encoding weights per neuron, for a given variable. The colored lines are distributions of these encoding weights restricted to timepoints in the trial where the time-dependent scale (standard deviation) of the task variable fell within the indicated bins (vertical colored bars in the inset plot). For comparability across neurons and task variables, encoding weights were expressed in units of 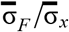, where 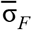 is the standard deviation of the activity level of a given neuron across all timepoints in the imaging session, and 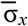 is the standard deviation of the task variable again across all timepoints. **(D)** Cross-validated variance explained for a model where ridge regression was used to predict the activity of each neuron (y-axis), vs. the neural-population-geometry regularized regression model used in Fig. 5-Fig. 7 (x-axis, see text and Methods). Each point in the left plot is for one neuron in an imaging session (including all sessions), where the variance explained is computed using all timepoints and trials in the data for that neuron. Inset: Distribution across neurons of differences in geometry-regularized vs. ridge-regularized variances explained, i.e. x minus y coordinates of points in the left plot. **(E-F)** Same as Fig. 7A-B, but using the geometry-regularized regression models (Methods) instead of ridge regression.

**Figure S7.**
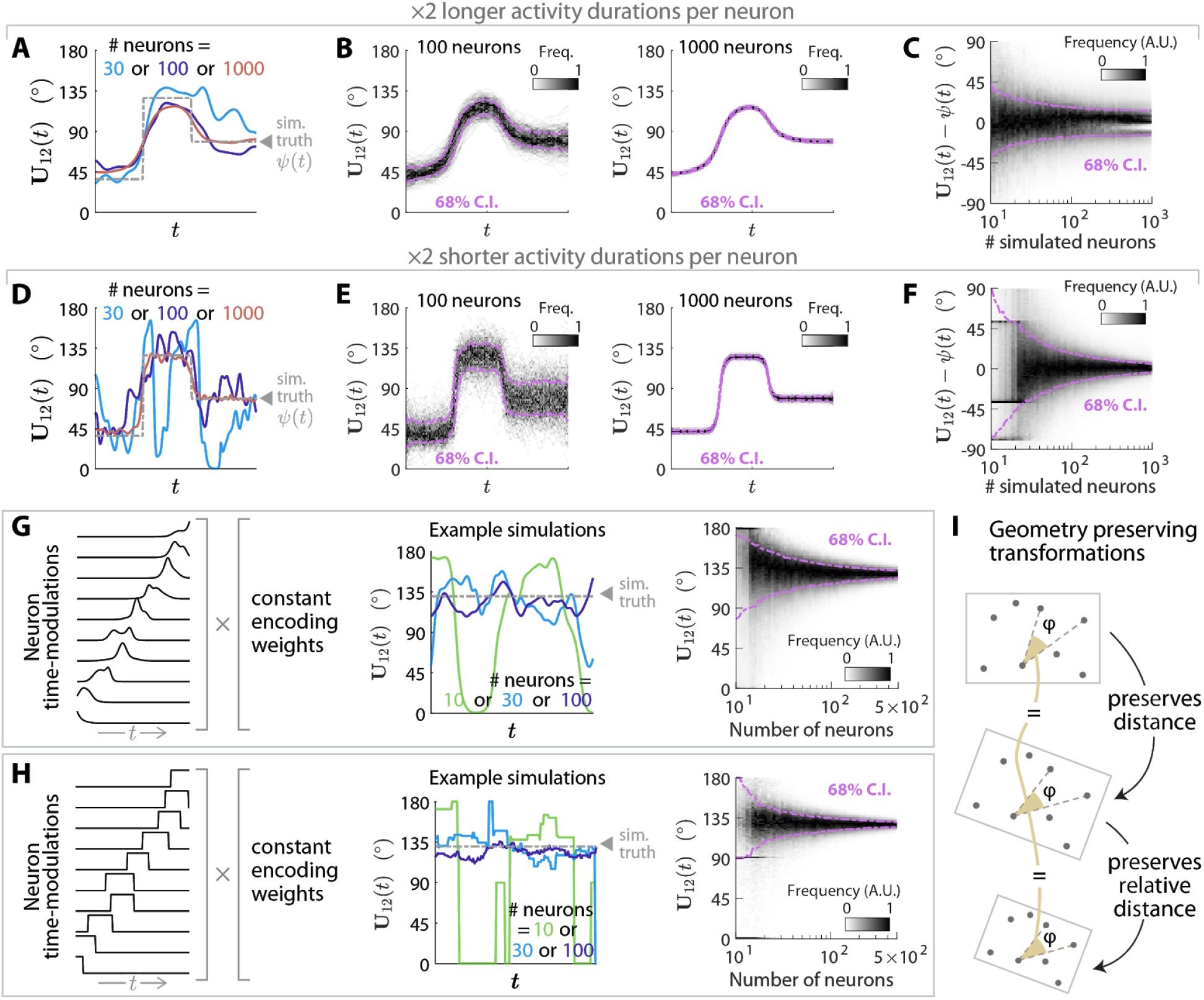
Convergence of simulated multiplicative sequential encoding geometries to different forms of design truth, as a function of neural population size. **(A)** Same as Fig. 7E, except that ψ(*t*) was designed to be a function with two abrupt steps as shown in the plot (dash-dotted gray line), and the generation parameters of *g_i_*(*t*) (Methods) were scaled by a factor of 2 to create neural activities that were longer in duration than the simulations discussed in the text. **(B)** Distribution across simulation experiments of the time-dependent angle between encoding directions, for the simulation scenario in (A). The left and right plots correspond to simulations with neural population sizes of 100 and 1000 respectively. **(C)** Distribution across simulation experiments and time of differences between ψ(*t*) vs. the angle between encoding directions, for the simulation scenario in (A) and as a function of the simulated neural population size. **(D-F)** As in (A-C), but the generation parameters of *g_i_*(*t*) were scaled by a factor of 0.5 instead to create neural activities that were shorter in duration than the simulations discussed in the text. **(G-H)** Same as Fig. 7E-F, but where ψ(*t*) was designed to be a constant 130°. **(I)** Illustration of two transformations that preserve distances (rotation) and relative distances (uniform scaling) between points. Also shown is an angle between two vectors (dashed lines), which remains the same if distances are preserved, and also if relative distances are preserved, as angles do not depend on lengths of vectors. The dot product between two vectors depends on both the cosine angle between the vectors as well as their magnitudes, and thus do change under uniform scaling. However if all coordinates are uniformly scaled by a factor of *s*, then the dot products between all pairs of vectors are scaled by same factor *s*^2^, and thus the relative magnitudes of dot products are unchanged.

**Figure S8.**
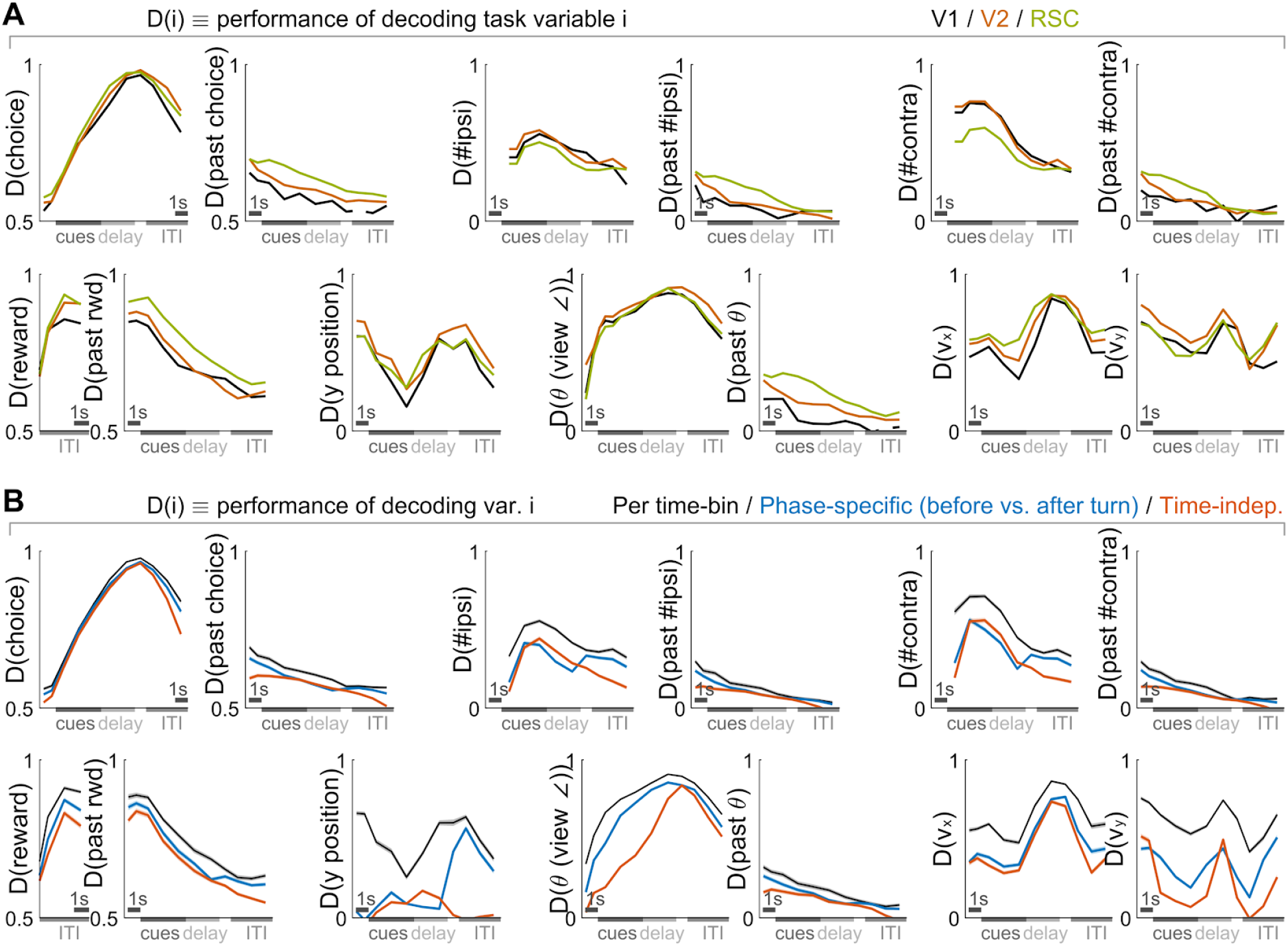
Cross-validated decoding accuracies for different variables and decoding methods. **(A)** Cross-validated performance for decoding thirteen task variables (individual plots), vs. time in the trial. For each variable, a different linear decoder (L2-regularized support vector machine) was trained per timepoint. For categorical variables, the performance measure is the cross-validated classification accuracy, where chance level is 0.5. For continuous variables, the performance measure is the cross-validated correlation between decoded and actual variable values, where chance level is 0. Performance was averaged across imaging sessions for a given area (lines), with points blanked out if < 5% of sessions had significant decoding performance vs. a permutation test (*p* ≤ 0.036 post correction for multiple comparisons). **(B)** Decoding performance for per-timepoint decoders in (A), vs. phase-specific decoders (blue lines), and decoders trained using data from all timepoints (red lines). The phase-specific decoders were trained using data in 2 separate phases of the trial. The first phase included all timepoints from the start of the trial to the end of the delay region, and these timepoints were treated like additional trials when training decoders. The second phase included the remaining timepoints from the start of the turn region to the end of the ITI. Decoding performance were evaluated at each timepoint, as in (A). Lines: mean across imaging sessions (i.e. including all cortical regions in (A)). Band: S.E.M.

